# Longitudinal profiling of the microbiome at four body sites reveals core stability and individualized dynamics during health and disease

**DOI:** 10.1101/2024.02.01.577565

**Authors:** Xin Zhou, Xiaotao Shen, Jethro S. Johnson, Daniel J. Spakowicz, Melissa Agnello, Wenyu Zhou, Monica Avina, Alexander Honkala, Faye Chleilat, Shirley Jingyi Chen, Kexin Cha, Shana Leopold, Chenchen Zhu, Lei Chen, Lin Lyu, Daniel Hornburg, Si Wu, Xinyue Zhang, Chao Jiang, Liuyiqi Jiang, Lihua Jiang, Ruiqi Jian, Andrew W. Brooks, Meng Wang, Kévin Contrepois, Peng Gao, Sophia Miryam Schüssler-Fiorenza Rose, Thi Dong Binh Tran, Hoan Nguyen, Alessandra Celli, Bo-Young Hong, Eddy J. Bautista, Yair Dorsett, Paula Kavathas, Yanjiao Zhou, Erica Sodergren, George M. Weinstock, Michael P. Snyder

## Abstract

To understand dynamic interplay between the human microbiome and host during health and disease, we analyzed the microbial composition, temporal dynamics, and associations with host multi-omics, immune and clinical markers of microbiomes from four body sites in 86 participants over six years. We found that microbiome stability and individuality are body-site-specific and heavily influenced by the host. The stool and oral microbiome were more stable than the skin and nasal microbiomes, possibly due to their interaction with the host and environment. Also, we identified individual-specific and commonly shared bacterial taxa, with individualized taxa showing greater stability. Interestingly, microbiome dynamics correlated across body sites, suggesting systemic coordination influenced by host-microbial-environment interactions. Notably, insulin-resistant individuals showed altered microbial stability and associations between microbiome, molecular markers, and clinical features, suggesting their disrupted interaction in metabolic disease. Our study offers comprehensive views of multi-site microbial dynamics and their relationship with host health and disease.

**Study Highlights:**

1. The stability of the human microbiome varies among individuals and body sites.
2. Highly individualized microbial genera are more stable over time.
3. At each of the four body sites, systematic interactions between the environment, the host and bacteria can be detected.
4. Individuals with insulin resistance have lower microbiome stability, a more diversified skin microbiome, and significantly altered host-microbiome interactions.

## Introduction

The human microbiome comprises highly dynamic microbial communities inhabiting various body sites ^1–5^, engaging in intricate host-microbial interactions that display territory-specific complexity^6–10^. Advancements in multi-omics profiling technologies have catalyzed the elucidation of the molecular mechanisms underlying microbial ecology and their interactions with host systems, unveiling the critical roles of the microbiome in normal physiological processes such as aging^11–13^ as well as diseases including inflammatory bowel disease (IBD)^14–16^, cardiovascular disease^17–19^, and type 2 diabetes mellitus (T2DM)^20–23^.

The etiology and pathogenesis of insulin resistance and T2DM have been closely linked to the human microbiome^24–27^. Patients with impaired insulin and glucose homeostasis exhibit microbiome composition alterations in the gut^21,23,25–29^, skin^30–33^, and other body sites^34–40^, reflecting an ecological dysbiosis characterized by altered microbial alpha diversity^25,41^, decreased compositional stability^42,43^, and greater inter-individual variability^25^. Compromised integrity of mucosal and skin barriers, often associated with insulin resistance, may potentiate microbial translocation, thereby exacerbating systemic inflammation^44–47^. Although human microbiome studies are often, by necessity, observational, a causal relationship between microbiome dysbiosis and impaired glucose/insulin homeostasis has been demonstrated in patients and animal models and through human microbiome manipulation^41,48,49^.

While prior studies on the microbiome and glucose homeostasis have been informative, they exhibit certain limitations. Firstly, these studies^21,50,51^ often lack longitudinal microbiome sampling, essential for capturing individuality and stability features, thus limiting fundamental insights into host-microbe interactions^52–54^. Secondly, they primarily focus on the microbiome from a single habitat^4,50,55–58^, overlooking the importance of simultaneous multi-region sampling for assessing microbiome site-specific dynamics, and their interplay across various host microenvironments^9,59–62^. Lastly, these studies^8,18,21,63–66^ do not concurrently measure multiple host clinical and molecular phenotypes, impeding the exploration of the molecular relationships underpinning health and disease-related host-microbiome interactions^67,68^.

Collaborative initiatives like the Integrative Human Microbiome Project (iHMP, https://www.hmpdacc.org/ihmp/) and Integrative Personal Omics Profiling (iPOP, https://med.stanford.edu/ipop.html) offer avenues to surmount previous studies’ limitations by investigating well-characterized human longitudinal cohorts^69–71^. In this study, we examined the relationships between multi-site microbiomes and host health in the context of prediabetes, through characterizing the microbiome collected from four body sites in 86 adults for over six years and examining their associations with host omics and clinical characteristics. We described unique longitudinal trajectories for microbiomes across different body sites, demonstrating their responsiveness to both host-specific and environmental factors. Noteworthy is the higher stability of personalized microbiomes compared to shared ones, illustrating the robustness of individual microbial stability. Even so, we found the microbial equilibrium can be disrupted by events such as viral infections, potentially contributing to chronic dysbiosis associated with metabolic disorders. Moreover, we identified extensive correlations between the microbiomes at each body site and clinical indicators, including cytokines and insulin resistance, underscoring their impact on both immune response and metabolic wellbeing.

## Results

### Description of the study design

We analyzed the host microbiome at four distinct body sites (stool, skin, tongue/oral cavity, and nasal) from a cohort of 86 participants, sampled quarterly for up to six years (mean 1,126.6 ± 455.8 days). The cohort comprised 41 males and 45 females, aged between 29 and 75 years old (mean 55 ± 9.8 years old), with BMIs ranging from 19.1 to 40.8 kg/m^2 (28.31 ± 4.44 kg/m^2) (**Table S1**). The gut microbiome was evaluated using stool samples, whereas skin, oral, and nasal microbiomes were sampled using swabs of the retroauricular crease, oral cavity/tongue, and anterior nares, respectively (**Fig. 1a**, see Methods). Sampling occurred quarterly when participants were healthy, with an additional 3-7 samples collected within five weeks (12% of the total) during periods of stress, such as respiratory illness, vaccination, or antibiotic use. The 16S ribosomal RNA gene sequencing method employed in this study targeted a variable region (see Methods) to facilitate the detection of amplicon sequence variants (ASVs)^72^, enabling the identification and differentiation of most bacterial taxa at the genus and species levels^73,74^.

**Figure 1.**
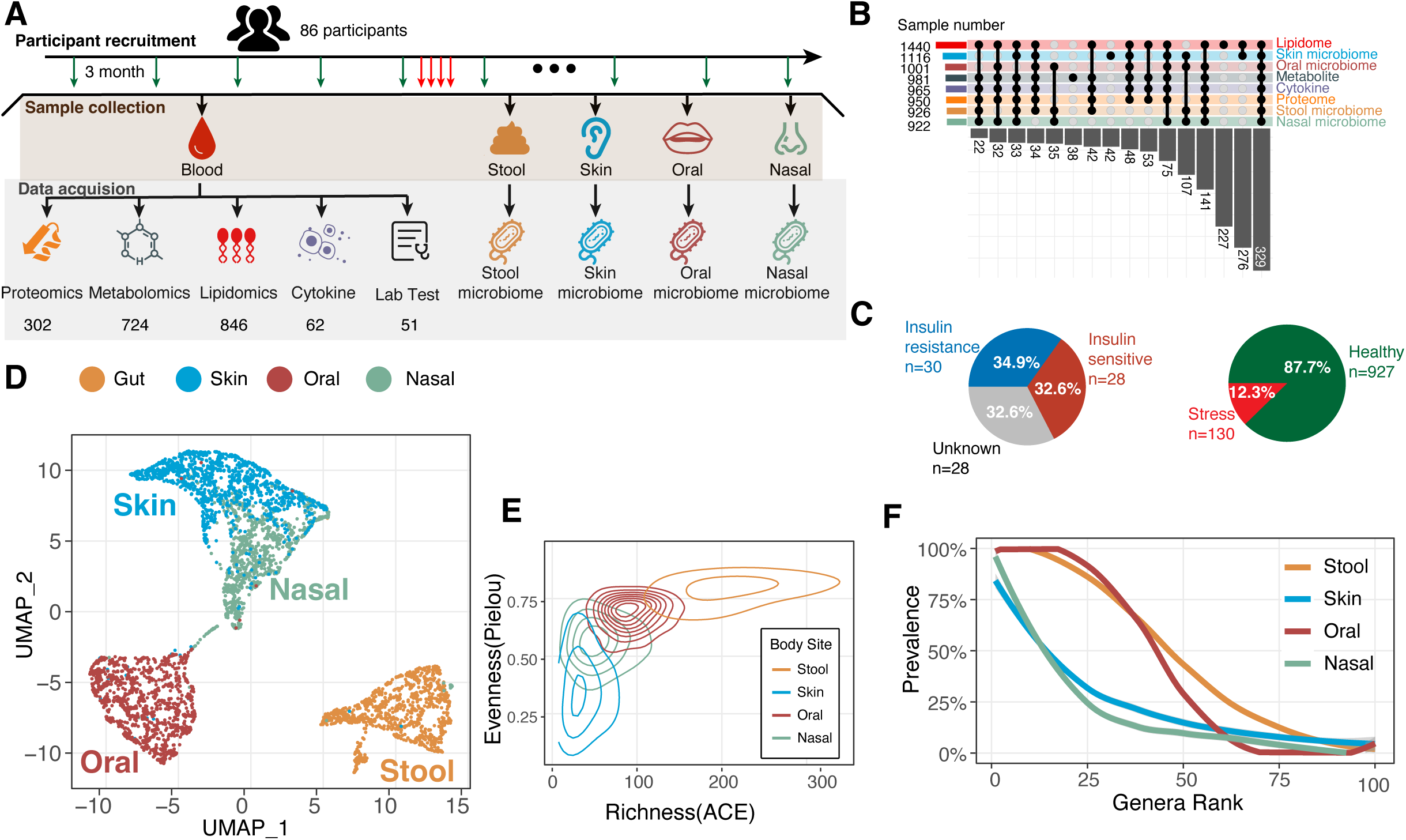
Longitudinal Profiles of the Microbiome at Four Body Sites. A. Study Design. The study design involved the examination of participants every three months, with additional visits and sample collections from participants who reported experiencing stress. During each visit, samples were collected, including blood, oral/nasal/skin swabs, and stool samples. These samples were used to generate multi-omics data encompassing proteomics, metabolomics, lipidomics, and cytokines (including chemokines and growth factors). Additionally, blood samples were sent for clinical laboratory testing. B. Data Overlap Between Omics Types: This figure depicts the overlap of sample numbers among different omics types. Each omics type is represented in a specific color and marked on the right Y-axis, with the number of samples from each type indicated on the left Y axis. The size of the interaction (overlap) is shown on the X axis where dots on the graph are intersected by a vertical line. C. Proportion of Stress and Healthy Samples: This section displays the proportion of samples collected during periods of stress (including infection, immunization, antibiotic use, etc.) and healthy periods. It also indicates the proportion of samples collected from insulin-sensitive and insulin-resistant participants. D. UMAP of Body Site Microbiomes: A Uniform Manifold Approximation and Projection (UMAP) plot illustrates the microbiomes of the four body sites. Each dot on the plot corresponds to a single microbiome sample from a specific body site, denoted by its color. E. Density Distribution of Microbiome Samples: This segment shows the distribution of microbiome samples from the four body sites, using the observed species richness (ACE metric) on the x-axis and evenness (Pileous metric) on the y-axis. F. Rank Prevalence Curve of Genera: The 100 microbiome genera with the highest longitudinal prevalence at each body site are displayed in this graph. They are sorted from high to low prevalence on the x-axis, and genera from the same body site are connected by a colored curve.

A unique feature of this cohort is the extensive phenotyping of participants at each timepoint using multi-omics analyses and clinical markers (**Fig. 1b**; see Methods). Multi-omics analyses employed in this study comprised untargeted proteomics (302 proteins), untargeted metabolomics (724 annotated metabolic features), targeted lipidomics (846 annotated lipids), and 62 targeted cytokine and growth factor measurements. Additionally, 51 clinical markers, including C-reactive protein (CRP), fasting glucose (FG), hemoglobin A1C (HbA1C), low-density lipoprotein (LDL), and high-density lipoprotein (HDL), were assessed from plasma samples at each timepoint. (**Fig. 1a**). Glucose control assessments, comprising an annual oral glucose tolerance test for all participants and a gold-standard steady-state plasma glucose (SSPG) measurement^75^ for 58 individuals, classified 28 as insulin-sensitive (IS) and 30 as insulin-resistant (IR) in this cohort. (**Fig 1c**) Overall, we analyzed a total of 3,058 visits, 5,432 biological samples (1,467 plasma samples, 926 stool samples, 1,116 skin samples, 1,001 oral samples, and 922 nasal samples), as well as clinical tests, generating a total of 118,124,374 measurements. The microbial and other data can be found on our data portal: https://portal.hmpdacc.org/.

### Microbial Demographics and Personalization Across Body Sites: Unraveling the Impact of Diet and Environment

We initially examined the overall demographic composition of the microbiome at each of the four body sites by utilizing dimension reduction such as Uniform Manifold Approximation and Projection (UMAP) (**Fig. 1d**) and Principal Coordinates Analysis (PCoA) (**Fig. S1**). In accordance with previous studies^7,62,76–78^, we observed a distinct separation between body sites, including a clear separation of the skin and nasal samples, highlighting the pronounced territory specificity of each microbiome^6,7,79^.

Micro-biotypes, like enterotypes in the stool microbiome^50,77,78^, are present in all body sites, with their community structure predominantly influenced by specific taxa. The stool microbiome primarily exhibited a gradient of abundance distributions between *Bacteroidetes* and *Firmicutes*, except for a few samples with high *Prevotella*. The recently identified core genus *Phocaeicola*^80^ had minimal impact on the overall *Bacteroidetes/Firmicutes* gradient, but samples with high *Phocaeicola* and *Bacteroides* formed distinct clusters. (**Fig. S1**) The oral microbiome from our cohort was primarily composed of *Prevotella*, *Streptococcus*, *Veillonella*, *Haemophilus*, *Neisseria*, and *Leptotrichia*, as previously described^81–83^. The skin and nasal microbiome samples exhibited the greatest similarity and jointly displayed a triangular distribution, primarily driven by three dominant genera: *Cutibacterium*, *Corynebacterium*, and *Staphylococcus* (**Fig. 1d, Fig. S1**) ^4,8,62,79^. The distribution of these microbial genera, consistent with other cohorts ^8,59,84^, remains largely unaffected by our study design that includes participants with diverse insulin sensitivities (**Fig. S2**). Our study, however, extends these observations by longitudinally comparing inter- and intra-individual covariance across all four body sites within the same cohort.

Intraclass (intra-individual) correlation coefficient (ICC) analysis confirmed that microbial personalization is more pronounced at the ASV level than at broader taxonomic resolutions (**Fig. S3**), highlighting stronger individualization with finer taxonomy. Notably, despite similar ecological characteristics and dominant bacterial constituents, the nasal microbiome manifested greater personalization than the skin microbiome (**Fig. S4A**), possibly because nasal microbiome dynamics are more host dependent^79^.

While environmental factors like season and diet can influence the human microbiome, our results suggest that the impact of seasonality on the gut microbiome, despite mixed results in previous studies^50,85–87^, is relatively modest compared to individual influences across the four body sites. Likely due to its direct external exposure, the skin microbiome exhibited the most distinct seasonal effect. This was followed by the oral microbiome, which could be shaped by seasonal dietary changes^51,88,89^, as it explained more dietary variance compared to other body site microbiomes (**Fig S5a**). We also observed a significant fluctuation of the skin and oral microbiome evenness (relative representation of species, measured by Pielou’s evenness index) that the two unique communities both become more even during summer with an increased number of different microbes (richness, measured by Abundance-based Coverage Estimator, ACE), (**Fig. S4b**) possibly triggered by environmental changes such as temperature and humidity^85,90^. We further examined the relationship between the environmental exposure and microbiome composition among the two participants with accessible environmental and chemical exposure data^90^. We found stronger exposome-skin microbiome covariance than in other body sites (**Fig. S5b**) with noticeable environmental impacts on internal sites like oral and stool microbiomes^91^, supporting our earlier single-individual observation^92^. By modeling^85^ the microbial dynamics across seasons, we found more pronounced alterations in skin and nasal microbiomes than in stool and oral microbiomes. (**Table S2**) These findings not only extend our knowledge of gut microbiome specificity and individuality^56,93^ to other body sites but also broaden our understanding of environmental influences on various microbiomes, with diet shaping oral microbiomes and environmental exposure impacting skin and nasal microbiomes.

### Microbiome from Distinct Body Sites are Ecologically Unique and Altered in Insulin Resistance

Beyond the influence of personal and environmental factors, we observed distinct ecological attributes among the microbiome at the four body sites (**Fig. 1e**). The stool microbiome exhibited the greatest richness and evenness, underscoring its essential complexity and functional significance. In contrast, the skin microbiome displayed a more skewed population due to its lower evenness compared to the nasal microbiome, despite their similar richness distributions. These ecological features, which often shift with disease progression, aligned with previous findings of IR-related gut dysbiosis^25,70^, characterized by a significant decrease in stool microbiome’s alpha diversity (t = 2.8462, p-value = 0.0067). Furthermore, shifts in the richness-evenness scatter plots for all four body sites (**Fig. S6a**) suggested systemic dysbiosis in IR individuals, highlighted by significantly higher skin microbiome richness (t = -2.9102, p-value = 0.0057) (**Fig. S6b**) and evenness (t = -2.4393, p-value = 0.019) (**Fig. S6c**). These shifts indicate that IR-associated dysbiosis extends beyond the gut microbiome.

Importantly, extending longitudinal observations from single body sites^56,94–96^, our analysis defines a “core microbiome” (see Methods) as microbes consistently present over time, representing potentially indispensable^97,98^ genera at each of the body sites. Interestingly, we found that stool and oral microbiomes maintain more than 25 highly prevalent core genera, whereas nasal and skin microbiomes only had three (**Fig. 1f**). Importantly, some highly prevalent (i.e., frequency within an individual) genera have low relative abundance, (i.e., *Coprococcus* Mean Prevalence = 80.75%; Mean Relative Abundance = 0.544%), highlighting the potential significance of low-abundance strains universally (**Fig. S7**). Intriguingly, the observed richness of core genera in stool and oral microbiome is negatively associated with steady-state plasma glucose (SSPG) (Spearman Rho = -0.52, p-value = 0.00047) and BMI (Spearman Rho = -0.40, p-value = 0.005), respectively (**Fig. S8**), indicating that IR and obesity may be associated with a loss of core microbiomes at these sites.

Given this novel link, we further explored differences in the microbiome ecology between insulin resistant (IR) and insulin sensitive (IS) individuals. IR subjects had a significantly higher number of skin core genera (t = -2.5856, p-value = 0.014) and a lower number of stool core genera (t = 2.9659, p-value = 0.0051) compared to IS individuals (**Fig. S9a**). Notably, several butyrate-producing bacteria (i.e., *Coprococcus*, *Parasutterella*, and *Butyricicoccus*) were enriched in IS stool core microbiomes, whereas diabetes-related opportunistic skin pathogens (i.e., *Finegoldia*^99–101^ and *Acinetobacter*^102–105^) were enriched in the skin core microbiome of IR individuals (**Table S3**). In addition, we found a clear divergence of the rank prevalence curves in stool and skin microbiome (**Fig. S9b**), demonstrating a global microbial prevalence shift in IR individuals at these two sites.

Additionally, several taxa differed significantly between IR and IS individuals in relative abundance. The stool microbiome of IR individuals showed an increase of genus *Phocaeicola* (LEfSe_effect size: 0.03; BH-adjusted p-value = 0.017) and a reduction of the genus Unclassified *Ruminococcaceae* (LEfSe_effect size: 0.017; BH-adjusted p-value = 0.0039), whereas the skin microbiome exhibited a decrease in the genus *Cutibacterium* (LEfSe_effect size: 0.069; BH-adjusted p-value = 0.007) and an increase of the genus *Peptoniphilus* (LEfSe_effect size: 0.0076; BH-adjusted p-value = 0.0022), which has been previously associated with diabetes-related skin dysbiosis^100,106,107^ and necrotizing infections^108^ (**Fig. S10; Table S4**). No such differences were observed in oral and nasal microbiomes via this method. These findings reinforced the hypothesis that the skin and stool microbiome’s stability is altered in IR individuals. Overall, our results reveal that IR participants exhibit a stool microbiome with reduced richness and butyrate-producing bacteria, and a skin microbiome more susceptible to opportunistic pathogens.

### Distinct Taxon-Specific Stability and Individuality in Microbiomes Across Body Sites: A Potential Pathway for Personalized Interventions

We next examined microbiome stability at the genus level across multiple body sites over time, hypothesizing stability is taxon- and site-specific. We defined a novel metric “Degree of Microbial Individuality” (DMI, see Methods) for each genus, which measures similarity within an individual relative to the population; a high DMI means the stability of the different microbes is highly specific to an individual. We also calculated a “Family Score” (FS, see Methods) to evaluate microbial dissimilarity within households. Expanding from observations of the stool microbiome^54,70,109^, we noted that skin, oral and nasal microbiome also demonstrated significantly less intra-individual and within-family variation compared to the variation observed between different individuals (**Fig. 2a, Table S5**). Within each individual, we observed the highest rate of complete turnover of ASV within a genus over time in the nasal microbiome (35.5% of cases), compared to stool (24.2%), skin (24.4%), and oral sites (2.7%). This finding underscores significant intra-individual ecological dynamics at the sub-genus level within microbiomes, despite overall community stability.

**Figure 2.**
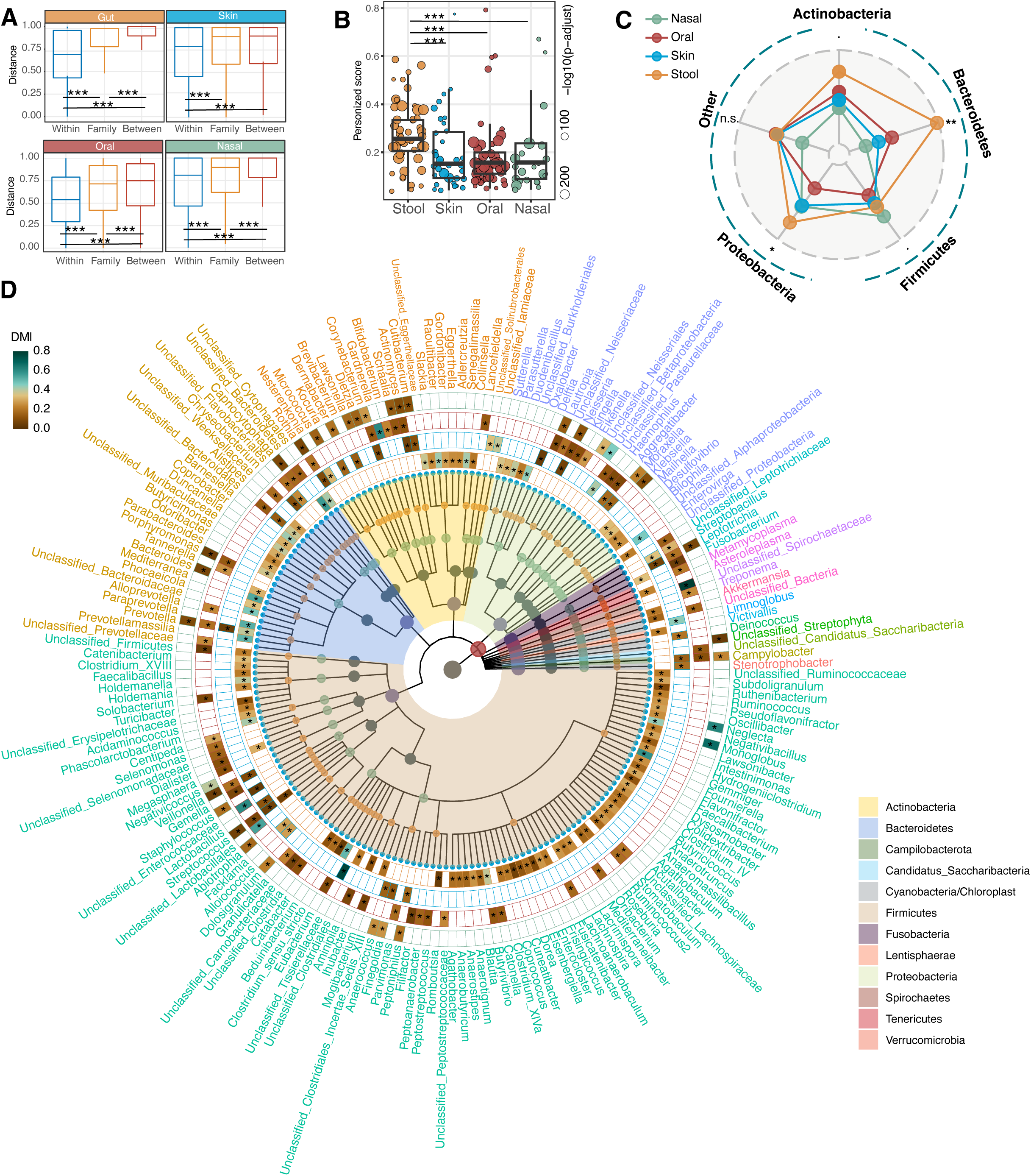
The Individuality of the Microbiome Differs Significantly Across Genera and Body Sites. A. Bray Curtis Dissimilarity Among Sample Groups: The Bray Curtis (BC) dissimilarity between sample pairs from different groups: “Within” (sample pairs from the same participants), “Family” (sample pairs from different participants living in the same household), and “Between” (sample pairs from different participants). The significance of these differences was determined using the Two-Sided Wilcoxon Rank test. The adjusted p-values are depicted with asterisks (*: p < 0.05; **: p < 0.01; ***: p < 0.001). B. Degree of Microbial Individuality (DMI) Scores: The DMI scores at different body sites are depicted in this section. Only genera with significant differences between intra- and inter-individual BC distances are included. The size of the dot indicates the log-10 transformed, BH-adjusted p-value when comparing the BC intra-individual and BC inter-individual distances. The significance of the difference between body sites were determined using the Two-Sided Wilcoxon Rank test. The adjusted p-values are depicted with asterisks (*: p < 0.05; **: p < 0.01; ***: p < 0.001). C. Radar Plot of Average DMI According to Body Sites and Phyla: This radar plot shows the average DMI for different body sites and phyla, with the DMI range from 0 (center) to 0.4 (edge of circle). A Kruskal-Wallis rank sum test was performed to for each of the following phyla: *Actinobacteria*: KW chi-sq = 9.512, BH adjusted p-value = 0.039; *Bacteroidetes*: KW chi-sq = 27.801, BH adjusted p-value = 0.00002; *Firmicutes*: KW chi-sq = 14.157, BH adjusted p-value = 0.0081); *Proteobacteria*: KW chi-sq = 9.5081, BH adjusted p-value = 0.039; Other genera not belonging to the aforementioned four phyla: KW chi-sq = 2.522, BH adjusted p-value = 0.47. The adjusted p-values are depicted with asterisks (*: p < 0.05; **: p < 0.01; ***: p < 0.001). D. Cladogram of DMI Across All Genera: The cladogram within the circles displays the phylogenetic relationships among all nodes (genera). The heatmap circles represent the color-coded DMI score, with four circles indicating the DMI of the genus for the stool, skin, oral, and nasal microbiome, from inner to outer circle. Asterisks on the heatmap for a particular species and body site signify a significant difference between intra- and inter-individual BC distances, as determined by the permutation test.

The DMI, irrespective of relative abundance, were predominantly high in the stool microbiome (**Fig. 2b**), particularly within the *Bacteroidetes* phylum (**Fig. 2c, Table S6**), possibly due to its pronounced adaptive evolution^110,111^ and substantial colonization resistance^112^. Furthermore, the stool microbiome had the lowest FS, indicating high genus dissimilarity within households, whereas oral and nasal microbiomes shared greater similarity (**Fig. S11**), likely due to common living environments^56,113^ or direct microbiome exchanges^114^.

Our data revealed substantial DMI variance across body sites, potentially attributable to inherent niche-specific taxonomic complexities. (**Fig. 2d, Table S7**). For example, *Corynebacterium* and *Bacteroides* showed the highest DMI in nasal and stool microbiomes, respectively. These results suggest that specific microbial taxa, adapting to their respective niches, may exhibit enhanced individualization. Therefore, the microbial community at each body site is largely shaped by these niche-specific interactions. In contrast, environmental bacteria such as *Klebsiella*^115^ and *Haemophilus*^116^ displayed uniformly low DMI across all examined body sites. This evidence further suggests that external environmental factors might exert a relatively weaker influence on the individuality of the native host microbiome. Interestingly, after adjusting individuals’ collective DMI by each genera’s relative abundance (see Methods), we found a notable increase of individuality in the stool microbiomes of IR individuals (**Fig. S12**), likely due to the increased *Bacteroidetes* among IR individuals.

Overall, the computation of DMI and FS for each specific genus offers an overarching perspective on microbial host specificity. Meanwhile, it provides crucial insights into the taxonomic composition of the community and potential influences of environmental factors on the host’s microbiome. Additionally, the DMI measurements provide important ecological characteristics about micro-biotypes, such as those related to ‘enterotype’ in stool microbiome ^78,117^ or ‘cutotypes’ in skin microbiome^63^.

### Interplay of Microbial Individuality and Stability: Insights from Longitudinal Multi-Site Microbiome Analysis

Prior studies^4,93–95,118–120^ demonstrated that microbiome stability is highly personalized. Consequently, we explored the relationship between microbial individuality and stability across body sites. We first examined the genus recolonization rate, measured by ASV consistency when a genus is detectable after being undetected in one or more consecutive samples. The overall recolonization rate (measured by *1 - Pairwise Jaccard Distance*^121,122^) was significantly associated with DMI on all three body sites except for the oral microbiome (**Fig. 3a**), which intrinsically maintains a high recolonization rate. Such correlations were strongest in the nasal microbiome (**Fig. 3a**), potentially explaining the high ICC observed above. Surprisingly, no difference in recolonization rate was found between IR and IS individuals. (**Fig. S13**). Our results suggest that highly individualized strains are more likely to recolonize, validating a hypothesis raised by other fecal transplantation studies^123,124^ and implying its potential efficacy in treating IR-related dysbiosis.

**Figure 3.**
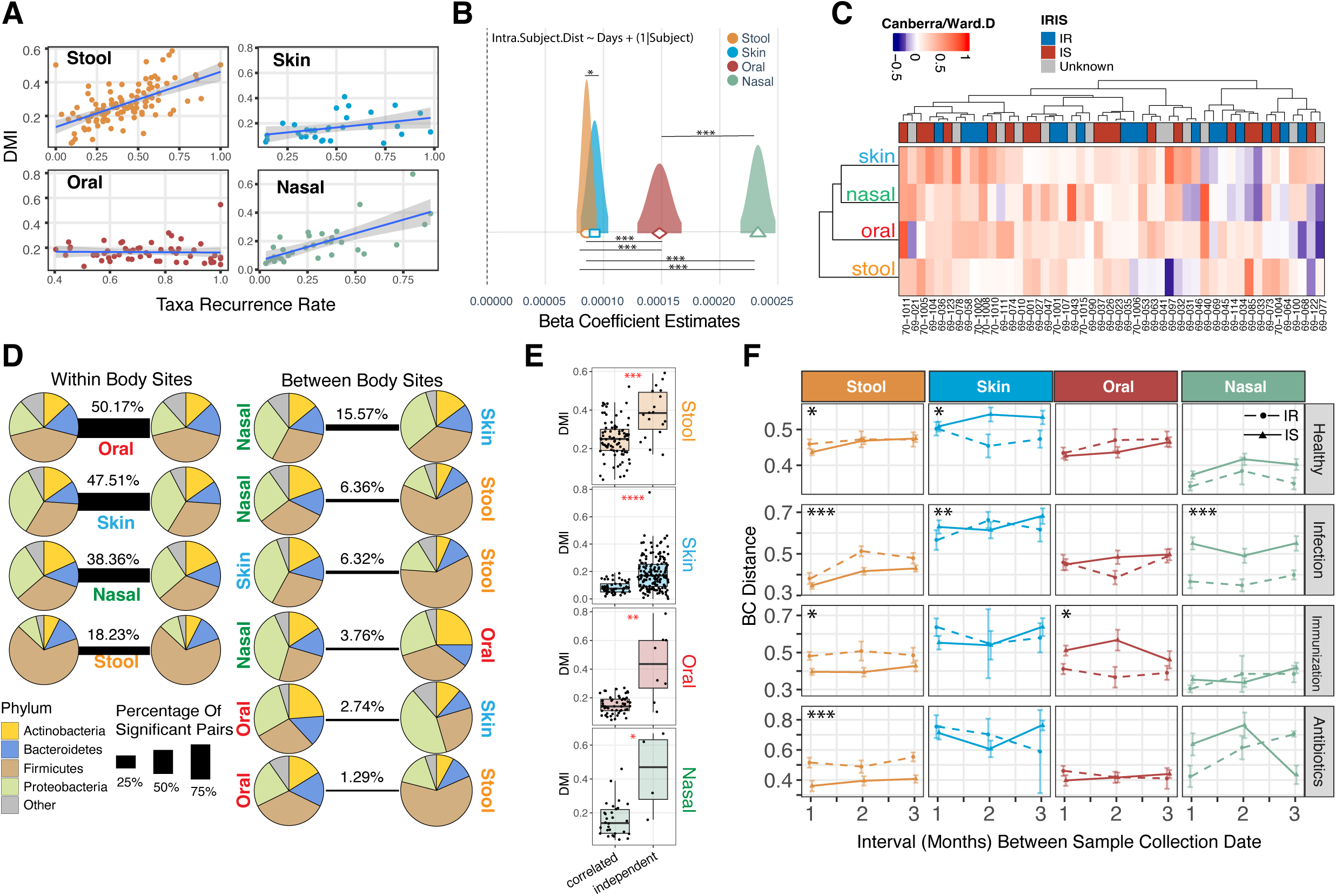
Temporal Stability of Microbiomes Associated with Individuality and Disrupted by Stress Events. A. Correlation Between Taxa-Recurrence Rates and Mean DMI: A correlation plot was created between the taxa-recurrence rates (measured by 1-Jaccard Distance) in all intra-individual samples and genus pairs (X-axis) and the mean DMI of this genus (Y-axis). The spearman correlation coefficient (rho) and p-value for each correlation are: Stool (n = 102): rho: 0.62, p-value = 3.07 × 10-12. Skin (n = 29): rho = 0.37, p-value = 0.027. Oral (n =54): beta estimate = -0.216, p-value = 0.1133. Nasal (n = 31): beta estimate = 0.69. P-value = 1.83×10-5. B. Relationship Between Dissimilarity and Collection Date Interval: The dissimilarity and collection date interval of intra-individual sample pairs were analyzed. Linear regression was estimated using a linear mixed-effects model. The beta coefficient distribution associated with the date interval was plotted, and a cross-body site comparison was made by setting the body site variable as an interaction term. Adjusted p-values were annotated as *: p < 0.05; **: p < 0.01; ***: p < 0.001. C. Inter-Body Site Correlation of BC-Distance Trends: A heatmap was constructed to represent the individual-based correlation coefficient between sample pair’s BC distances and the collection date intervals. Hierarchical clustering was performed using the Canberra distance and Ward’s linkage method. Insulin sensitivity status was marked at the bottom of the column clustering dendrogram (IR [blue]: Insulin Resistant; IS [red]: Insulin Sensitive; Unknown [gray]: do not have insulin suppression test results). D. Spearman Correlation Between Intra-Individual Microbiome Relative Abundance: Correlations within the same body site and between body sites of the same individual were represented. A pie chart was used to display the percentage of phylum among significantly correlated pairs. The percentage indicated the number of significant correlations relative to the total number of possible correlation pairs. E. Comparison of DMI Between Correlated and Non-Correlated Genera: The DMI between the genera that are correlated with each other and those that do not correlate with each other within a given body site were compared. A Two-Sided Wilcoxon Rank test was performed to test the difference in DMI, and p-values were annotated as *: p < 0.05; **: p < 0.01; ***: p < 0.001. F. Microbiome Shifts from Healthy to Stress Events Over Three Months: The BC distance of intra-individual sample points for all subjects at 1, 2, and 3 months, grouped by their insulin sensitivity stage, was examined. Comparisons were made for each body site during healthy-to-healthy time points, healthy to infection time points, healthy to immunization time points, and healthy to antibiotics time points. A two-way ANOVA was performed to determine if IR and IS show significantly different mean values of BC distance. BH-adjusted p-values are annotated as follows *: p < 0.05; **: p < 0.01; ***: p < 0.001.

The longitudinal data also enabled us to evaluate microbiome stability over time by tracking the dissimilarity between sample pairs in relation to collection date-intervals, which was reported to be higher in IBD-related gut dysbiosis^15^. Our analysis revealed that the stool microbiome changed more slowly over time, with the nasal site exhibiting the fastest rate of change (*p-value* < 0.001) (**Fig. 3b**). Additionally, IR individuals showed significantly lower stability in stool and skin microbiomes than IS individuals, as evidenced by linear mixed models (Stool *p-value*: 1.82 × 10-06, Skin *p-value*: 2.84 × 10-12), corroborating our findings of greater microbial abundance disparities in these samples between IR and IS participants. (**Fig. S10**).

### Intra- and Inter-Individual Correlations of Microbiome Dynamics Across Body Sites: Implications for Microbial Interdependence and Territory Specificity

We next investigated whether microbiome dynamics were co-associated across body sites within each individual, both at the community level and for individual taxa. Hierarchical clustering demonstrated a strong link in personal microbiome dynamics between the skin and nasal sites, whereas the dynamics of the stool microbiome was less correlated with other body sites on an individual level. (**Fig. 3c**). The correlation of microbiome stability across body sites indicates systemic microbial coordination, potentially regulated by the common mucosal system^125,126^. Comparison of time-related stability correlations across body sites between IS and IR individuals, revealed that both groups had a strong skin-nasal link, but a skin-oral correlation was exclusive to IS individuals. (**Fig. S14**). This implies that IR status could undermine the stability of the skin or oral microbiome, possibly due to the compromised host regulation of microbiome stability or due to impaired mucosal immunity related to IR^127^, a key factor for between-body-sites microbial interactions^125,126,128,129^.

Considering that the microbiome often operates as inter-dependent guilds^130–132^ and that intra-individual microbiome changes typically involve multiple taxa, we conducted a systematic analysis of longitudinal dynamics among all bacterial members per body site to determine which body sites demonstrate the highest microbial inter-dependency. Consistent with previous study of the stool microbiome, we found that 18.23% of all stool genus pairings (5,671) were significantly correlated within individuals, mostly involving genera *Firmicutes* (**Fig. 3d**). Surprisingly, co-association was highest in the oral microbiome (50.17%), followed by skin (47.51%) and nasal (38.36%), highlighting strong synergistic interactions among microbiome members at each body site. (**Table S8**)

We further investigated the microbial crosstalk between body sites by searching for sample-wise microbial collinearity between body sites. As expected, members from the skin and nasal sites were the most correlated (15.57% of all possible pairs) (**Fig. 3d, Table S8**). However, remarkable territory specificity was still evident among core microbiomes of each body site (**Table S9**). For instance, three core genera - *Corynebacterium*, *Staphylococcus*, and *Cutibacterium* - exhibited no longitudinal correlation between skin and nasal sites, despite being the predominant genera in both sites. Similarly, oral *Prevotella*’s abundance did not correlate with its abundance in the stool microbiome, consistent with our observation of high FS in oral but not stool *Prevotella* (**Table S7**). Although dysbiotic microbiome translocation between body sites has been reported^133–135^, our results suggest that these translocation cases likely do not involve the core taxa specific to a particular body site. Moreover, genera that are more interdependent across all body sites exhibit lower DMI compared to less interdependent genera (**Fig 3e, Fig. S15**), indicating that highly individualized microbiomes exhibit greater temporal stability^93,136^. Expanding on our findings of micro-biotypes across body sites (**Fig S1**), we observed that dominant taxa driving these biotypes exhibit significant inter-individual correlations. Examples include significant correlations of *Cutibacterium* levels in one’s skin with their *Cutibacterium* level in nasal (beta = 0.56, *BH-adjusted p-value = 0.0012*), their *Bacteroides* in stool (beta = 0.52, *BH-adjusted p-value = 0.0038*), and their *Leptotrichia* in oral microbiomes (beta = 0.43, *BH-adjusted p-value = 0.0088*). Conversely, individuals with high Unclassified *Ruminococcaceae* in stool correlates with their low *Veillonella* in oral microbiome (beta = -0.35, *BH-adjusted p-value = 0.015*) (**Table S10**). These findings clearly suggest that the establishment of micro-biotypes are governed by personalized factors.

### Acute Events Impact Microbiome Dynamics Across Body Sites and are Influenced by Insulin Resistance

We next examined the impact of three strong perturbation events - infection, vaccination, and antibiotic usage - on the microbiome, events known to cause stool microbiome disruptions^70,76,125,137^. Within a three-month period, the stool microbiome of IR individuals exhibited greater temporal shifts between healthy visits and perturbations compared to IS individuals (**Fig. 3f**). This pattern, however, was not consistently discernible across other body sites, such as the oral microbiome during vaccinations or the nasal microbiome during infection, suggesting a site-specific microbiome response to these events. Unlike IS individuals, IR individuals displayed reduced stool microbiome evenness and lacked the variability in nasal microbiome evenness observed in IS individuals during respiratory infection (**Fig. S16**), possibly due to higher IR-associated mucosal inflammation^138–140^ masking or superseding infection-induced local microbiome changes observed in IS individuals. During the course of infection, we identified a transient increase in several genera such as *Alistipes* in the stool, *Peptoniphilus* on the skin, Unclassified *Prevotellaceae* in the oral cavity, and *Oribacterium* in the nasal cavity. Conversely, we noticed a decrease in the presence of *Clostridium* cluster IV in the stool, Unclassified *Clostridia* in the oral cavity, and Unclassified *Neisseriales* in the nasal cavity (**Fig. S17, Table S11**). Although the extent to which transient microbial shifts contribute to IR-related dysbiosis is not clear, the notable increase of core microbiomes in IR skin samples (**Fig. S9**) may suggest a correlation between acute perturbations and long-term changes in the core microbiome composition. Overall, our findings indicate that dysbiosis can manifest differently across body sites, potentially through site-specific mechanisms. For instance, IR-related temporary disruptions in the stool microbiome seem to be characterized by a loss of core microbiome species producing short chain fatty acids. In contrast, in the less complex skin and nasal microbiomes, dysbiosis might involve the acquisition of opportunistic pathogenic species such as *Peptoniphilus*.

### The Interplay Between Host Immune System and Microbiome Across Body Sites: Insights Into Insulin Resistance and Inflammation

The dynamics of microbial stability are likely to be affected by multiple host regulatory mechanisms, particularly the immune system^44,129,141–144^. We profiled a panel of 62 circulating cytokines, chemokines, and growth factors to assess the immune status of each individual at each timepoint. Using our previous developed model^145^, we examined the interplay between host circulating cytokines and microbiome abundance at all four body sites. Overall, we identified 477 stool, 226 skin, 318 oral, and 221 nasal significant microbiome-cytokine associations based on Credible Interval (CI) that are highly body-site-specific (**Fig. 4a**, **Table S12**). The stool and oral microbiome showed a significantly broader microbiome interaction spectrum than skin and nasal microbiome. (**Fig. 4a**). Interestingly, the cytokines associated with epithelial/endothelial growth and vascular inflammation (EGF: 42 interactions, VCAM-1: 39 interactions, IL-22: 39 interactions), IL-1 family members (IL-1b: 44 interactions, IL-1Ra: 34 interactions), and leptin (39 interactions) demonstrated the highest number of interactions with the microbiome. Beta correlation coefficient comparison across all cytokines further identified a subgroup of cytokines including IL-1B, IL-1Ra, MCP3(CCL-7), and IL-23 as the strongest correlative cytokines with the microbiome (**Fig. S18**). The clear pattern of body-site-specific interactions may explain the niche specificity of genera. For example, *Moraxella* shows a negative correlation with 23 cytokines on the skin, yet only with three in the nasal cavity (**Table S13**). This reduced interaction in the nasal cavity, suggesting a lower immune response, could account for the higher prevalence of *Moraxella* in this location^7,146^.

**Figure 4.**
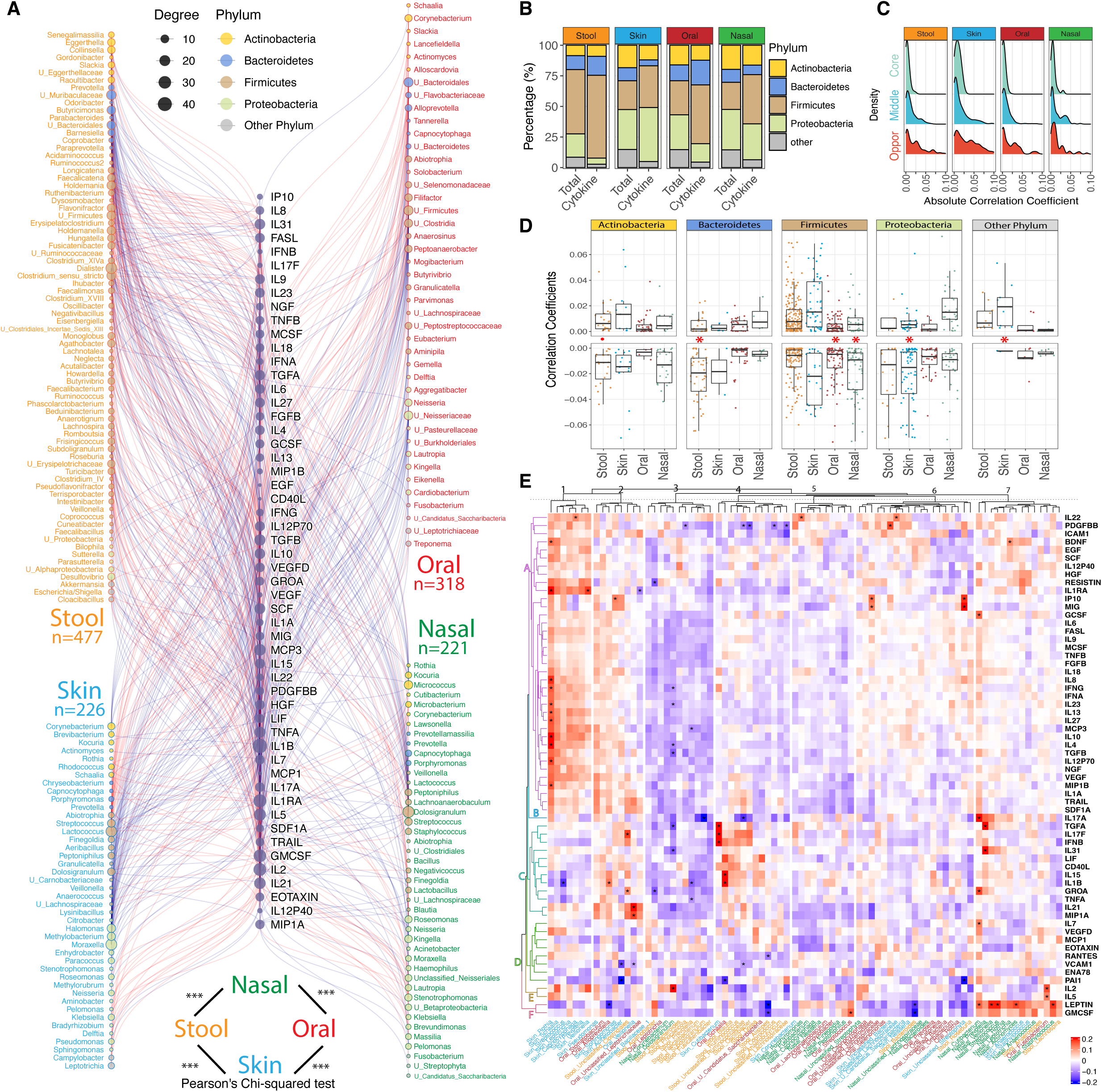
Systematic Connections Between Circulating Cytokines and Microbiomes. A. A customized mixed-effects model was used to evaluate the correlation between cytokines and microbiomes from various body sites. The association between a particular cytokine and microbiome is depicted by a red line for positive coefficients and by a blue line for negative coefficients. The circle size corresponds to the number of significantly correlated cytokines with the microbiome genera for which the absolute correlation coefficient is greater than 0.005. The number of significant correlations was annotated below each body site, and comparisons between the probability of identifying significant correlations between body sites were plotted at the bottom, and a Pearson’s Chi-squared test was performed with significance being annotated as *: p < 0.05; **: p < 0.01; ***: p < 0.001. The origin of the annotated genus’ body site is indicated by the color of the genus label. (Orange: Stool; Blue: Skin; Red: Oral; Green: Nasal). B. Percentage of Cytokine-Related Genera by Phylum: The percentage of cytokine-related genera was summarized by their phylum for all genera within a body site (Column: Total) and only genera significantly associated with cytokines (Column: Cytokine). Chi-square test between *Firmicutes* occurrence in Total and Cytokine column were performed, and the statistics are: X_stool_^2^ = 19.343, P_stool_ = 1.092 × 10^-5^; X_skin_^2^ = 10.418, P_skin_ = 0.001248; X_oral_^2^ = 30.935, P_oral_ = 2.668 × 10^-8^; X_nasal_^2^= 31.396, P_nasal_ = 2.104 × 10^-8^ C. Association Between Low-Prevalence Microbiomes and Cytokines: A density plot displays the absolute value of the correlation coefficient for all pairs of cytokine and microbiome that are significantly associated. The result was separately plotted based on the mean prevalence of the genus. Comparisons between the correlation coefficient (absolute value) of core genera and opportunistic genera were performed using the Two-Sided Wilcoxon Rank test.: stool: W = 1,715, p-value = 1.181×10-12, skin: W = 223, p-value = 0.004041, oral: W = 1,055, p-value = 7.877×10-6, nasal: W = 482, p-value = 0.004691. D. Comparison of Correlation Coefficient by Body Site and Phylum: The comparison of correlation coefficients by body site and phylum was made. Adjusted p-values in the middle row represent the Wilcoxon signed rank sum test between correlation coefficients that are positive and absolute values of all negative correlation coefficients in the same column. E. Spearman Correlation Coefficient Between Cytokines and Diverse Genera: The Spearman correlation coefficient was calculated between cytokines and the most diverse 20 genera in each body site. The column and row were clustered using hierarchical clustering. Annotations on the bottom are color-coded based on the body sites to which each genus belongs. (Orange: Stool; Blue: Skin; Dark Red: Oral; Green: Nasal)

We next examined whether certain phyla frequently interact with cytokines, potentially influencing their stability and individuality. Members of the *Firmicutes* phylum (primarily *Clostridia*) were significantly overrepresented among cytokine-correlated microbes across all body sites (**Fig. 4b**), indicating their close relationship with the host immune system. This interaction may account for their higher FS and lower DMI in the stool microbiome compared to other phyla. Additionally, an increase of *Firmicutes* is associated with conditions like obesity-related gut dysbiosis^147^, IBD-related oral dysbiosis^148^, and psoriasis-related skin dysbiosis^149^, which share inflammation as a common factor.

Interestingly, cytokines appear to play a pivotal role in shaping an individual’s core microbiome and in curbing the colonization of non-commensal bacteria, including many from the *Proteobacteria* phylum. At all four body sites we found that opportunistic microbes (longitudinal prevalence < 20%) demonstrated a stronger correlation with cytokines than core microbiomes (longitudinal prevalence > 80%) (**Fig. 4c**). This correlation is largely driven by *Proteobacteria* rather than *Firmicutes*, as *Proteobacteria* consistently constitutes a larger segment of the opportunistic microbiome compared to the core microbiome (**Fig. S19**). *Proteobacteria*, known for their high immunogenicity, often carry potent lipopolysaccharides (LPS) and instigate the downstream immune cascade^150–154^. Previous studies^125,155,156^ have reported an increase in *Proteobacteria* abundance during inflammation. Contrary to these findings, our study revealed that the correlations between cytokines and *Proteobacteria* abundance are mostly negative, except for *Proteobacteria* members in the nasal microbiome, which exhibit no significant difference between positive and negative correlations (**Fig. 4D**). The negative correlation of *Proteobacteria* is stronger in stool (W = 25,315, p-value = 0.04439), skin (W = 4,995, p-value = 0.005214), and oral sites (W = 9,443, p-value = 9.649×10-5), but weaker in nasal sites (W = 5,218, p-value = 0.2693). Furthermore, all cytokine correlations (n =10) from opportunistic stool *Proteobacteria* are negative, while many high prevalence *Proteobacteria* members exhibit positive correlations with cytokines (**Fig. S20**). Our results suggest that inflammation might contribute to the adaptation of more prevalent (or individualized) *Proteobacteria*^125,157^.

The host response by cytokines and chemokines may be linked with the observed richness (ASV complexity) of bacterial genera, in addition to their relative abundance. To explore this bi-directional relationship, we examined the correlation between cytokines and the richness of the 20 most diverse genera per body site. We found that the richness of several prevalent stool microbiome genera within the *Bacteroidetes* phylum, such as *Prevotella*, *Phocaeicola*, and *Parabacteroides*, form a cluster (Column 3, **Fig. 4E**) with primarily negative associations, in line with our finding that the relative abundances of *Bacteroidetes* members are more likely to be negatively correlated with cytokines (**Fig. 4D**). We also observed that leptin and GM-CSF, both strongly associated with BMI (**Fig. S21**), show the strongest overall correlation with richness (Row Cluster F, **Fig. 4E**). This extends findings from numerous studies^147,158–160^ on obesity’s influence on the gut microbiome diversity to additional body sites. Furthermore, we discovered that seven skin genera (column cluster 1, **Fig. 4e**) positively correlated with a cytokine cluster (row cluster A, **Fig. 4e**), indicating that the richness of specific skin genera (e.g., *Rothia*, *Veillonella*, *Streptococcus*) is positively associated with the level of plasma cytokines. Consequently, the observed increase in the skin microbiome richness in IR individuals (**Fig. S6B**) may suggest a diminished host selection of the skin microbiome during inflammatory periods. Overall, our analysis elucidates the interplay between chronic low-grade inflammation and microbial dynamics across various body sites. It highlights the role of the immune system on microbiome composition and individualization, and how these dynamics are both influenced by and contribute to insulin resistance.

### The Microbiome is Highly Connected with Host Molecules: Unraveling the Role in Insulin Resistance and Inflammation

To more generally explore the relationship^67,161,162^ between the microbiome and internal host molecules and its role in IR, we examined the correlations between microbiome genera and plasma proteins, lipids, and metabolites in the host. We first modularized the lipidomics data to address its high collinearity (**Fig. S22, Table S14**), then applied a linear mixed model to residualize all omics data (see Methods), reducing between-individual variation and emphasizing longitudinal intra-individual relationships.

Interestingly, the microbiome-host molecule interaction network partitions according to internal molecular composition, rather than microbiome body sites (**Fig. 5A**), suggesting that certain taxa are primarily influenced by their interactions with internal molecules, rather than the individual taxa driving the host molecular composition. Notably, the key taxa that are typically associated with enterotypes: *Bacteroides*, *Prevotella*, and Unclassified *Ruminococcaceae*, exhibit a clear preference for the lipidome, proteome, and metabolome regions, respectively (**Fig 5a, Table S15**). The close association between *Prevotella* and proteins has been previously documented^163,164^, as well as the relationship between Bacteroides and lipids^165,166^. However, our findings extend this knowledge to include both additional taxa and multiple body sites, suggesting that these connections are not only site- and taxa-specific but also systemic and robust.

**Figure 5.**
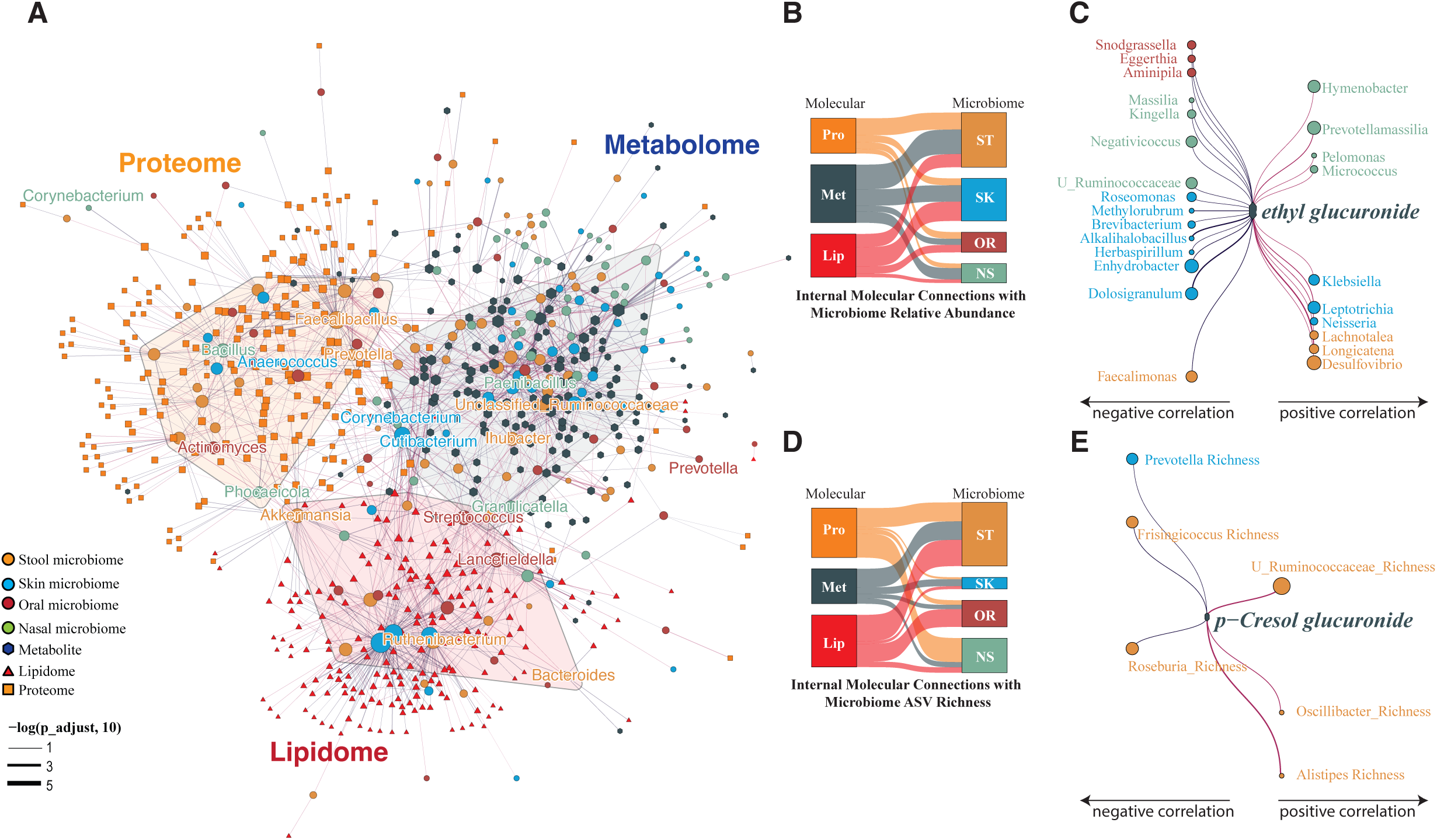
Interactions Between Plasma Metabolites, Lipids, Proteomics, and Microbiome Over Time. A. Network of Correlations: Displayed is a network showing correlations between the relative abundance of microbiome genera at four body locations and plasma analytics. The confidence of the correlation is represented by lines between nodes. Microbiomes (Dark yellow filled circle: Stool; Blue filled circle Skin; Dark red filled circle: Oral filled circle; Green: Nasal) and plasma analytics (dark blue filled hexagon: Metabolome; orange filled square: Proteome; Red filled triangle: lipidome) was color coded. B. Summary of Correlations Between Plasma Analytics and Microbiome Relative Abundance: This summary shows the number of significantly associated pairs between plasma analytics and microbiome relative abundance, indicated by the heights of the squares and the thickness of lines. Pro (light orange): Proteome; Met (Gray): metabolome; Lip (red): lipidome, ST (dark orange): stool microbiome; SK (blue): skin microbiome; OR (dark red): oral microbiome; NS (light green): nasal microbiome. The number of significantly (BH-adjusted p-value < 0.2) associated pairs is shown by the heights of the squares and thickness of lines. C. Correlations Between Genera and the Metabolite Ethyl Glucuronide: Presented are correlations between genera (relative abundance) and the metabolite ethyl glucuronide. Genera with a positive correlation are placed on the right with red lines connected, while those with a negative correlation are on the left with blue lines connected. The size of the dots indicates the number of significantly associated pairs that are related to each genus. D. Summary of Correlations Between Plasma Analytics and Microbiome Observed ASV Richness: This summary shows the number of significantly associated pairs between plasma analytics and microbiome observed richness. Details can be referred to in the legend section of part b. E. Correlations Between Genera and the Metabolite p-Cresol Glucuronide: Shown are correlations between genera (observed richness) and the metabolite p-Cresol glucuronide. Genera with a positive correlation are placed on the right with red lines connected, while those with a negative correlation are on the left with blue lines connected. The size of the dots indicates the number of significantly associated pairs that are related to each genus.

**Figure 6.**
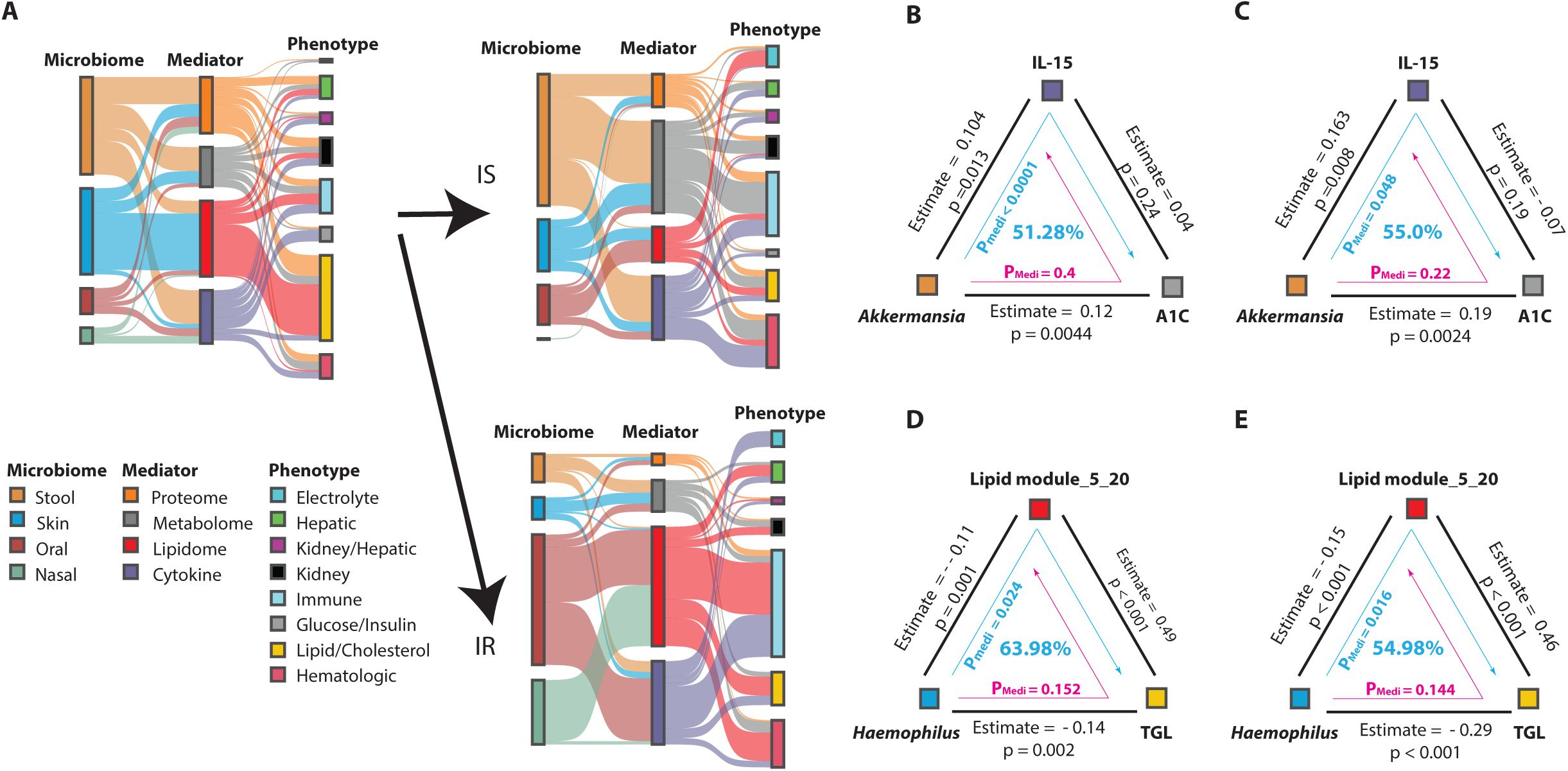
Causal Inference Decodes Microbiome-Driven Phenotypic Modifications Mediated by Internal Molecules and Cytokines. A. Microbiome and Phenotype Linkage Analysis: The diagram illustrates the linkage analysis mediated by internal molecules and cytokines. Each column’s color denotes the body site of the microbiome as modulator (left), the type of analytics as mediator (middle), and the class of phenotypes as consequence (right). The heights of each column represent the number of detected associations. The left upper panel displays mediation linkage for the entire dataset, while the right panel displays the same type of mediation linkage for only insulin-sensitive (IS) and insulin-resistant (IR) participants. Comparisons between IS and IR regarding each mediated effect were performed using a Fisher exact t test. B. *Akkermansia*’s Mediation Effect on Blood A1C Level via Plasma IL-15: The relative abundance of Akkermansia from the stool microbiome is shown to causally contribute to the blood A1C level via plasma IL-15 among all individuals, with a significant mediation effect: Pmedi < 0.0001, 51.28% mediation effect. C. *Akkermansia*’s Mediation Effect on Blood A1C Level in Insulin Sensitive Individuals: A similar mediation effect is observed specifically in insulin-sensitive individuals: Pmedi =0.048, 55.0% mediation effect D. *Haemophilus*’s Mediation Effect on Plasma Triglycerides Level: The relative abundance of Haemophilus parainfluenzae from the skin microbiome is demonstrated to causally contribute to the plasma level of triglycerides among all individuals, with a significant mediation effect: Pmedi = 0.024, 63.98% mediation effect. E. *Haemophilus*’s Mediation Effect on Plasma Triglycerides Level in Insulin Sensitive Individuals: This mediation effect is also apparent in insulin-sensitive individuals: Pmedi =0.016, 54.98% mediation effect.

Many taxa-molecule interactions were consistent across different body sites, validating the robustness of these relationships. For instance, *Haemophilus* from both skin and oral sites demonstrated a strong connection with the lipidome, while Unclassified *Ruminococcaceae* from stool and nasal sites were primarily associated with the metabolome. The skin microbiome exhibited the highest connectivity in the microbe-lipidome association, whereas the stool microbiome was mainly connected to the metabolome and proteome, as previously reported^167,168^. (**Fig. 5b**). Pathway enrichment analysis of proteins linked to microbiome further supported these connections and their potential functional implications. The skin microbiome-related proteins were enriched for pathways regulating lipid metabolism and transport, whereas the stool, nasal, and oral microbiomes were more strongly connected to the host immune response, including complement activation and humoral immune response. Interestingly, the oral microbiome exhibited significant associations with proteins linked with regulation of proteolysis, hydrolase activity, and enzyme, peptidase, and lipase inhibitor activity. **(Fig. S23**). Strikingly, the complexity of the network between the stool microbiome and host molecular relationships was significantly reduced in individuals with IR (**Fig. S24**), indicating a loss of balance in these individuals.

Our study reveals several metabolites, such as alcohol associated metabolite ethyl glucuronide^169^, interact with microbiomes across all four body sites, demonstrating a more global effects of alcohol beyond the gut and oral microbiome^170–172^. Notably, skin *Neisseria* and *Klebsiella*, recognized for their acetaldehyde-^173,174^ and ethanol-producing^175,176^ potential, showed a positive correlation with plasma ethyl glucuronide level (**Fig. 5c**). Conversely, *Faecalimonas*, typically an acetate-producing bacteria^177,178^ and sensitive to alcohol^179^, exhibited a significant negative correlation with ethyl glucuronide. Furthermore, *Desulfovibrio*, a key genus promoting microbiome-related metabolic syndrome^180,181^, was positively correlated with ethyl glucuronide. These findings suggest that alcohol metabolism is associated with microbial dysbiosis across various body sites.

We also observed notable interactions between microbiome genera richness, a recognized indicator^182,183^ of host metabolic status, and internal plasma analytics. These interactions were most pronounced in the stool microbiome, likely due to its high richness. (**Fig. 5d, Fig. S25, Table S16**). Intriguingly, we discovered a positive association between p-Cresol glucuronide, a bacteriostatic^184^ metabolite produced exclusively by the anaerobic gut microbiome and linked to insulin resistance^27,185^, and the richness of Unclassified *Ruminococcaceae* and *Oscillibacter* in the stool. Conversely, it was negatively associated with genera typically considered metabolically beneficial, such as *Roseburia*^186–188^. (**Fig. 5e**) These findings suggest that p-Cresol glucuronide may influence the development of insulin resistance-related dysbiosis by altering host tolerance to certain genera belonging to *Clostridia*.

In conclusion, our results not only validate several previous findings but also generate many novel host-microbial associations, enhancing our understanding of the complex interplay between the human microbiome, metabolites, and host health.

### Elucidating Microbiome-Mediated Effects on Clinical Phenotypes: A Comprehensive Mediation Analysis across Four Body Sites

We next investigated potential causal linkages between microbes, host molecules and clinical phenotypes using mediation analysis^189,190^. This analysis quantifies the extent to which a mediator variable, such as internal omics or cytokines/chemokines, contributes to the relationship between an independent variable (the microbiome) and a dependent variable (clinical phenotypes, **Table S17**), and provides a confidence level, derived from comparing the association results when the roles of the mediator and the dependent variable are reversed^56,189,191^. Using the assumption that the microbiome impacts clinical readouts, we specifically probed for interactions likely mediated by internal analytes such as metabolites^192^. Based on this premise, we identified 330 directionally significant mediation effects involving microbial taxa across all four body sites, with 207 and 164 of these mediation effects detected in IS and IR participants, respectively (**Table S18)** These outcomes suggest potential causal relationships between the microbiome and host health exist and vary with IR status, without ruling out the influence of unidentified confounding variables. Compared to IS, we observed a significant decrease in cytokine-mediated effects of stool microbiome on hematologic parameters (p-value = 0.002) and metabolome-mediated effects of stool microbiome on immune phenotypes (p-value = 0.002) in IR individuals. Furthermore, there was an absence of lipidome-mediated associations between skin microbiome and host plasma lipids/cholesterol in IR participants (p-value = 0.031), suggesting a dysregulation in specific microbiome-metabolic interactions related to hematologic/immune and lipid/cholesterol homeostasis in IR individuals. **(Table S19)**

In contrast, the oral microbiome mediated a large proportion of immune profiles via the modulation of lipidome (p-value = 0.035) and cytokines (p-value = 0.0001) in IR relative to IS participants, primarily through a negative relationship among major oral core microbiomes such as *Veillonella* (**Table S18**), These causal relationships align with previous findings suggesting that diabetes-related oral dysbiosis may arise from a combination of the loss of core commensal oral taxa^34,37,193^ and an increase in the pathogenicity of resident oral bacteria during impaired glucose metabolism, as demonstrated by human observational studies^194,195^ and animal research^36,40^.

## Discussion

This study presents the first systematic analysis of multi-site microbiome ecology and their host relationships among the HMP prediabetic cohort. With date-matched microbiome and host omics data, we not only expand our knowledge of the ecology, stability, and individuality of microbiome from various body sites, but also provide mechanism-generating hypotheses on host-microbiome interactions in the context of T2D risks. We demonstrate a number of novel observations: 1) There is a “core” microbiome that is highly stable (in terms of presence) over time and an opportunistic microbiome that is highly variable and more likely to interact with the immune system; these are body-site-specific with some consistent correlations of microbes between sites. 2) Correlations between microbiomes across body sites and extensive interactions with host factors indicate systemic coordination and interactions across the human body. 3) Highly individualized microbiomes are differentially associated with distinct environmental factors (i.e., season, diet, chemical and biological exposome), and presumably these factors shape the microbiome in a body sites-specific manner. However, these effects do not override the variance contributed by individuals, suggesting that the host is still the largest confounding factor for the variation observed in the microbiome. 4) Finally, individuals with IR have a less stable microbiome with more diverse microbiome members on skin, possibly associated with upper respiratory infection, as well as significantly altered host-microbiome interactions.

Our study revealed that the stool and oral microbiomes exhibit the highest level of individualization, likely due to the unique influence of personalized dietary habits and host-specific factors, such as IR, impacting the digestive system^196^. On the other hand, the skin and nasal microbiomes are less individualized, possibly owing more to individual environmental exposure^90,113^. We observed site-specific impact of environmental factors, such as seasonal variations, on the microbiomes of these four body sites. For instance, a decrease in stool microbiome richness in late summer (September) corresponded with previous findings of worsened insulin sensitivity during this period^197,198^. Similarly, we noted a decline in the richness and evenness of the oral microbiome from late summer through winter, suggesting a potential influence of environmental factors like the availability of fresh food and changes in sunlight exposure (which affects Vitamin D synthesis). Meanwhile, changes in humidity from January to April were associated with an increase in skin microbiome richness and a decrease in nasal microbiome richness^90^. These observations underscore the critical role of the host in modulating the microbiome in response to environmental fluctuations and highlight the significance of longitudinal environmental factors in shaping the microbial community.

Leveraging dense longitudinal sampling, we were able to quantify the stability and degree of individuality of the microbiomes at different body sites. We found a strong correlation between microbiome stability and individuality, suggesting an active role of the host in the establishment of commensal bacterial populations. Notably, this individuality appears to be taxon-specific, pointing to the possibility that the stability of an individual’s microbiome may be influenced by the dominant bacterial taxa they carry. For instance, we observed a higher degree of microbiome individuality (DMI) in stool samples for members of the *Bacteroidetes* phylum, and that the *Bacteroidetes/Firmicutes* ratio had novel biological implications regarding their stability, and providing another mechanistic interpretation to concept of enterotypes^50,77,78^, especially the new Bac2 enterotype which has been closely linked to metabolic disorders^117,199^. Furthermore, we found a strong correlation between microbiome micro-biotypes across different body sites within the same individual, with individuals showing high levels of *Bacteroides* in the stool also having increased *Cutibacterium* on the skin and *Prevotella* in the oral microbiome. Combined with our findings on the correlation between microbiome individuality and stability, the low-level engagement between the core microbiome and cytokines, and the clear host molecular type-specific patterns in the host-microbiome interactome, we propose that the colonization and adaptation of bacteria across multiple body sites is not a random process. Rather, host factors play an active role in these processes, maintaining a certain level of equilibrium in the microbiome community.

Our cross-site comparisons underscore the site-specific nature of the microbiome. For instance, we found that the recurrence rate of oral bacteria does not increase with DMI, possibly due to a stable and conserved community structure within the oral microbiome across different individuals^83,200^. This, coupled with the oral microbiome’s high intra-site microbial interdependence and minimal correlation with other body sites, underscores its specialization. The unique oral environment, shaped by factors such as saliva, teeth, gums, tongue, and distinct nutrient availability, likely contributes to this specialization^201,202^.

One important system for driving microbial individuality and stability is the host immune system. The immune system is well known to interact with microbes at multiple body sites^141,203,204^, and this interaction modulates both the microbes that are present, as well as their functional benefits (e.g., beneficial microbial signals such as those that maintains barrier functions^44,205^). Our longitudinal Bayesian model reveals that the interactions between the microbiome and cytokines, while present, are subtle. Specifically, certain genera exhibit an approximate 1.5-fold change in response to cytokine variations. The interaction between inflammatory cytokines and the microbiome demonstrated that low prevalence genera, such as those belonging to the stool *Proteobacteria*, are likely reduced during host inflammatory events. We also revealed a systematic relationship between cytokines and the genera complexity of the microbiome at each body site. Notably, the diversity within a subset of the skin microbiome positively correlates, while that within the stool microbiome negatively correlates with the same group of cytokines. Given the established association of IR with low-grade chronic inflammation^206^, we believe that the observed changes in diversity among IR individuals are related to their associated cytokine profiles.

We also identified a surprisingly large number (2180) of interactions between host plasma biomolecules and microbiomes from different body sites (**Table S15**). Some of the relationships, such as alcohol metabolites and the gut microbiome, have been previously documented^207,208^. Stool *Bacteroides*, *Prevotella* and Unclassified *Ruminococcaceae* are separately located in lipidomics, proteomics and metabolomics zones of the correlation network, indicating that these host factors strongly interact with the different types of microbes. We also find many correlations across body sites with host factors; for example, the correlated *Bacteroides* in the stool and *Cutibacterium* in the skin are both closely related to lipid metabolism^209–213^. Our multi-omics analyses suggest potential causality of host factors in these relationships, reinforcing the idea of host-driven systematic microbial coordination across body sites as proposed in gut-brain axis ^214,215^ and gut-lung axis^216,217^ studies. Since many of the metabolites, lipids and proteins are signaling molecules (e.g., chemokines, hormones, peptides), these molecules may play important roles in organismal communication across the entire host-microbiome ecosystem. Importantly, this interaction presumably occurs at the individual-specific level, and significantly altered at disease stage.

Intriguingly, we found the relative abundance of *Klebsiella* on skin was positively associated with metabolites of alcohol. This indicates that alcohol intake may change the host into a *Klebsiella*-tolerant environment, resulting in the adaptation and expansion of pathogenic *Klebsiella* as previously described^218^. This correlative relationship supports previous findings in alcohol-associated pneumonia, where alcohol consumption and increased susceptibility to *Klebsiella* in the lungs may be a result of either intestinal *Klebsiella*-specific T-cell sequestration^218,219^ or alcohol-related impairment of tryptophan catabolite production/processing in the gut microbiome, which restricts pulmonary immune cell trafficking^220,221^.

Insulin resistance (IR) appears to disrupt the intricate balance between the host and microbiome, as demonstrated by an unstable, dysbiotic microbiome in IR individuals. In our study, we discovered marked differences in the microbiome composition, diversity, and core members of the stool and skin microbiomes in IR individuals compared to their insulin-sensitive counterparts. Notably, the systematic shift in microbiome prevalence indicates an entire microbial community’s transformation instead of the abnormality of a few isolated members (**Fig. S9B**). This dysbiosis can potentially alter the complex interaction between the microbiome and host in IR subjects. Consistently, we detected a decrease in host-microbe coordination and a missing association between oral and skin microbiome stability. Further, our mediation analysis validates our result about the reduced gut microbiome-cytokine interaction in IR individuals^70,145^, and revealed heightened pro-inflammatory signals linked to the microbiomes of the upper respiratory tract (oral and nasal microbiome) in IR individuals. Our in-depth exploration of acute microbiome changes during infection provides additional substantiation for this theory. We observed a significant increase in IR-enriched skin genera such as *Peptoniphilus* and *Intrasporangium* (**Fig. S15**) during the course of a respiratory viral infection, while a decline in specific genera in the stool microbiome known for butyrate production, like *Clostridium*_IV, *Lawsonibacter*, and *Intestinimonas*^222–225^. While these findings do not explicitly validate the hypothetic link^226^ between respiratory viral infection, chronic metabolic dysregulation, and related complications, they suggest that infection-related microbial shifts may be associated with, or even cause, the dysbiosis and complications of metabolic syndrome as observed in animal models^227^.

## Limitations of the Study

Despite the significant findings, our study has limitations. The DMI and FS calculations primarily rely on the Bray-Curtis dissimilarity. Therefore, study design such as sample size, population representation, choice of dissimilarity metric, and genus internal complexity may impact the accuracy and interpretation of DMI values. Additionally, although DMI can provide a quantitative measure of microbiome individuality, understanding its biological or ecological significance can be challenging due to the potential influence of host genetics, environmental factors, and host regulation.

When the study began, 16S rRNA sequencing was highly efficient for analyzing microbiomes rich in human DNA, such as those in nasal and skin swabs. However, our team, along with others, later identified its limitations in accurately capturing specific bacterial genera in stool and skin microbiomes when focusing on only certain variable regions.^73,228^. Furthermore, unlike metagenomic sequencing, 16S analysis does not provide information on viruses and fungi within a microbiome community, nor does it reveal gene function potentials. Also, our study does not capture a collective profile of the entire skin or oral microbiome, given the distinct characteristics of species at various sites^7,62,119^. Instead, it represents the microbiome from a single swab taken from each indicated body site. Furthermore, the geographical and lifestyle constraints of our participants could restrict the broad applicability of our conclusions to more diverse and complex populations^229,230^. Moreover, we note that our analyses primarily determine associations or statistical mediation effects, using p-values as cut-off points. Despite the bolstered study power afforded by a longitudinal design^231,232^, some observations with a smaller sample or effect size may still be overlooked. For example, our study’s insights into insulin resistance are based on a cohort size of 28 IS and 30 IR individuals, and even smaller for events like infection. Analysis of larger cohorts may reveal further insights. Furthermore, the use of p-values has its limitations regardless of the cut off^233^, such as its dependence on sample size,^243^, the potential for misinterpretation^234,235^, and an inflated type 1 error rate with multiple tests^236,237^. These factors should be taken into consideration when interpreting our results. Finally, while our study highlights notable associations and suggests potential causations, it does not establish causation. Our mediation analysis aims to infer directionality and potential causality, but unobserved factors could influence these findings. To confirm causative links, more targeted follow-up studies are essential.

In spite of these limitations our studies provide a number of novel observations concerning the individuality and stability of the microbiome across multiple body sites during health and disease and in individuals with different IR/IS status. These observations have important implications in modulation human molecular health using personalized prebiotics and probiotics. Our data also provides a valuable and unique resource for the general scientific community.

## Methods

### Participant Recruitment

Participant recruitment was conducted as part of the National Institutes of Health (NIH) integrated Human Microbiome Project, under the auspices of the integrative personal omics profiling (iPOP) study, registered with Stanford University’s Institutional Review Boards (#23602). Eligible participants were either at risk for type 2 diabetes or voluntarily interested in diabetes-related research. Exclusion criteria encompassed hypertriglyceridemia > 4.0 mg/ml^−1^, uncontrolled hypertension, uncontrolled psychiatric disease, previous bariatric surgery, pregnancy or lactation, eating disorders (i.e., binge eating disorder, anorexia nervosa, or bulimia nervosa), alcohol use disorder, or failure to provide five consecutive samples from at least one body site (stool, skin, nasal, or oral). Consequently, data from 86 participants met the criteria for inclusion in this analysis.

### Microbiome Sample Collection and Sequencing

Stool samples were self-collected by participants and other samples were collected by study coordinators following iPOP study standard operating procedures (SOP), as adapted from HMP_SOP corresponding sections (HMP_MOP_Version12_0_072910)^70^. Briefly, retroauricular areas were rubbed with pre-moistened swabs under pressure for skin sampling, anterior nares for nasal sampling, and rear of the oropharynx for oral sampling. Samples are stored at -80 C immediately after arrival. Stool and nasal samples were further processed and sequenced in-house at the Jackson Laboratory for Genomic Medicine (JAX-GM, Farmington, CT, USA) and detailed methods are described previously^70^, while oral and skin samples were sent to uBiome (uBiome, San Francisco, CA, USA) for further processing.

After 30 minutes of beads-beating lysis, skin and oral samples were processed using a silica-guanidinium thiocyanate-based nucleic acid isolation protocol^238–240^ on a liquid-handling robot. The 16S rRNA variable region V4 was amplified by 35 cycles of PCR using the primer 515F (5’-GTGCCAGCMGCCGCGGTAA-3’) and 806R (5’-GGACTACHVGGGTWTC TAAT-3’)^241^. The DNA from each sample was barcoded and combined to create a sequencing library. The sequencing library was then purified using columns and microfluidic DNA fractionation^242^ to reduce unwanted DNA fragments. Bio-Rad MyiQ was used to quantify the DNA concentration of the library using the Kapa iCycler qPCR kit (Bio-Rad Laboratories, Hercules, CA, USA). Sequencing was performed on the Illumina NextSeq 500 Platform (Illumina, San Diego, CA, USA) via 2 * 150 bp paired-end sequencing protocol^243^.

Raw sequencing data from the stool samples and nasal samples are acquired from our previous publication^70^. Briefly, 16S rRNA gene from V1∼V3 hyper-variable region was sequenced with primer pair of 27F (5′-AGAGTTTGATCCTGGCTCAG-3′) and 534R (5′-ATTACCGCGGCTGCTGG-3′) and being barcoded and sequenced on the Illumina MiSeq sequencing platform through a V3 2 × 300 sequencing protocol. The same cutoff used in skin and oral sequencing data was applied to stool and nasal sequencing data in demultiplexing. After demultiplexing, reads with Q-scores less than 35 and ambiguous bases (Ns) are trimmed for additional analysis.

### Microbiome data processing

Demultiplexed sequenced samples were saved as FASTQ files using BCL2FASTQ software (Version 2.20, Illumina, CA, USA). Sequences with barcode mismatches, primer mismatches exceeding one, or Q-scores below 30 were excluded. Due to low overlap between forward and reverse reads according to FLASH (Version 1.2.11), only forward reads were selected for further processing.

Microbiome sequencing data from four body sites were combined and processed using the DADA2 R package (version 1.16)^244^. Sequences were filtered to remove ambiguous bases (maxN=0) and those with more than two expected errors (maxEE=2). After filtering, inter-sample composition analysis was performed based on the learned error rate. An amplicon sequence variant (ASV) table was constructed, and chimeras were removed using the DADA2 workflow consensus method. Reads passing all filters were aligned against a trained database of target 16S rRNA gene sequences and taxonomic annotations derived from Version 18 of The Ribosomal Database Project (RDP)^245^ Taxonomy release (Aug 14, 2020). Relative ASV abundance was determined by dividing the count associated with that taxon by the total number of filtered reads. Samples with depths below 1,000 reads were removed due to insufficient sequencing depths^246^ following the HMP consortium standard^15,70^. The Local Outlier Factor (LOF) of each point was calculated on a sequencing depth to richness (observed ASV) plot, and samples with a LOF greater than 3 (n=7) were removed due to an abnormal richness-sequencing depth relationship. The average sample sequencing depth after quality control was 23,554 for stool microbiome, 74,515 for skin microbiome, 132,912 for oral microbiome, and 24,899 for nasal microbiome. Batch effects were estimated using PERMANOVA analysis when both batch and subject ID were included, with results showing that the total variance explained by the 23 batches was 2% for stool, 3% for nasal, 0.6% for oral, and 3.6% for skin samples. The batch effects were considered small, and no correction for batch effects was applied in the analysis.

### Lipidomics analyses

Lipid extraction and data generation were performed as previously described^247–249^. Briefly, complex lipids were extracted from 40 µL of EDTA-plasma using a mixture of methyl tertiary-butyl ether, methanol, and water, followed by biphasic separation. Lipids were then analyzed using the Lipidyzer platform, which consists of a DMS device (SelexION Technology, Framingham, MA, USA) and a QTRAP 5500 (Sciex). Lipids were quantified using a mixture of 58 labeled internal standards provided with the platform (cat# 5040156, Sciex, Redwood City, CA, USA), and lipid abundances were reported in nmol/g.

To address the high collinearity of the lipidomic data, a customized clustering method was designed. Specifically, the lipidomics data were divided into six clusters using Fuzzy c-means clustering (R package “Mfuzz” (version 3.15)). For the lipids within each cluster, correlations were computed, and lipids with high correlative relationships (Spearman correlation > 0.8 and BH-adjusted p-values < 0.05) were grouped into the same module. Community analysis (‘fastgreedy.community’ function from R package “igraph” (v1.3.5)) was employed to detect the modules. For lipids not assigned to any of the modules, their original lipid species annotations were used for downstream analysis (**Table S14**).

### Metabolomics Analyses

Untargeted metabolic profiling was performed using a broad-spectrum LC-MS platform using a combination of reverse-phase liquid chromatography (RPLC) and hydrophilic interaction liquid chromatography (HILIC) separations and high-resolution MS^70,250^. Briefly, plasma metabolites were extracted following solvent precipitation using a mixture of ice-cold acetone, acetonitrile, and methanol (1:1:1, v/v). Hydrophilic metabolites were separated on a ZIC-HILIC (2.1 × 100 mm, 3.5 μm, 200 Å; Merck Millipore) while hydrophobic metabolites were separated on a Zorbax SBaq columns (2.1 × 50 mm, 1.7 μm, 100 Å; Agilent Technologies). Data was acquired on a Thermo Q Exactive plus mass spectrometer for HILIC and a Thermo Q Exactive mass spectrometer for RPLC. Raw data were processed using Progenesis QI (v2.3, Nonlinear Dynamics, Waters) and metabolites were formally identified by matching fragmentation spectra and retention time to analytical-grade standards or matching experimental MS/MS to fragmentation spectra in publicly available databases. A total of 726 annotated metabolites were retained for downstream analysis.

### Proteomics Analyses

Plasma proteins were characterized using a TripleTOF 6600 system (Sciex) via liquid chromatography-mass spectrometry (LC-MS) with SWATH acquisition, following the methodology outlined in a previous study^70^. In every injection, 8-µg of tryptic peptides, derived from undepleted plasma, were loaded onto a ChromXP C18 column (0.3 × 150 mm, 3 μm, 120 Å, Sciex). The separation of peptides was achieved through a 43-minute gradient ranging from 4% to 32% B. High sensitivity MS/MS mode was utilized to construct variable Q1 window SWATH Acquisition methods (100 windows) with Analyst TF Software (v1.7). Scoring of peak groups was performed with PyProphet (v2.0.1)^251^ and alignment of peak groups with TRIC^252^, each adhering to stringent confidence thresholds (1% FDR at peptide level; 10% FDR at protein level). The abundance of proteins was calculated as the cumulative sum of the three most abundant peptides.

### Luminex Multiplex Assays for Targeted Cytokine, Chemokine, and Growth Factors

The evaluation of circulating cytokines, chemokines, and growth factors was undertaken employing established procedures from the Stanford Human Immune Monitoring Center (HIMC). Specifically, EDTA-plasma was scrutinized using a Human 62-plex Luminex multiplex assay, consisting of conjugated antibodies (Affymetrix, Santa Clara, California). The raw data obtained from the assay were normalized against the median fluorescence intensity (MFI) value. Subsequently, variance stabilizing transformation (VST) was applied to the data to eradicate the batch effect, adhering to our previously outlined methodology^145^. Measurements featuring background noise (CHEX) exceeding five standard deviations from the mean (mean ± 5 × SD) were omitted from the data.

### Exposome and Associated Environmental Analyses

As previously outlined^90,253^, the process of data collection for the exposome and associated environmental elements proceeded accordingly. The chemical exposome was sampled using the RTI MicroPEM V3.2 personal exposure monitor (RTI International, Research Triangle Park, NC, USA) for two participants. The MicroPEM, an active air sampling apparatus, operates by circulating air at a rate of 0.5 L/min. It was modified to house a customized cartridge containing 200 mg of zeolite adsorbent beads (Sigma 2-0304, Sigma-Aldrich Corp., St. Louis, MO USA) positioned at the airflow’s termination to gather both hydrophobic and hydrophilic compounds. Each sampling session spanned approximately five days. Post-session, the cartridge was detached and preserved at -80 L until subsequent processing. Chemical extraction involved the resuspension of zeolite beads in a sterile Eppendorf LoBind tube filled with 1 mL of Mass Spec grade methanol. Following a 20-minute incubation period at room temperature, the samples were subjected to a 20-minute centrifugation process at 22,000 g, also at room temperature. The samples were then analyzed using a Waters UPLC-coupled Exactive Orbitrap Mass Spectrometer (Thermo, Waltham, MA, USA), yielding a collection of 158 exposome chemicals. Environmental data, on the other hand, were sourced from several origins. Parameters such as temperature, humidity, and sampling flow rate were directly recorded by the MicroPEM. GPS coordinates of the participants provided geographical data. Additional meteorological and demographic information was obtained from public data repositories including the Climate Data Online (CDO), the US Census Bureau, and local weather stations. This culminated in the collection of 10 environmental feature data points.

### Dietary Analyses

A total of 25 food items were included in a questionnaire that was completed voluntarily by participants during their routine visit in the study, using a diet questionnaire hosted on https://www.projectredcap.org/ from our previous report^145^. The frequency of consuming each food item was scored in downstream analysis. For breads, biscuits, cakes, pies and pastries, we score the frequency from 0∼4, with the associated frequency per day: 0) less than 1, 1)1 per day, 2)2∼3 per day, 3)4∼5 per day, 4)6 or more. For other foods we ask the participants to state the frequency from 6+ times per day to less than once per month.

### Insulin-Suppression Test

A subset of eligible consenting participants (N=58) underwent an Insulin-suppression Test (IST), as a measure of insulin-mediated glucose uptake, to evaluate the insulin sensitivity status. Following a 12-hour overnight fast, participants were administered an infusion comprising 0.27 ug/m2 min of octreotide, 25m U/m2 min of insulin, and 240 mg/m2 min of glucose over a three-hour period during their visit to Stanford’s Clinical and Translational Research Unit (CTRU). Blood samples were procured at ten-minute intervals during the final half-hour of the infusion, resulting in a total of four blood draws. These samples were analyzed to determine plasma glucose and insulin levels. The mean value of the four steady-state plasma glucose (SSPG) and insulin concentrations were subsequently calculated. Participants were then categorized based on their SSPG values^254^: those with SSPG < 150 mg/dl were classified as insulin sensitive (IS) (n=28), while those with SSPG ≥ 150 mg/dl were classified as insulin resistant (IR) (n =30). Participants who were unable to provide measurements due to personal or medical circumstances were allocated to an indeterminate group (Unknown) (n= 28).

### Clinical Lab Test

Clinical lab tests were performed at the Stanford Clinical Lab following its guideline of blood and urine collection and submission (https://stanfordlab.com/test-directory.html). The test includes a metabolic panel, complete blood count panel, glucose, HbA1C, insulin measurements, hsCRP, IgM, lipid panel, kidney panel, liver panel. Detailed measurements and annotations are provided as a supplementary table. (**Table S1**7)

### UMAP for Microbiome Distribution

The distribution of the microbiome was visualized using Uniform Manifold Approximation and Projection (UMAP), facilitated by the R package “Seurat (Version 4.0)”. Prior to the application of UMAP, the count data were first normalized to relative abundance and scaled to represent one million reads per sample. A total of 1524 variable features were identified and encapsulated within a Seurat object. From this point, a distance matrix was produced through the utilization of the R package “Vegan (Version 2.6-2)”, employing Bray Curtis dissimilarity. This distance matrix was subsequently transformed via Principal Coordinate Analysis (PCoA). The first ten dimensions from the PCoA (out of a total of 1094 generated) were used to determine neighboring relationships. The UMAP projection was then calculated, with default settings employed. This projection was made possible by invoking Python’s UMAP via reticulate. The final UMAP results were visualized using the first two dimensions.

### Intraclass Correlation

Intraclass correlation Coefficient (ICC) was calculated from Linear Mixed Models, in which we modeled random intercepts but a fixed slope, allowing different personal levels between individuals^70^. We first linearly transformed each analyte (when applicable) and standardized the total variation to 1 before applying ‘lmer’ function from R package “lme4 (V1.1-30)”, with the formula as:

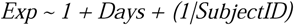

Where *Exp* was the linearly transformed and standardized values of each analyte, *Days* was the length of time individuals participated in the study, *SubjectID* was the subject ID associated with each participant.

We then used ICC as the proportion of total variation explained by subject structure in the cohort by V_subjectID_/V_total_, in which V was the variance from the corresponding component extracted by Var_Corr_ and V_total_ was 1.

### Permutation Test on Bray Curtis Distance Comparison

The Bray-Curtis (BC) distance was used to quantify the degree of similarity between two microbiome samples, with the ASV serving as the unit for calculating dissimilarity for the complete microbiome sample or for specific taxa. Similarity metrics were calculated pairwise for both intra-individual and inter-individual comparisons. A permutation test was employed to estimate the null distribution while accounting for the varying sample sizes of each participant. The null hypothesis being tested was that there is no difference between intra-individual and inter-individual distances.

For each microbial genus, a test statistic was calculated as the mean difference in BC distances between intra-individual and inter-individual comparisons. To estimate the null distribution, all sample labels were randomly permuted, and the BC distances were computed pairwise. This process was repeated 10,000 times, generating a null distribution of test statistics.

P-values were then calculated by determining the proportion of permuted test statistics that were at least as extreme as the observed test statistic. In the case of multiple comparisons, such as for different microbial genera, p-values were adjusted using the BH procedure to control the false discovery rate. Statistical significance was determined using a threshold of BH adjusted p-value < 0.1.

### Degree of Microbial Individuality

Degree of Microbial Individuality (DMI) was measured as the mathematical difference between a given genus regarding their populational median of the inter-individual BC distance and the median of the intra-individual BC distance. To assess the robustness and variability of the DMI estimates, we employed a bootstrap resampling technique^255^.

First, we summarized the between-sample distance of genera with longitudinal prevalence greater than 10% at a given body site. The distance of sample pairs was then allocated into inter-individual or intra-individual groups. For each genus, we resampled the data with replacement and computed the DMI using the following formula:

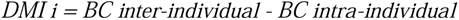

This resampling process was repeated multiple times to obtain a distribution of DMI estimates, which enabled us to assess the variability and robustness of our results.

By employing the bootstrap resampling technique, we aimed to gain insights into the reliability of our DMI calculations and understand how sensitive they were to potential variations in the data. This approach provided a more comprehensive understanding of the degree of microbial individuality across various body sites and taxa. To ensure the accuracy of our results, we considered a DMI to be reliable only if that DMI’s standard deviation (SD) of the bootstrapped distribution is less than 15 times of the mean DMI. This criterion helped to filter out any DMI estimates that might be influenced by high variability or uncertainty in the data. Detail of the bootstrap result was included in our supplementary materials (**Table S7**), where we used column “confidence” to distinguish DMI values with high or low reliability. The total number of genera reported per region after bootstraping included: 105 for stool, 35 for skin, 63 for oral, and 33 for nasal samples.

To quantify the cumulative DMI for each individual, the DMI score was multiplied by the average relative abundance of each genus for a given individual. This generated a weighted DMI (abundance_dmi) that represented the product of the DMI score and the genus’s relative abundance. The total DMI for each individual was computed by summing these weighted DMI values across all genera. This approach offered a comprehensive measure of the overall DMI per individual, accounting for the contribution of each genus weighted by its relative abundance in the individual’s microbiome.

### Family Score

To assess the impact of a shared living environment on microbiome variability, we introduced a metric called the Family Score (FS). The FS represents the relative influence of a shared environment on the inter-individual dissimilarity of a given genus within cohabitating pairs. We began by excluding genera with a longitudinal prevalence of less than 10%, leaving 141 for stool, 119 for skin, 41 for oral, and 33 for nasal samples. For each remaining genus, we computed the “within-family” inter-individual Bray-Curtis (BC) distance between cohabiting pairs. We also calculated the median inter-individual BC distance (BC inter-individual) and intra-individual BC distance (BC intra-individual). Using these values, we computed the FS for a given genus ‘*i*’ with the following formula:

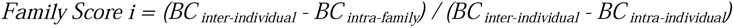

This formula normalizes the FS to a scale that allows for comparisons across families and non-families. An FS of 0 indicates that the shared living environment has no impact on inter-individual dissimilarity, while an FS of 1 suggests that living in the same environment causes inter-individual dissimilarity to resemble intra-individual dissimilarity. We truncated the FS at 0 and 1 to maintain consistency in the comparison scale. Any FS values greater than or equal to 1 were assigned a value of 1, and values less than or equal to 0 were assigned a value of 0.

### Classification of The Microbiome Genera by Their Longitudinal Prevalence

The microbiome genera were categorized as the core microbiome, opportunistic microbiome, and middle group based on their longitudinal prevalence^256–258^. Calculation of prevalence was based on the presence or absence of reads from each sample. For each sample, the relative abundance of each genus was first transformed to 1 if it was greater than 0; then, the proportion of 1 for each genus in each participant was determined as the longitudinal prevalence. Then the genera were assigned to a group based on their longitudinal prevalence: core microbiome: longitudinal prevalence > 80%; middle group: 20% ≤ longitudinal prevalence ≤ 80%; opportunistic microbiome: longitudinal prevalence < 20%.

### Body Site-Specific Longitudinal Model for Bray-Curtis distance

To estimate the effect of body site on the change in BC distance over time, we used a linear mixed effects regression with the BC distance as the response variable implemented in the “lme4” package in R^259^. The BC distance of all pairwise samples was transformed to a more normally distributed dataset. The transformation method used was the log-log transformation, which has been widely used in survival analysis studies. Specifically, the transformation was performed by applying the formula:

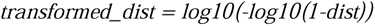

A total of eight different transformations were evaluated (variations of logarithmic, square root, arcsine, and ratio transformations of the original distance) to identify the most suitable transformation for our analysis. To assess the normality of the transformed distances, an Anderson-Darling test was performed, and the best method was selected by Anderson-Darling’s statistics A^260^.

Fixed effects included an interaction between the body site and time, and with a random intercept for each individual. Time was normalized to the days from the first sample for each individual. The model was optimized using the “nlopwrap” method^261^ and nested models were compared by likelihood ratio test using the “lmtest” package in R^262^. Differences in the slopes of the body sites over time were assessed by F-test with Satterthwaites’s degrees of freedom^263^. For the comparisons between body sites, the model was constructed as:

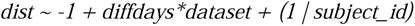

For the comparison between IR and IS participants, the model was constructed as:

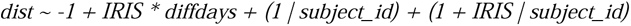

Where *dist* was the pairwise BC distance, *diffdays* was the date interval between two samples, *subject_id* was the subject ID associated with each participant, and *IRIS* was the insulin sensitivity status of each participant.

### Bayesian Mixed-Effects Model for Microbial Taxa and Cytokine Interactions

We utilized a Bayesian negative-binomial longitudinal mixed-effects model to analyze the interactions between microbial taxa and cytokines. This model choice addresses several key characteristics of our data: the compositional nature of microbiome data, the presence of zero-inflation and high skewness in cytokine measurements, and the repeated measurements of both cytokine and microbiome data in our study cohort (n=62 with longitudinal, date-matching measurements). The model’s capacity to handle the highly skewed cytokine data was a critical factor in its selection, especially given the dramatic range of cytokine surges observed in our dataset, ranging from a 10.8-fold increase for MIP1B to as much as a 618-fold increase for LEPTIN. Each genus’s reads count was modeled as a sparse-matrix response variable, with a plasma cytokine level MFI quantity and time as fixed effects and a random intercept for each individual, following the formula:

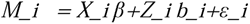

Where *Mi* is a vector of the genus-level microbe relative abundances for each participant *i*, *Xi* is the design matrix for the fixed effects, β. Each row of matrix *Xi* contains the terms (1) time (days post–study start), *Di*, and (2) cytokine measurements, *Yi*, from 1 to n. *Zi* is the random effects design vector of 1’s denoting a random intercept, *bi* is a scalar for each participant, and ε*i* is a zero-centered error term.

Posterior sampling was performed using four chains, 5,000 iterations per sample, and a 1,000-iteration burn-in of a No-U-Turn Sampler implemented in the “brms (Version 2.18.0)” package in R^264–267^. The iteration plots and posterior predictive distributions were visually inspected for chain convergence. Microbe genera and cytokines above the limit of detection in less than 10% of the samples were excluded from the analysis. A microbe-cytokine association was considered significant if the 95% credible interval on the fit coefficient of the cytokine term did not include zero.

In our model, the beta represents the effect size. We estimated the variance explained by the model using the “bayes_R2” function from the brms package, and this information is included in the final table. While we provide p-values for each pair, as inferred from a method detailed previously^268^, and the subsequent BH-adjusted p-values, it’s important to note that these are supplementary. In our Bayesian analysis, the determination of significance is based on the commonly practiced use of credible intervals derived from Markov Chain Monte Carlo (MCMC) sampling.

### Correlation Network Analysis

Correlation network analysis was conducted to construct a network between the microbiome (from stool, skin, oral, and nasal samples) and internal multi-omics data (proteome, metabolome, and lipidome) from plasma, following a modified version of published methods^70^. Initially, time points with unmatched collection dates for each pair of microbiome and internal omics data were excluded. Subjects with fewer than five samples for a specific microbiome type were also removed from the corresponding correlation analysis.

The microbiome data, which included relative abundance or observed ASV richness at the genus level, was processed by retaining only genera detected in at least 10% of all samples. Centered log ratio (CLR) normalization was applied to address compositionality in microbiome data using the R package “compositions” (Version 2.0-4). Proteome, metabolome, and lipidome module data were log_2_ transformed.

To account for repeated sampling from the same subject, a linear mixed-effects model was utilized, incorporating subject ID as a random effect using the R package “lme4” (Version 1.1-30). Spearman correlations were then calculated, and p-values were adjusted using the Benjamini & Hochberg method. For correlation network construction, BH adjusted p-values < 0.2 were included. In the linear mixed model assessing longitudinal effects, the estimated correlation coefficient served as the measure of effect size (**Fig. S26**).

To gauge the robustness of the point estimation for the proportion of microbiome genera that are significantly correlated, we employed a bootstrapping approach. Specifically, we determined the percentage of significantly correlated genera by dividing the number of significantly correlated pairs by the total possible pairs. We bootstrapped this procedure 20 times to obtain mean and standard deviation values for the percentage of significantly correlated microbiome genera. These values were then used to compare the scale of interdependency of microbiome across different body sites. To ascertain if one body site exhibits a significantly greater interdependency compared to another, we employed the two-sided Wilcoxon rank-sum test. The significance of these comparisons was determined using p-values adjusted for multiple testing using the BH correction.

The estimation of inter-individual microbiome-wide correlation was conducted utilizing the SparCC method (R package discordant, Version 1.25.0), a methodology specifically tailored for compositional data^269,270^. To prepare the microbiome relative abundance data for this analysis, the data were multiplied by 20000 and subsequently rounded, a process designed to convert proportions into data resembling counts. Following this, the SparCC method was employed to compute the correlation matrix. This matrix was then transformed into a long format data frame, with each row representing a pair of genera and their corresponding correlation coefficient. To assess the statistical significance of the observed correlations, a permutation test was implemented. For each pair of genera, the sample labels were randomly permuted, and the permuted SparCC correlation coefficient was calculated. We executed this procedure 10,000 times to produce a null distribution of SparCC correlation coefficients. Following this, we counted the instances where our observed correlation coefficient was encompassed within this null distribution. A finding was deemed “non-significant” if the correlation coefficient from any of the 10,000 permutations was more extreme than the coefficient from our original analysis, and subsequent raw p-value was computed based on the permutation results. Results with the BH adjusted p-value < 0.1 from this permutation was included in our final report.

The inter-omics correlation network was visualized using the R packages “ggraph” (Version 2.0.5), “igraph” (Version 1.3.2), and “tidygraph” (Version 1.2.1) under the “kk” layout.

### Pathway Enrichment for Proteins Correlated with the Microbiome

The enrichment of pathways corresponding to proteins associated with the microbiome from four different body sites was achieved via the R package “clusterProfiler (Version 3.15)”. The proteins in question served as input for pathway enrichment, specifically through Gene Ontology (GO) processes, enabling the identification of statistically significant pathways. The determination of these pathways was reliant on Fisher’s exact test.

GO terms of interest were isolated based on their Benjamini-Hochberg adjusted p-values; terms with p-values below 0.05 were retained for ensuing analyses. For the GO terms that demonstrated significant enrichment, the R package “simplifyEnrichment” was employed. This package facilitated the calculation of similarities between each pair of GO terms. The construction of a network only incorporated edges that exhibited similarities exceeding 0.70.

In order to discern modules within the correlation network, community analysis was undertaken utilizing the R package “igraph (Version 1.3.4)”. To encapsulate each module, only the GO term that presented with the smallest Benjamini-Hochberg adjusted p-value was preserved.

### Mediation Analysis

A mediation analysis was conducted to investigate the potential influence of microbiomes from stool, skin, oral, and nasal sources on phenotypes through internal multi-omics data, including proteome, metabolome, lipidome, and cytokine^56,271,272^. Phenotype data were obtained via clinical laboratory tests of plasma samples. The associations between the microbiome and phenotype (Direct Effect), as well as internal omics data (Indirect Effect), were initially determined as outlined in the ‘Correlation network analysis’ section. Only significant associations (BH adjusted p-values < 0.2) were considered for subsequent mediation analysis. The linear regression model from R package “mediation” was employed for the mediation analysis. Ultimately, pairs with significant Average Causal Mediation Effects (ACME, p-values < 0.05) were reported, representing the microbiome’s impact on phenotype measurements through internal multi-omics.

To control the false discovery rate (FDR), a reverse mediation analysis was performed by exchanging the mediator with effects (i.e., the microbiome influencing internal omics data via phenotype), and pairs with significant ACME in reverse mediation (p-values < 0.05) were excluded from the final results. Comparisons between body sites and insulin sensitivity statuses were conducted for each dataset using the Fisher exact test. Initially, it was hypothesized that the true meditative effect of each mediated pathway (e.g., genus *i* ∼ cytokine *j* ∼ immune *k*) was considerably large (n= 1 × 10^8^). Subsequently, the presence of a significantly different meditative linkage between the designated comparison pairs was assessed.

### Principal Variance Component Analysis (PVCA)

To assess the variation in microbiome data based on individual and season, the principal variance component analysis (PVCA)^273^ was performed via R package “pvca (Version 3.15)”. The PVCA is a combination of the principal component analysis and variance components analysis https://www.niehs.nih.gov/research/resources/software/biostatistics/pvca/index.cfm, which were originally employed to assess batch effects in microarray data^274^ and widely used for microbiome related variance decompositions^275,276^. For microbiome samples in each body site, the season was determined by subtracting the date of collection from the first day of the year (from 1-365 days). Each sample’s participant ID and season were then entered into the PVCA as variables. Then, the “ggtern (Version 3.3.5)” R package was used to visualize the data.

### Deconvolute the Environmental Effect on the Microbiome

#### Exposome and Diet Data Analysis

To investigate the influence of exposome and diet data on the microbiome from different body sites, exposome data (chemical and environmental) were collected and processed as previously described^253^. Diet data were collected and detailed in the methods section above. As an example, the analysis process for exposome chemical data is described below.

Microbiome samples with matching exposome chemical data within a 3-day period were selected for subsequent analysis (N_Chemical_ = 8, N_Environmental_ = 32 (N_Participant1_ = 13; N_Participant2_ = 19)). Microbiome data were normalized using the centered log ratio (CLR, “clr” function from R package “compositions”), and exposome data were log2-transformed and auto-scaled^92^. Principal component analysis (PCA) was performed on both microbiome and exposome data. Principal components (PCs) from the microbiome and exposome were further analyzed, with PCs accounting for over 80% of cumulative explained variation being included. A linear regression model was constructed using PCs of microbiome data as the dependent variable (Y) and corresponding exposome PCs as the independent variable (X). The R2 value was extracted to represent the exposome’s contribution to microbiome data. The same method was applied to evaluate the dietary effect on the microbiome from four body sites.

#### Seasonal Effects on Microbiome

Z-score normalized microbial data or microbial diversity was systematically analyzed using Generalized Additive Mixed Models (GAMMs)^277^. For each genus of interest, the model was formulated as:

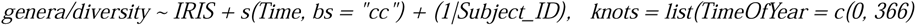

In this model, the response variable *’genera’* represents the z-score normalized microbial relative abundance. The fixed effects components include the status of insulin sensitivity (*IRIS*) and a cyclic cubic spline smoother for the Time variable, encapsulating potential cyclical patterns across the year (from 0 to 366). The term (*1|Subject_ID*) includes a random intercept for each subject, to account for within-subject correlation.

The model was fitted using the Restricted Maximum Likelihood (REML) method for robust estimation of smoothing parameters in a complex and unbalanced design and incorporated the use of ‘lmeControl’ function from the ‘nlme’ package in R to handle the optimization process of the mixed-effects models. This was conducted by specifying ‘optim’ as the optimizer for the model fit.

The resulting model provides insight into the temporal dynamics of gene expression and its relationship with insulin sensitivity status (IRIS), considering the random effects associated with each subject. The graphical representation of these models for each genus and p value for smooth terms were saved for further exploration.

#### The Effects of Infection on Microbiome

The analysis of the effects of infection on the microbiome involved genera that were significantly altered during infection periods. The methodology implemented was informed by a previous method for estimating infection processes based on self-reported symptoms^70^.

The infection status was classified into longitudinal categories: pre-healthy (-H) state, event early (EE) state, event late (EL) state, recovery (RE) state, and post-healthy (+H) state. The pre-healthy state comprised the healthy baselines observed within 186 days preceding the onset of the infection event. The EEs state was characterized by visits occurring between day 1 and day 6 of the event. The EL state spanned visits on days 7 to 14 since the onset of the event. The recovery state included visits within the 15–40-day period since the event’s inception, and the post-healthy state encompassed visits within the 186 days following the event.

The categorization of these states was designed as a continuous progression from the pre-healthy to post-healthy state, with the average duration of an infection event being 88 days. 58 infection events were detected among 32 participants (IS: 7 participants, IR:12 participants, Unknown:13 participants) in our study. Each event was assigned a unique identification consisting of the subject ID and the event number. The progression of infection states within an event was tracked in the order of pre-healthy, event early, event late, recovery, and post-healthy.

In order to assess the effects of these infection states and insulin sensitivity status on microbiome genera, GAMM was constructed:

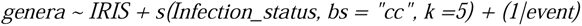

In this model, *’genera’* represents the z-score normalized microbial relative abundance, IRIS indicates the insulin sensitivity status of each participant, and *’Infection_status’* is a smoothing function of the longitudinal infection states with cyclic cubic regression splines. The term ‘(*1|event*)’ is a random intercept for each infection event.

For the evenness changes during infection among insulin resistant (IR) and insulin sensitive (IS) individuals, the model was reformulated as follows:

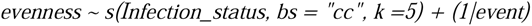

In this adjusted model, ‘evenness’ denotes the outcome variable, specifically the Subject ID based Z-score transformed Pielou’s evenness measure of the microbiome sample, indicating the diversity of the microbial community. The remaining components of the model and their interpretations align with the previously described model.

## Author Contribution

X. Zhou, X.S., J.S.J, D.J.S., G.W., M.P.S. conceptualized the study and devised the analysis plan. X. Zhou, W.Z., M. Agnello, S.R.L., M. Avina oversaw the comprehensive sample collection and conducted experimental benchwork. X. Zhou, W.Z., S.R.L., Y.Z., L.C., L.L., and E.J.B. were responsible for microbiome data acquisition and processing. D.H. and S.W. handled lipidome data acquisition and processing. C.J., Liuyiqi J., X. Zhang, X.S. undertook exposome data acquisition and processing. X. Zhou, W.Z., P.K. executed cytokine Luminex assay data acquisition and analysis. Lihua J., R.J. performed proteome data acquisition and analysis. K.C. and X.S. managed metabolomics data acquisition and analysis. J.S.J., D.J.S., X. Zhou, and X.S. designed and implemented longitudinal data modeling. W.Z., M. Avina, S.R.L., A.C. A.W.B. coordinated participants’ clinical visits and sample collection/inventory. X. Zhou, X.S., J.S.J., C.Z. carried out integrative omics analysis. X. Zhou and X.S. conducted the bootstrap and permutation analysis. M.P.S. and G.M.W. provided funding support and overall project supervision. X. Zhou, A.H., S. Chen provided additional data visualization. X. Zhou completed the data and code organization and publication, X. Zhou., J.S.J., D.J.S. and M.P.S. wrote the manuscript with contributions from all listed authors.

## Acknowledgment

We extend our sincere gratitude to all research participants for their dedication and involvement. Our thanks also go to Mrs. Ada Yee Ki Chen and Mrs. Lisa Stainton for their administrative support. We are grateful for the valuable insights provided by the members of Snyder Lab (Stanford University) and Weinstock Lab (The Jackson Laboratory of Genomic Medicine). Our appreciation extends to the Stanford Immune Monitoring Core and Ms. Yael Rosenberg-Hasson for their contribution to the cytokine Luminex Multiplex Assay. We also thank Dr. Victor Felix and Dr. Owen White (University of Maryland) for their assistance in uploading our raw sequencing data to the iHMP portal.

This work was funded by National Institutes of Health (NIH) Common Fund Human Microbiome Project U54_DE023789-01, NIH U54_DK102556-03, NIH R01_DK110186-05, NIH S10_OD020141-01, NIH R01 AT010232-04, NIH UL1 TR001085, NIH P30_DK116074, S10_OD023452-01. We are also sincerely grateful for the generous support from Leona M. and Harry B. Helmsley Charitable Trust (Grant No. G-2004-03820) and Innovative Medicines Accelerator (Grant No. IMA-1051) grant at Stanford University.

We acknowledge the fellowship support received by X. Zhou from the Stanford Aging and Ethnogeriatrics (SAGE) Research Center under NIH/NIA grant P30AG059307. The SAGE Center is part of the Resource Centers for Minority Aging Research (RCMAR) Program led by the National Institute on Aging (NIA) at the National Institutes of Health (NIH). We also acknowledge the fellowship support received by S.M.S.-F.R from the NIH K08 ES028825, A.W.B from 1F32DK126287 - 01A1, D.J.S from NIA-K01AG070310, and J.S.J from the Kennedy Trust for Rheumatology Research.

## Data Availability

The microbiome data are available at https://hmpdacc.org. Other omics data can be accessed through Stanford iPOP website at https://med.stanford.edu/ipop.html.

## Code Availability

The microbiome data specific to this study can be accessed at GitHub Repository (https://github.com/xzhou7/iHMP/tree/main/filename), which contains the file name of the file being uploaded (File: <u>xzhou_uploaded.files.txt</u>), and the specific file included in this study (File: <u>sampleDataALL.csv</u>). The metadata corresponding to these data files can be found at Stanford Data Repository (http://hmp2-data.stanford.edu/script.php?table=subject). These file names can be used (via *KitID* or *Barcode*) to query the microbiome .fastq file from NIH Integrative Human Microbiome Project Data Portal (https://hmpdacc.org/) and (via *<u>SampleID</u>*) to query the host multiomics data from Stanford Integrated Personal Omics Profiling Data Portal (https://med.stanford.edu/ipop.html). Alternatively, the R Objects related to this study are also available by contacting the corresponding author.

## Inclusion And Diversity Statement

Multiple authors of this paper and study participants of the research identify themselves as an underrepresented ethnic minority. One or more authors of this paper identifies themselves as part of the LGBTQ+ community. Multiple authors of this paper received support from programs that are designed to increase minority representation in science.

## Declaration of Generative AI and AI-assisted Technologies in the Writing Process

During the preparation of this work the author(s) used ChatGPT in order to improve the quality of scientific writing. After using this tool/service, the author(s) reviewed and edited the content as needed and take(s) full responsibility for the content of the publication.

