## Supplementary material for "Longitudinal profiling of the microbiome at four body sites reveals core stability and individualized dynamics during health and disease": SupplimentaryFigures

**Supplementary Figure S1. Relative Abundance of Representative Genera Displayed on Uniform Manifold Approximation and Projection**

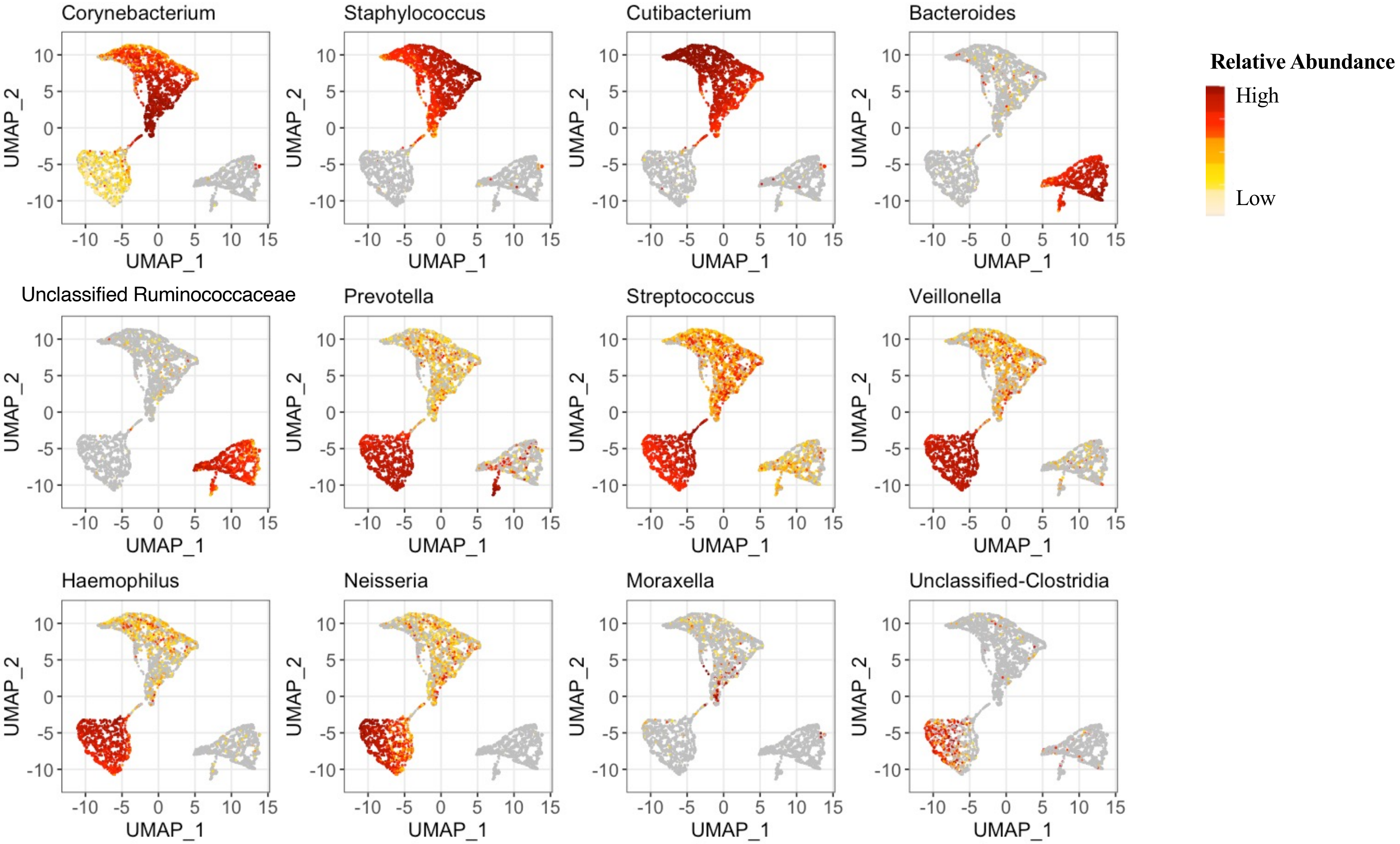

#### Supplementary Figure S2. Principal Coordinate Analysis Distribution of Samples Differing in Insulin Status

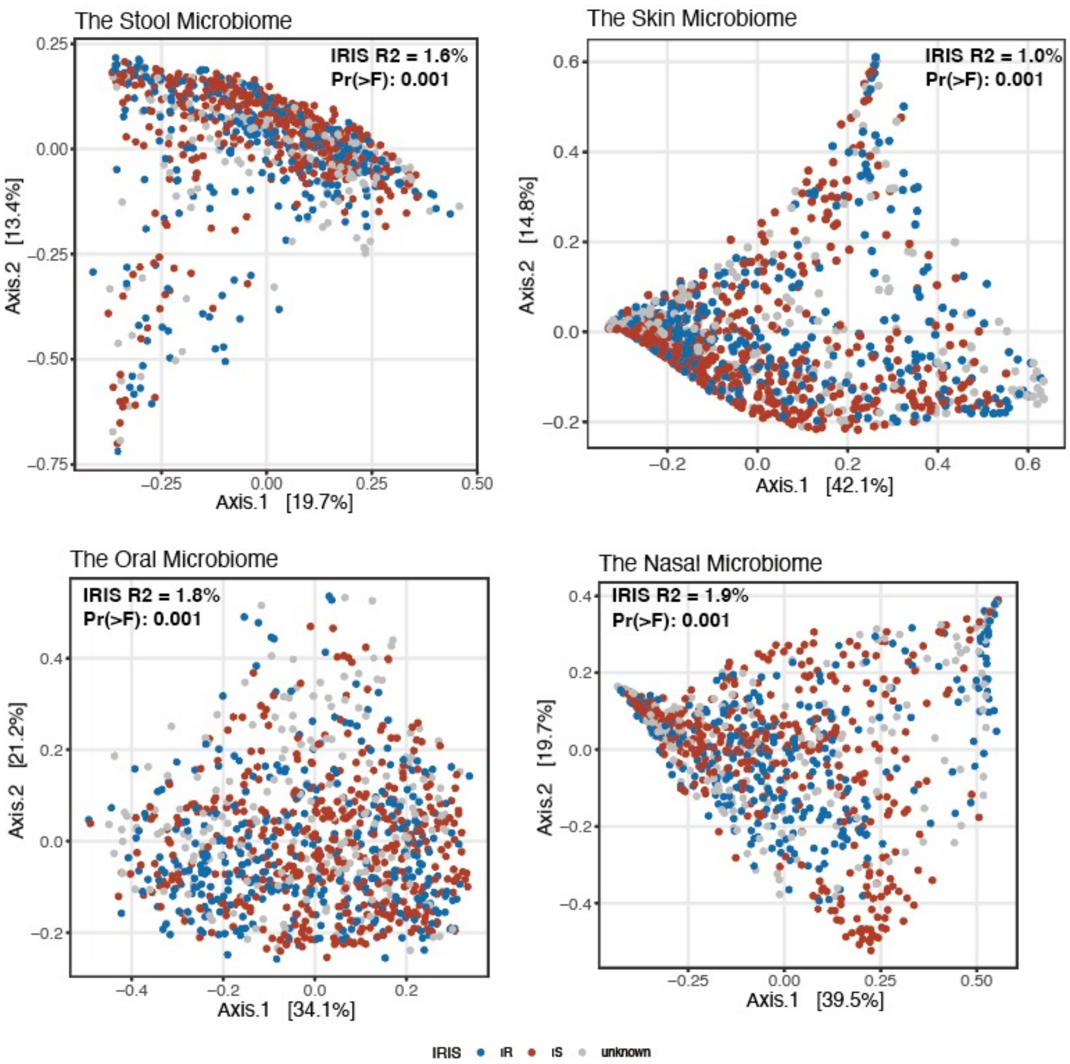

**Supplementary Figure S3. Intraclass Correlation of Microbiome at Each Taxonomy Level**

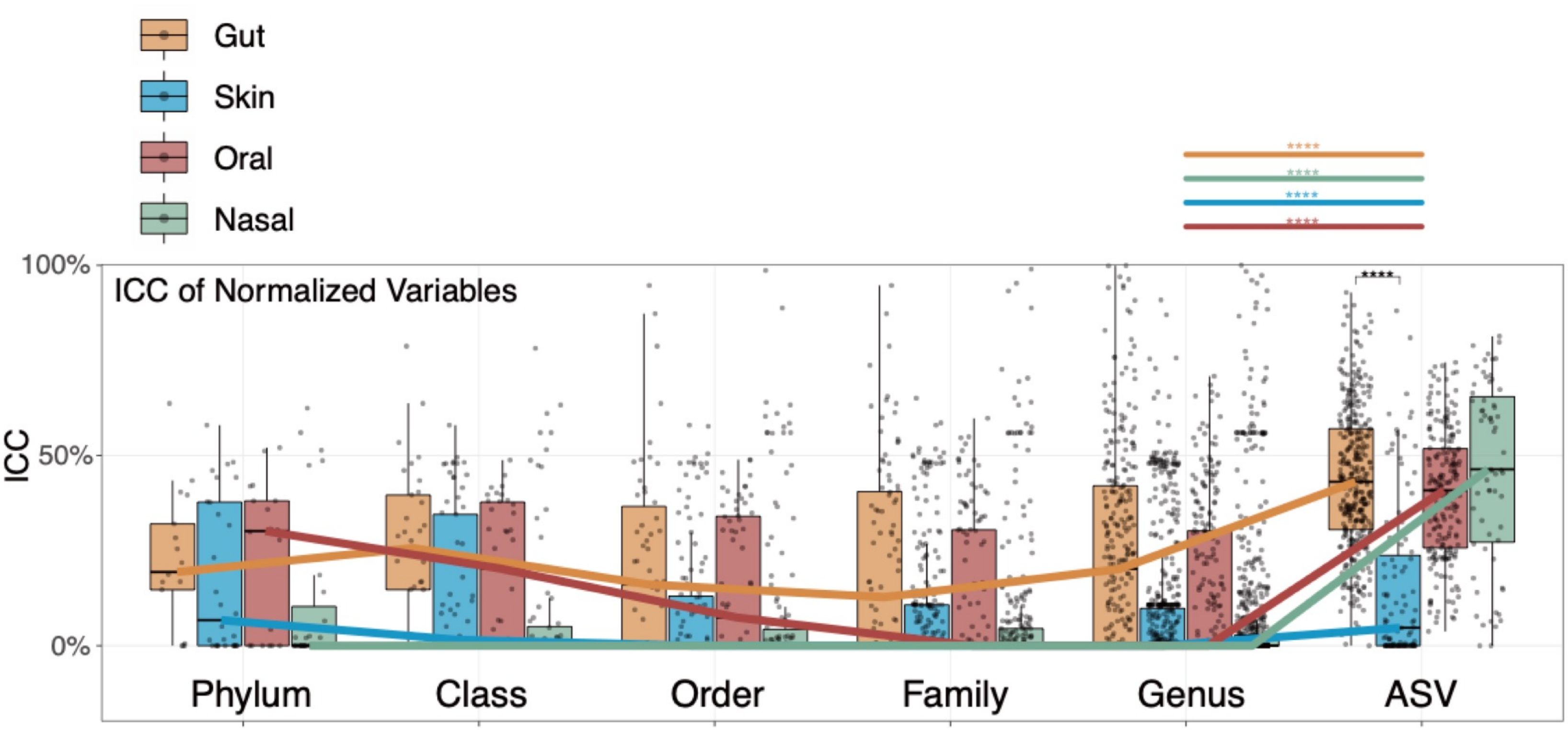

**Supplementary Figure S4. Microbiome Variance Explained by Individuality, Season, and Residuals.**

A

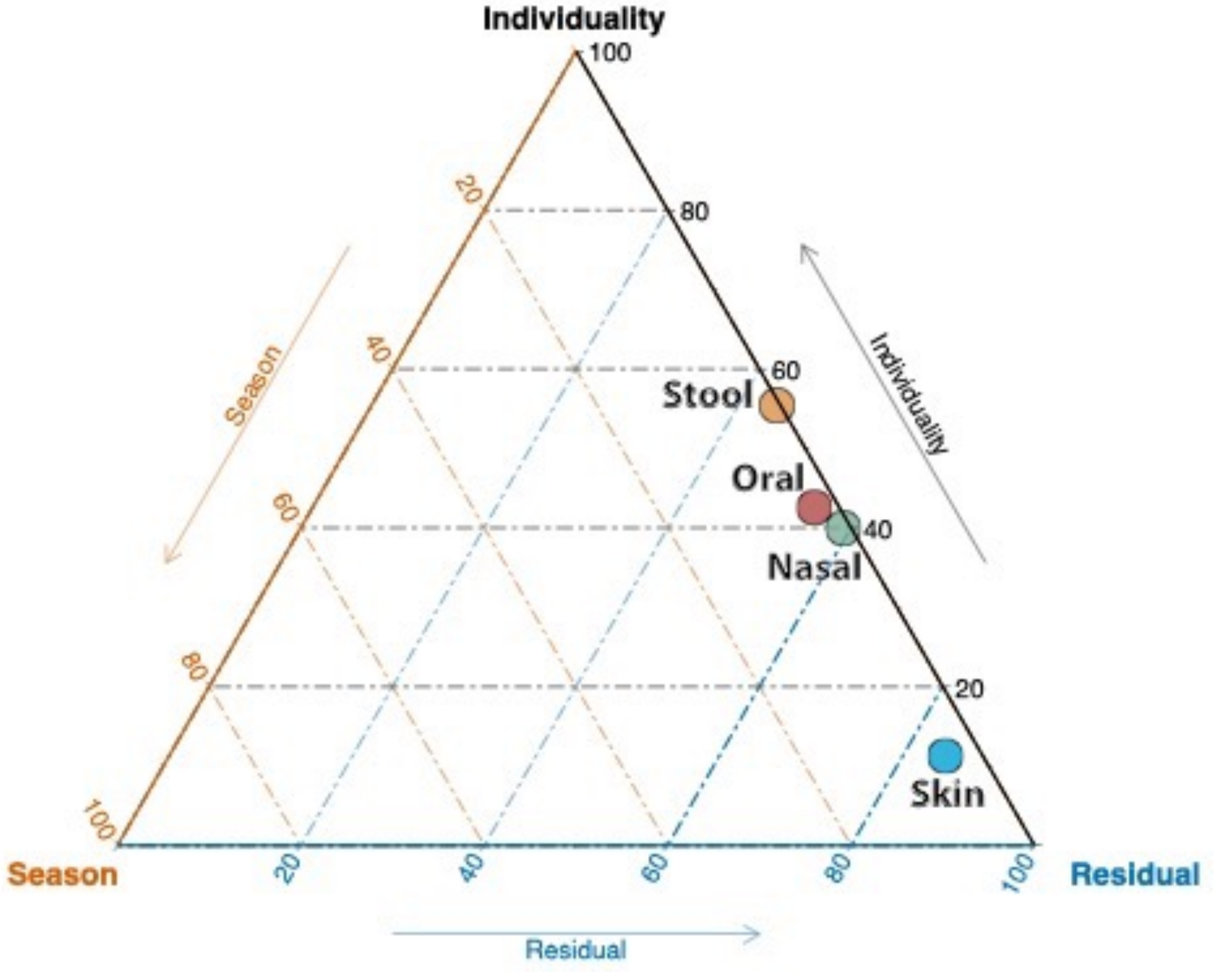

B

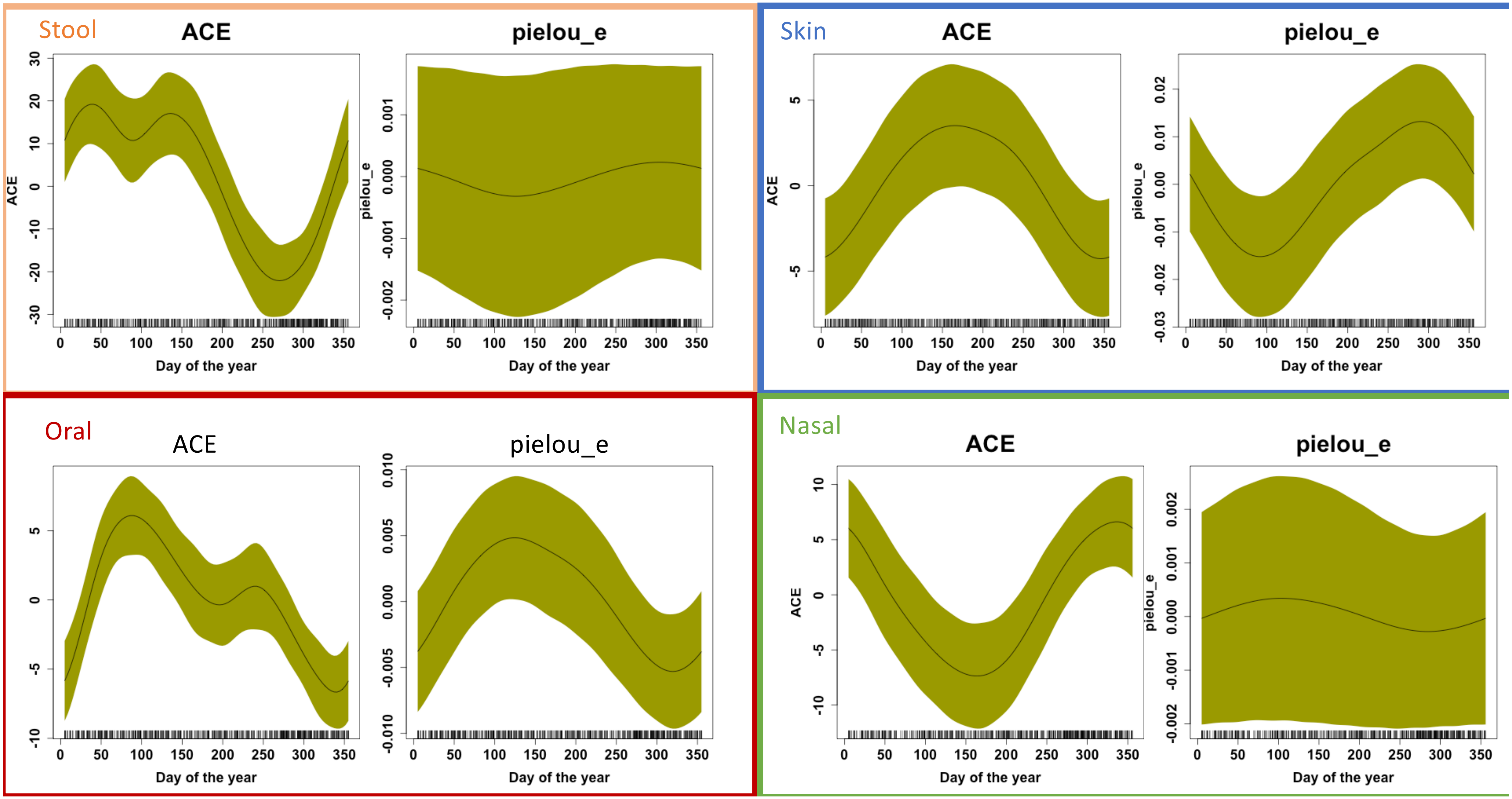
Stool
ACE
pielou_e
Skin
ACE
pielou_e
Oral
ACE
pielou_e
Nasal
ACE
pielou_e
Day of the year

**Supplementary Figure S5: Variance in Microbiome Explained by Diet and Exposome**

A

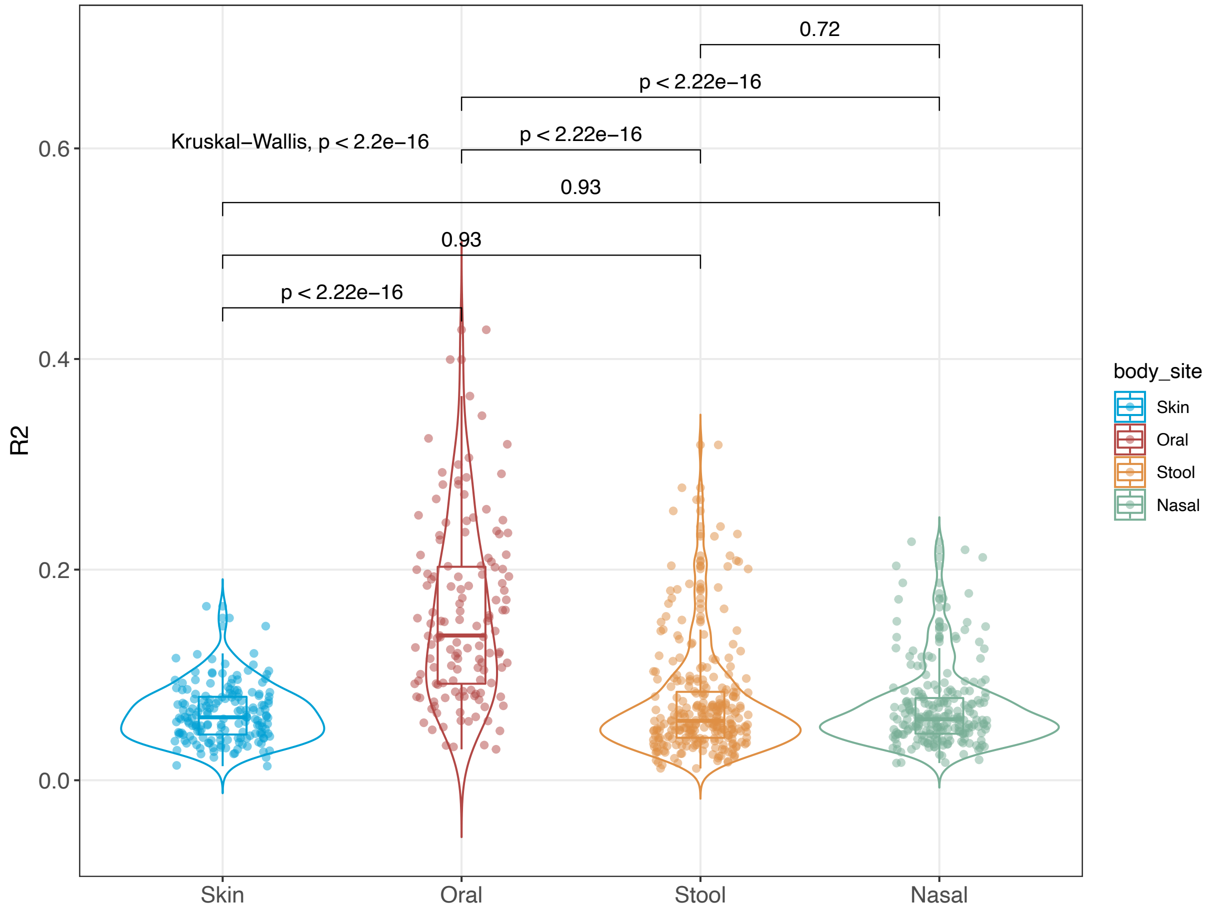

B

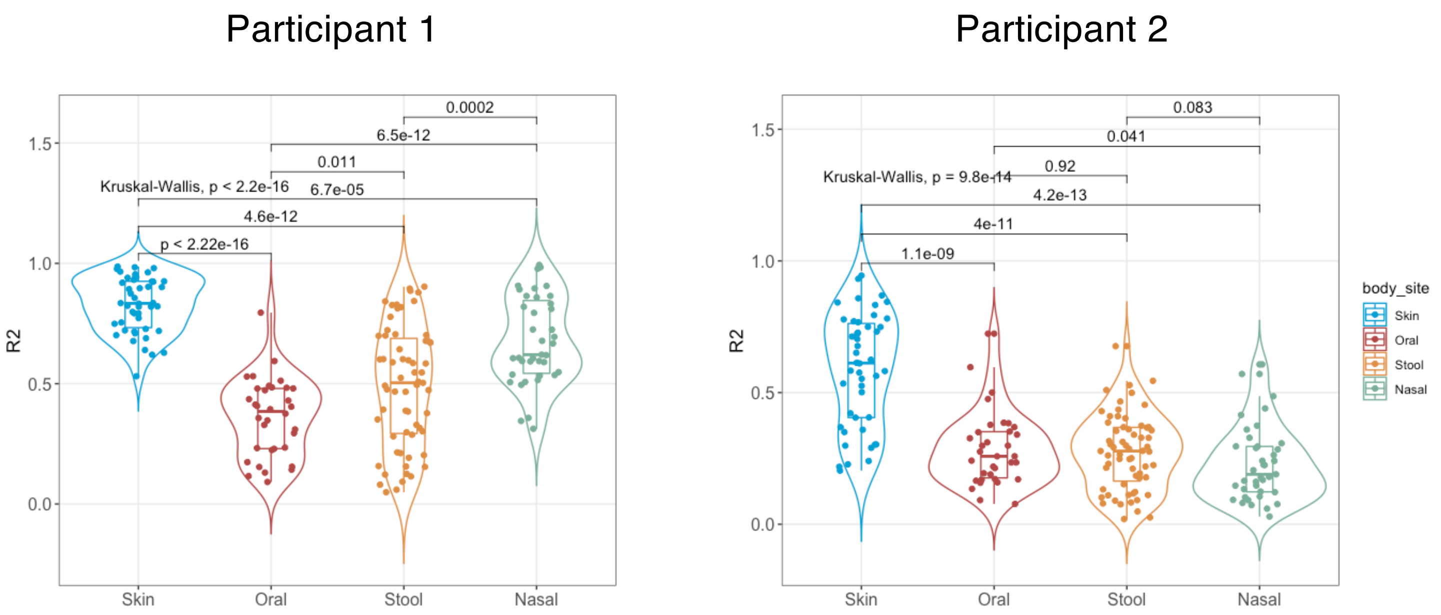
Participant 1
Kruskal-Wallis, p < 2.2e-16
p < 2.22e-16
4.6e-12
6.7e-05
0.011
6.5e-12
0.0002
R2
Skin
Oral
Stool
Nasal
Participant 2
Kruskal-Wallis, p = 9.8e-14
1.1e-09
4e-11
4.2e-13
0.92
0.041
0.083
body_site
Skin
Oral
Stool
Nasal

**Supplementary Figure S6. Shifts in Diversity and Evenness Between Insulin-Resistant and Insulin-Sensitive Individuals**

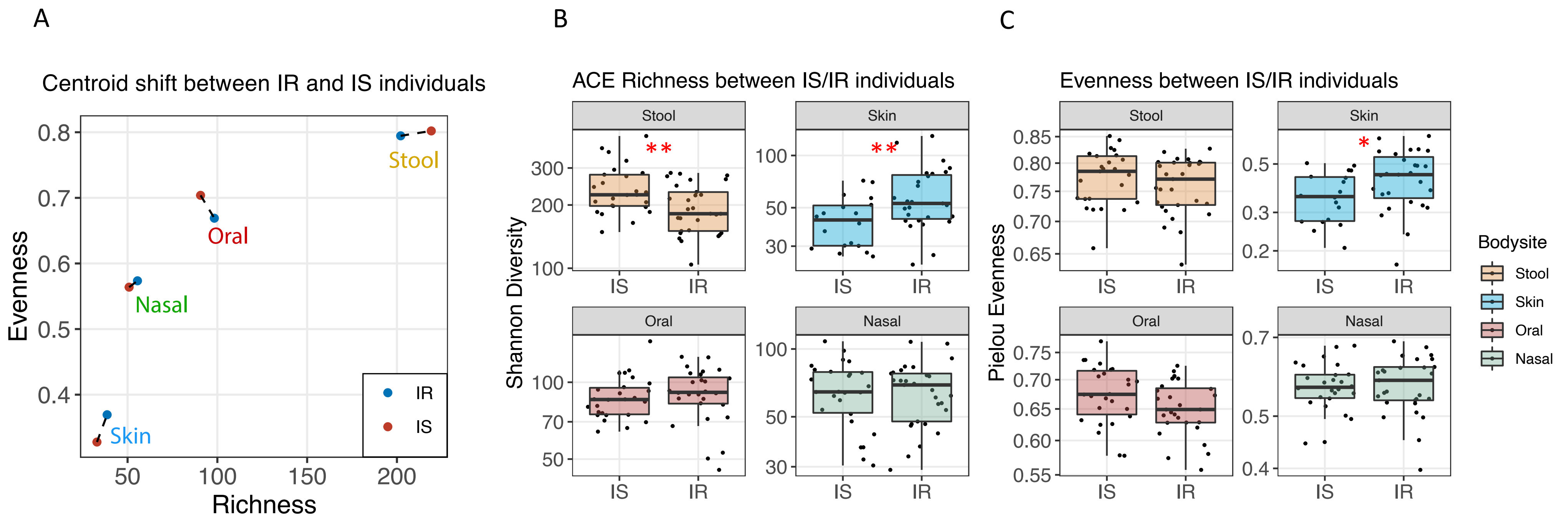

#### Supplementary Figure S7. Prevalence by Relative Abundance Plot

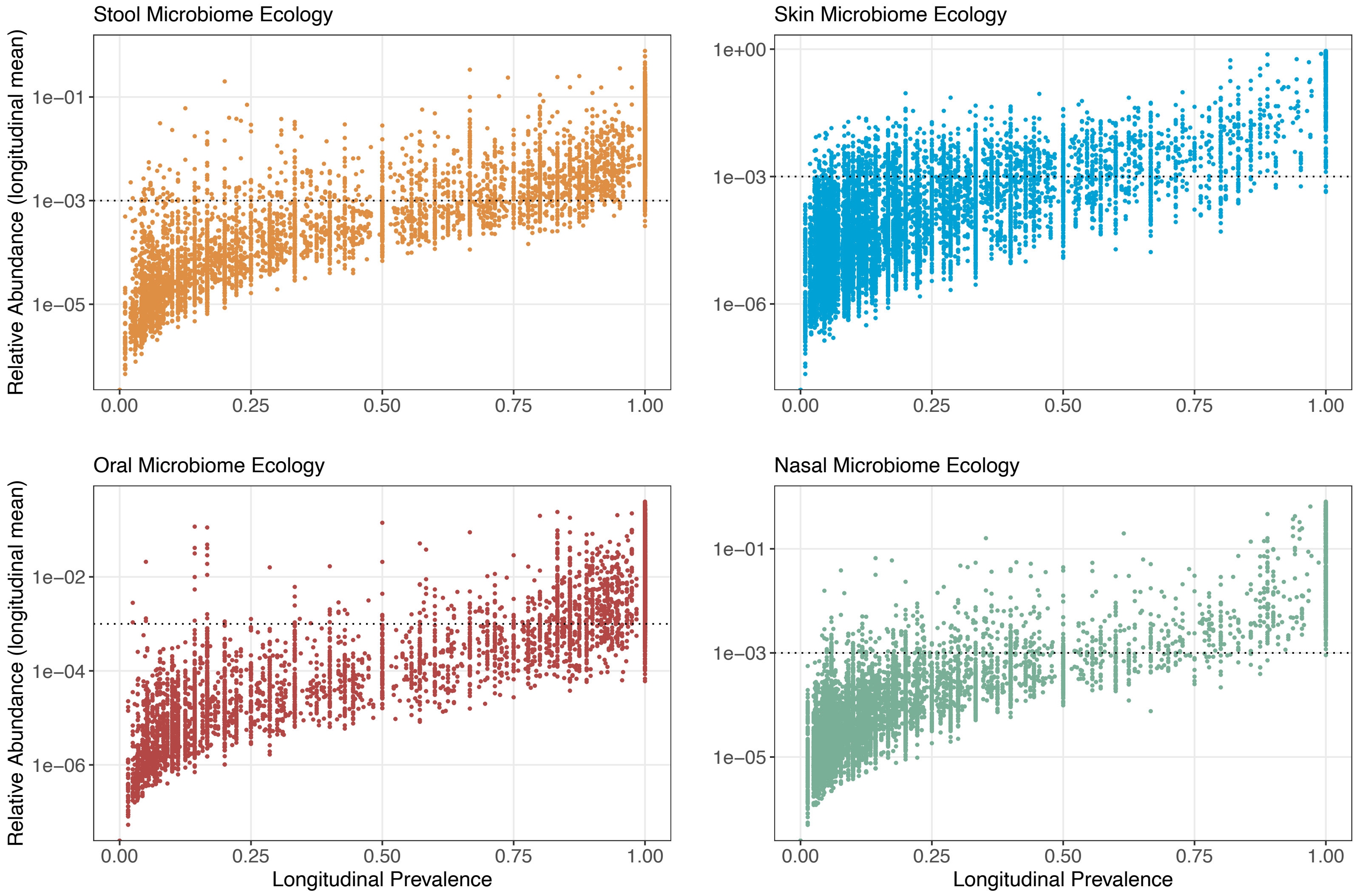

**Supplementary Figure S8. Relationship Between the Number of Core Microbiome Genera, Steady-State Plasma Glucose, and Body Mass Index**

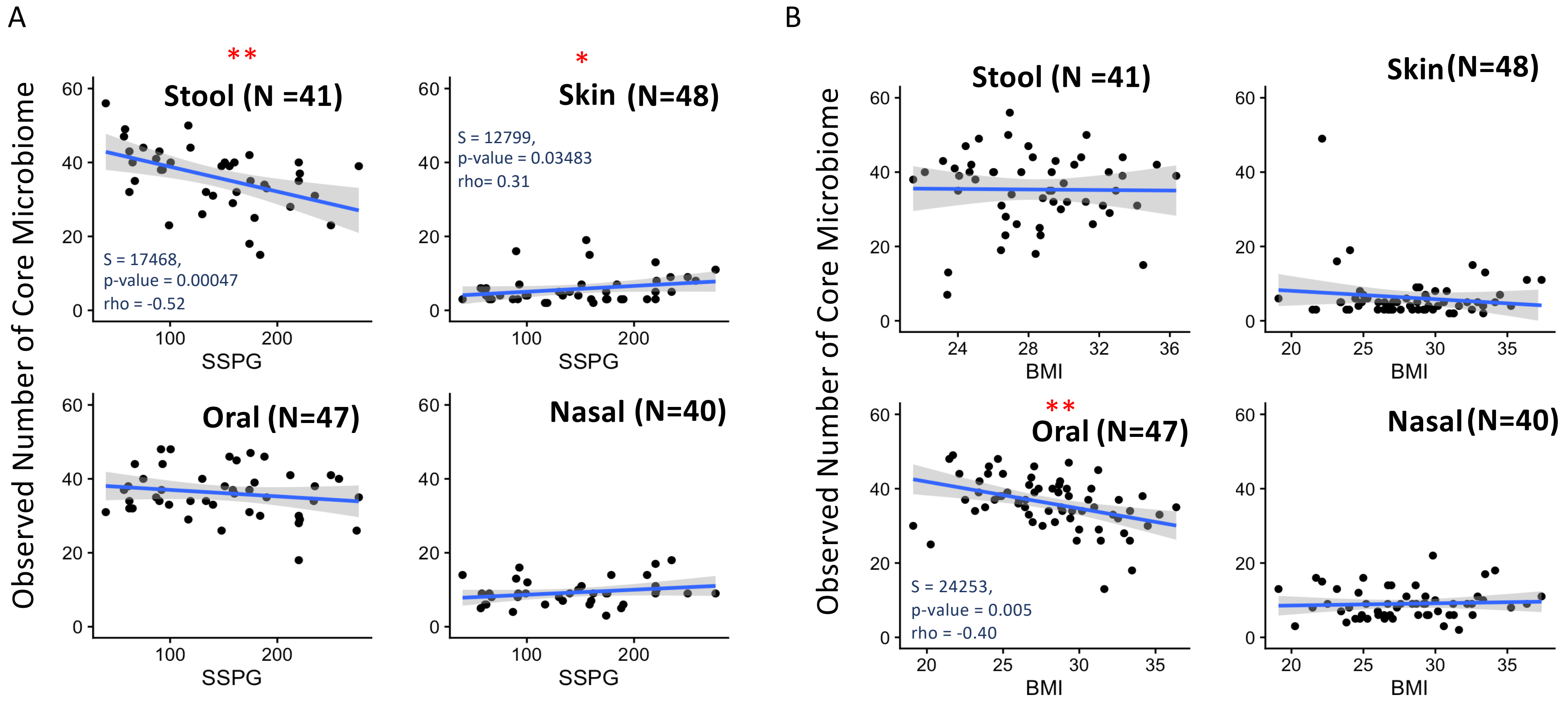

**Supplementary Figure S9. Number of Core Microbiome Genera in Insulin Sensitive and Insulin Resistant Individuals**

A

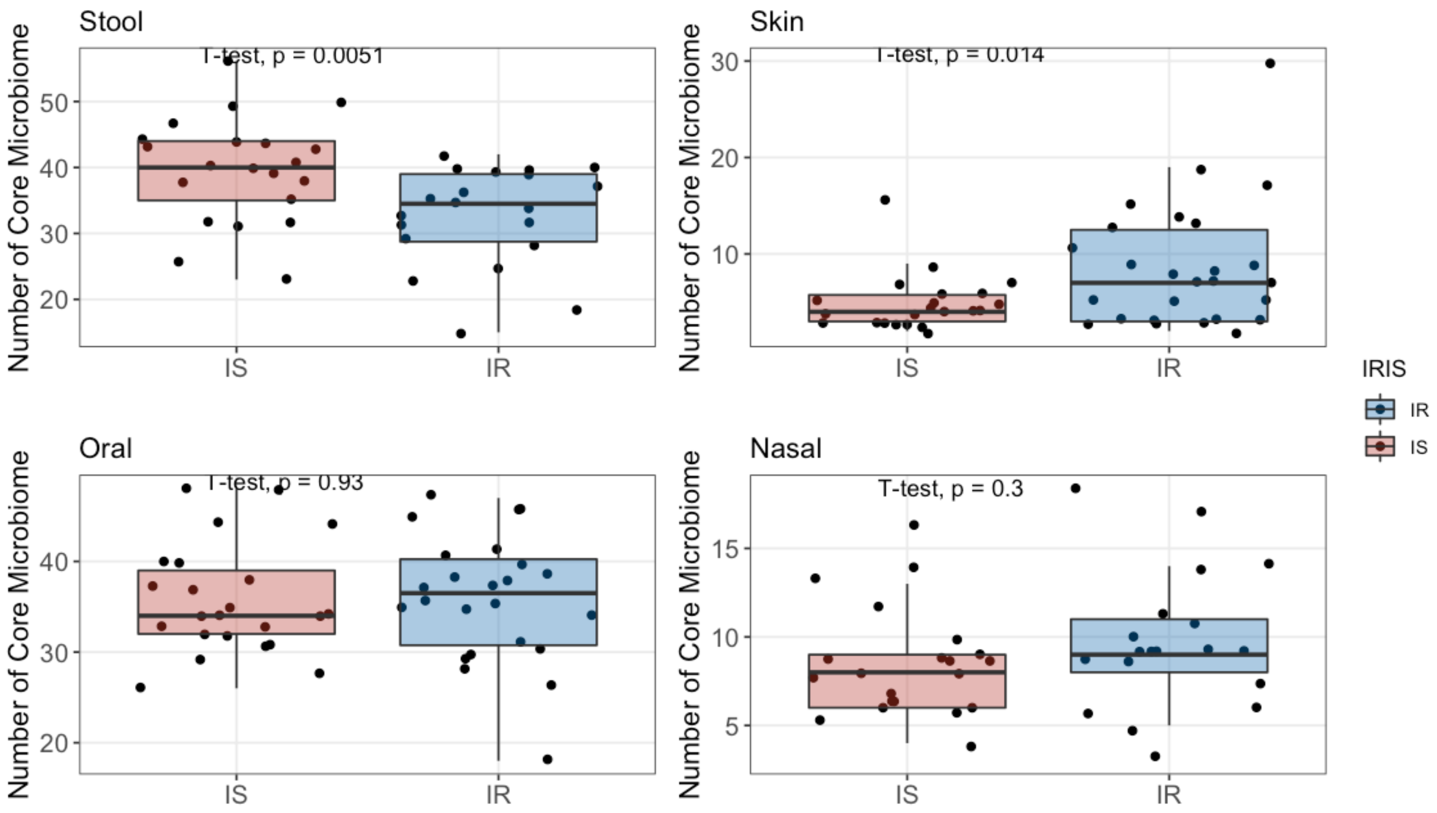

B

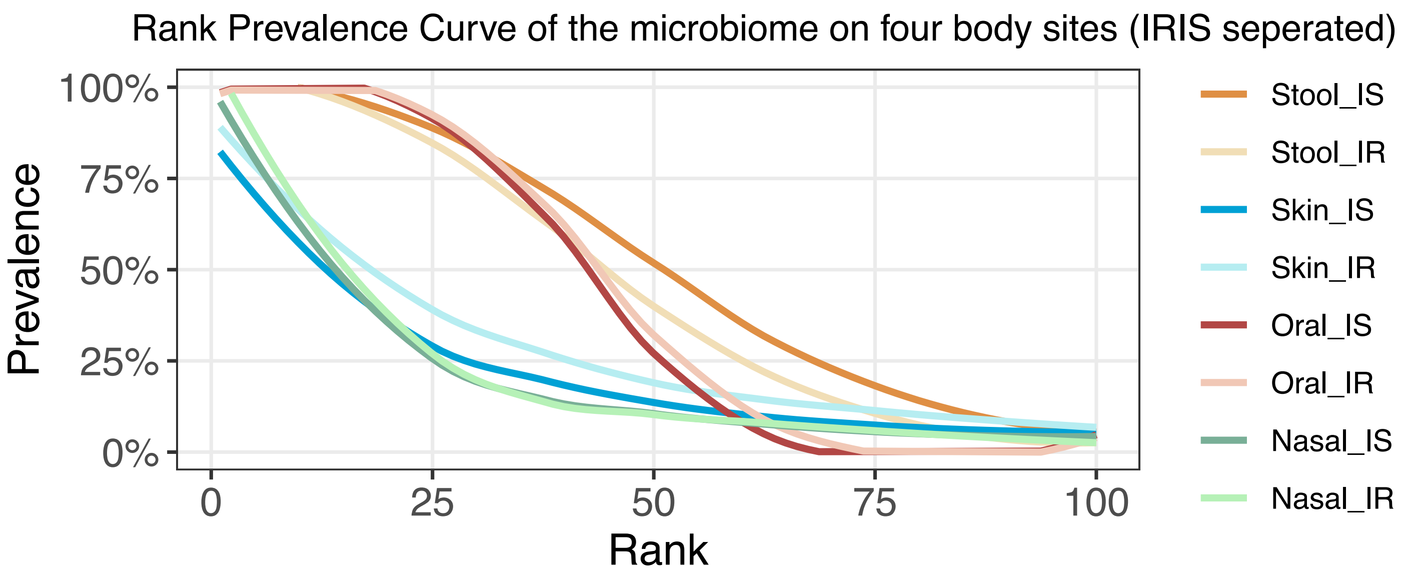
Rank Prevalence Curve of the microbiome on four body sites (IRIS seperated)
Prevalence
100%
75%
50%
25%
0%
0
25
50
75
100
Rank
Stool_IS
Stool_IR
Skin_IS
Skin_IR
Oral_IS
Oral_IR
Nasal_IS
Nasal_IR

#### Supplementary Figure S10: Effect Size of Taxa Differing in Relative Abundance Between Insulin Sensitive and Insulin Resistant Individuals

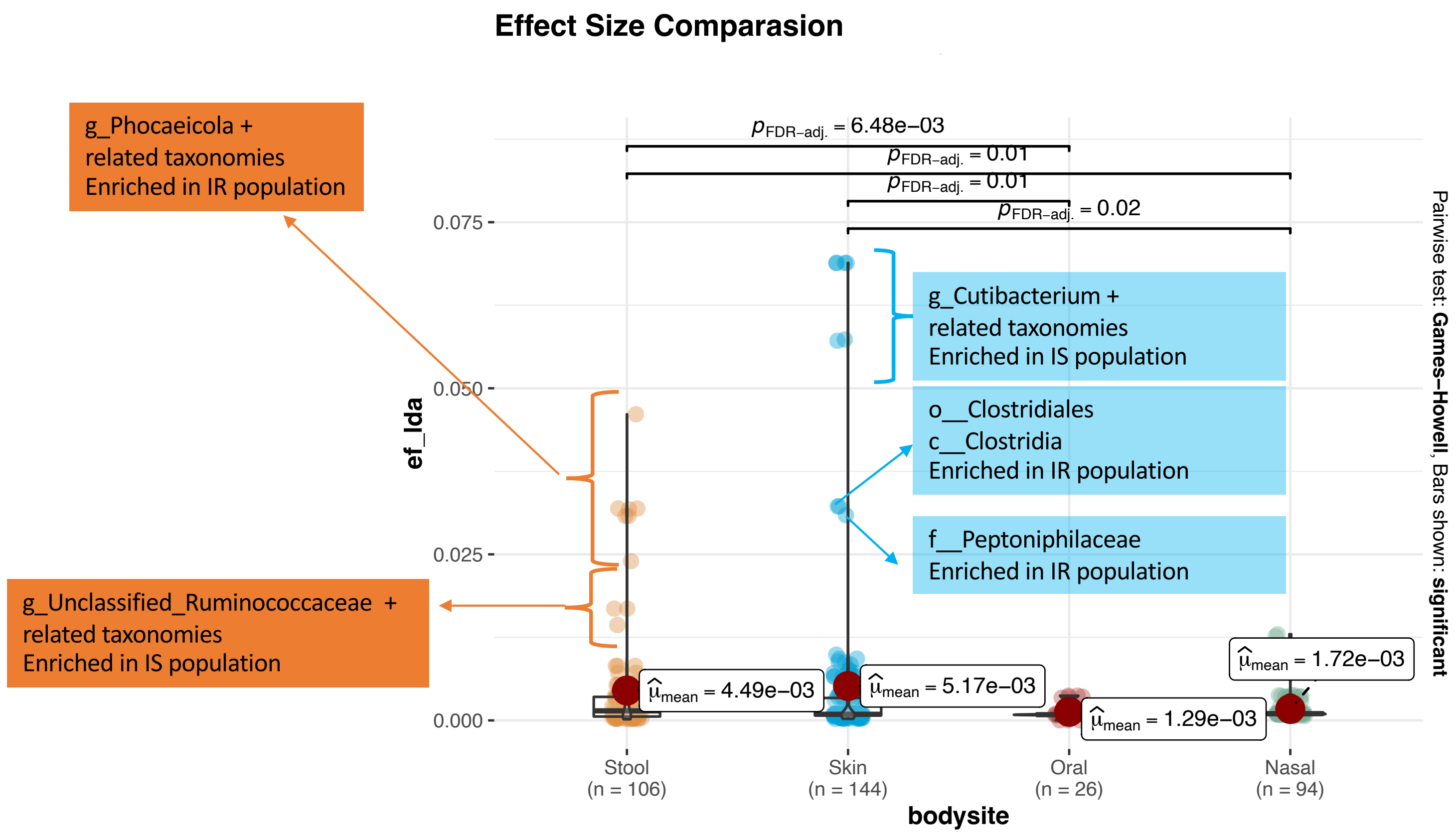

**Supplementary Figure S11. Histogram Distribution of Degree of Microbial Individuality and Family Score**

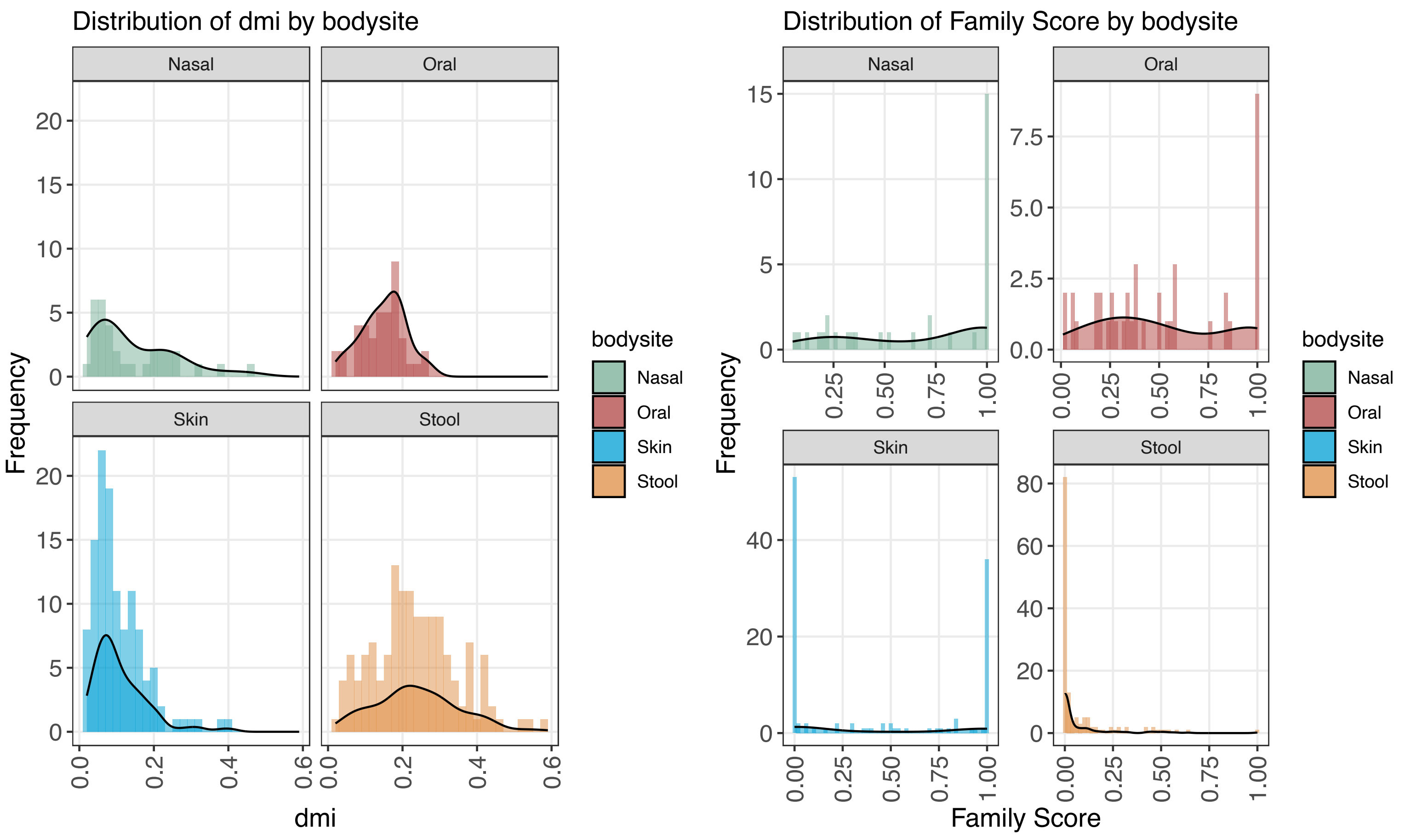

**Extended Data Figure S12. Comparative Analysis of Microbiome Individuality Across Different Insulin Sensitivity States**

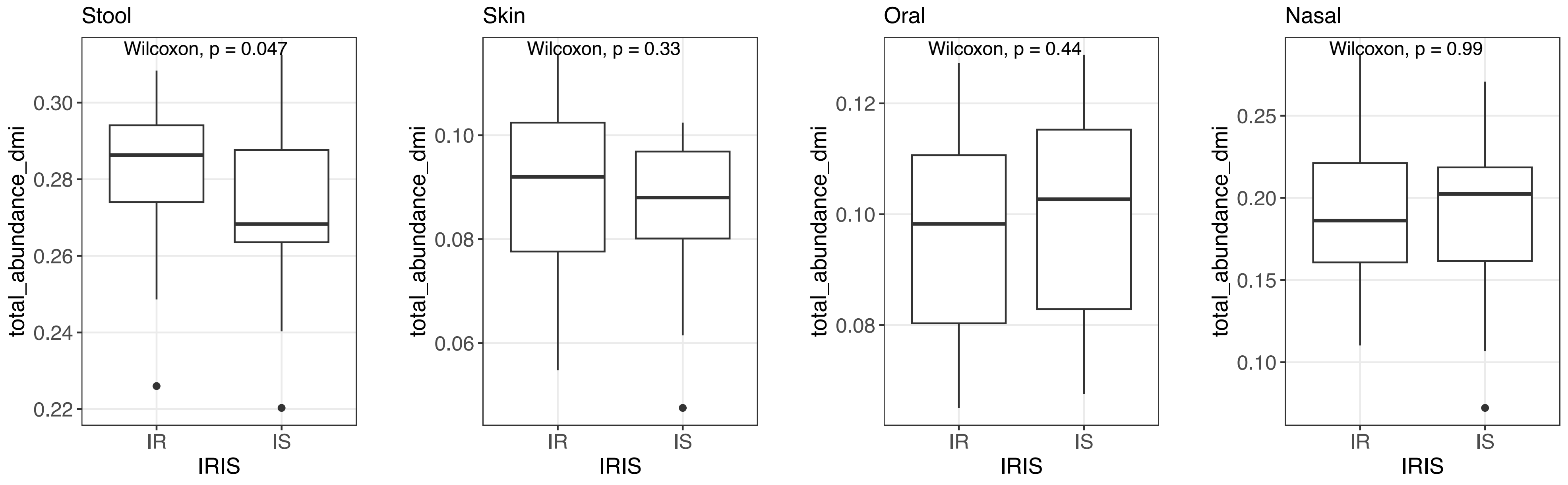

#### Supplementary Figure.S13 Strain replacement rate in insulin sensitive and insulin resistant individuals

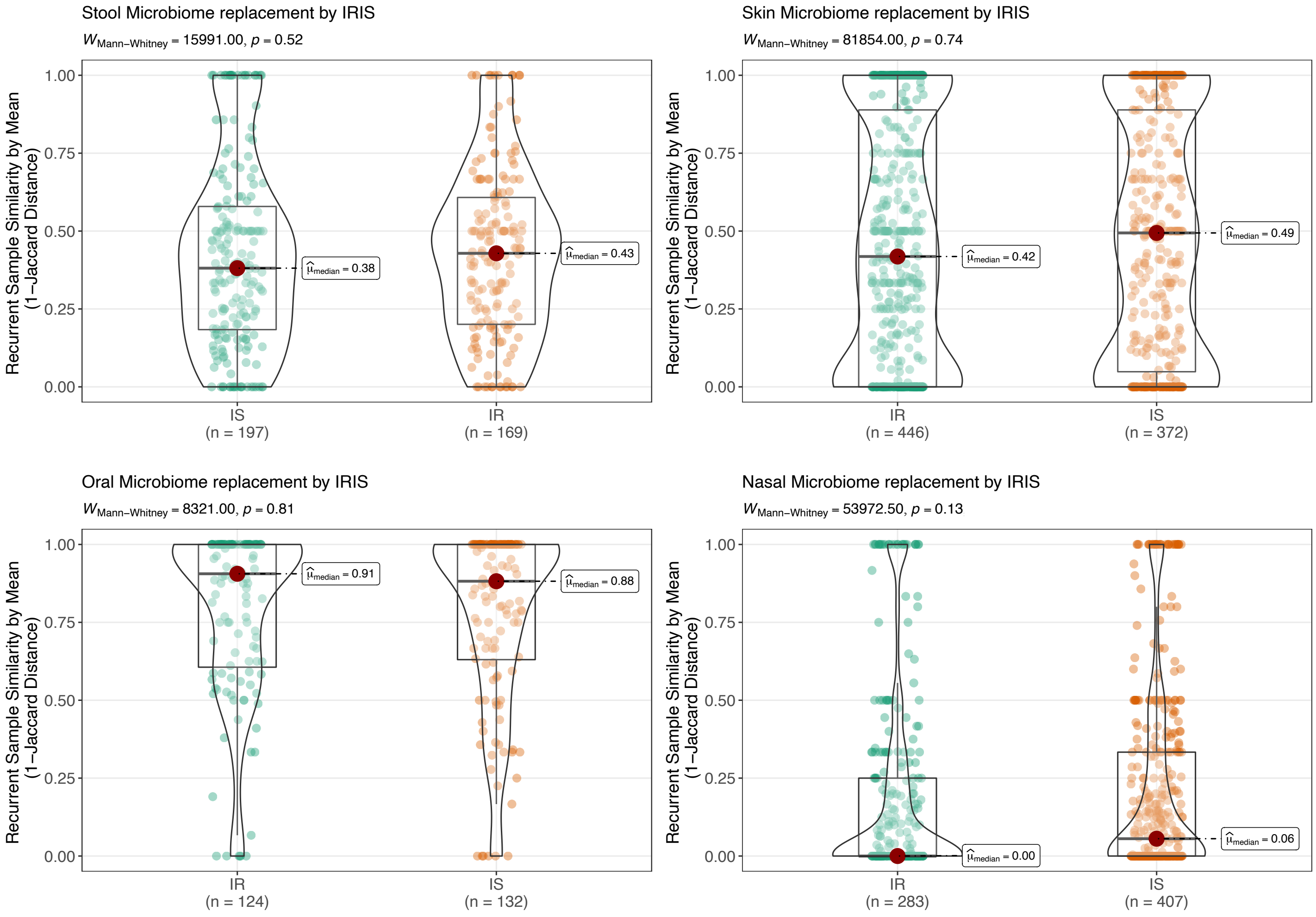

### Supplementary Figure. S14 Time related stability correlation between body sites in IS and IR group

A

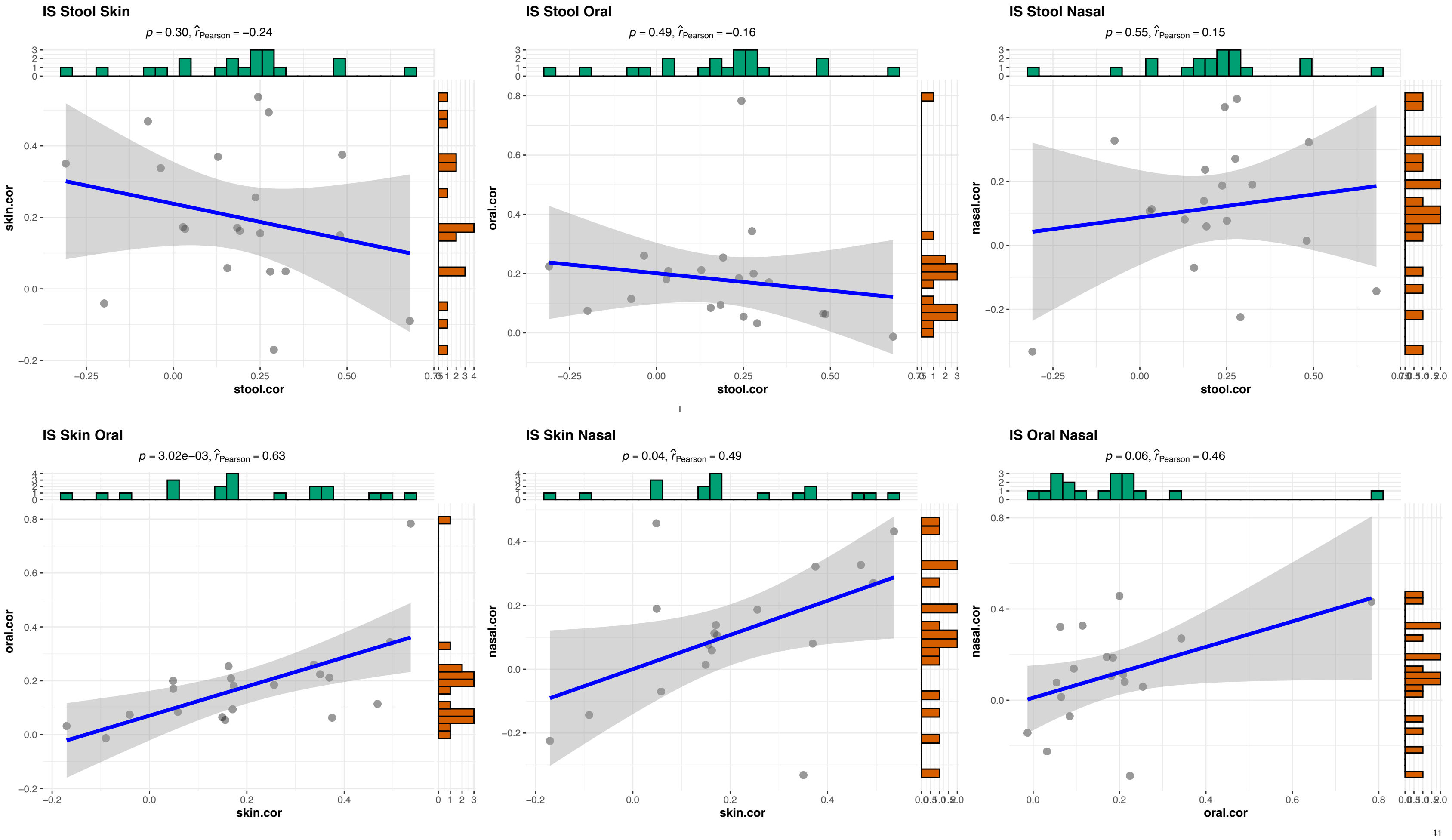

B

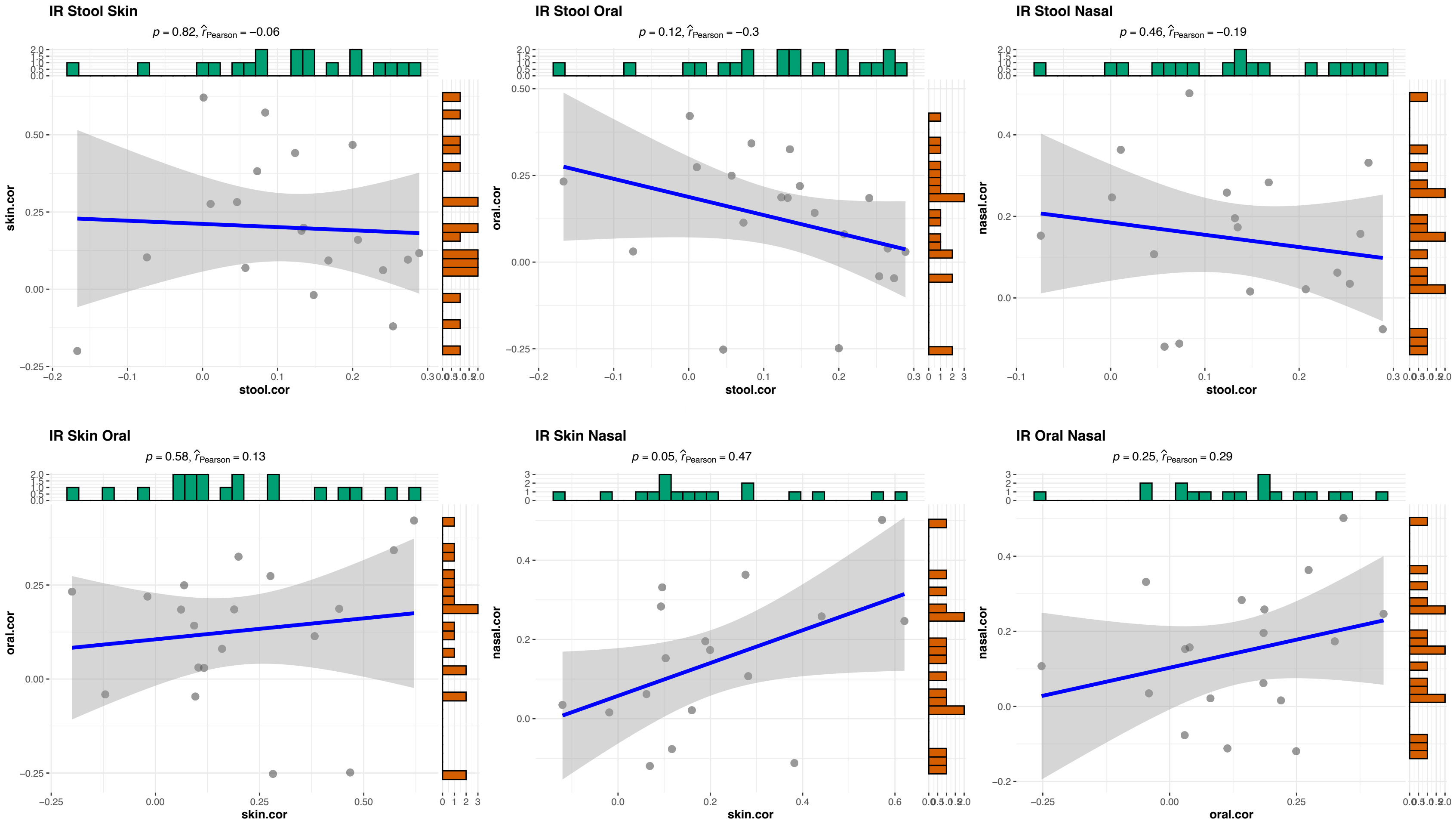

IR Stool Skin
p = 0.82, r̂Pearson = −0.06
skin.cor
stool.cor
IR Stool Oral
p = 0.12, r̂Pearson = −0.3
oral.cor
stool.cor
IR Stool Nasal
p = 0.46, r̂Pearson = −0.19
nasal.cor
stool.cor
IR Skin Oral
p = 0.58, r̂Pearson = 0.13
oral.cor
skin.cor
IR Skin Nasal
p = 0.05, r̂Pearson = 0.47
nasal.cor
skin.cor
IR Oral Nasal
p = 0.25, r̂Pearson = 0.29
nasal.cor
oral.cor

**Supplementary Figure S15. Degree of Microbial Individuality (DMI) comparison between bacteria genera correlated and uncorrelated between body sites**

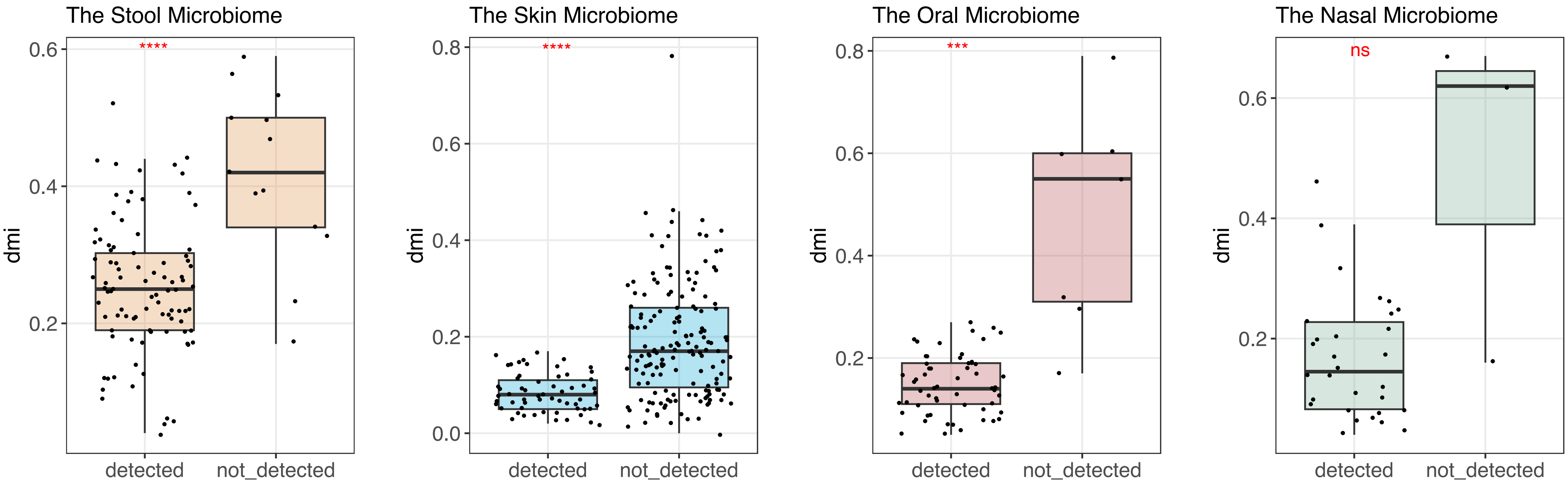

**Supplementary Figure S16: Microbial Evenness Change During Respiratory Viral Infection Among Insulin Sensitive and Insulin Resistant Individuals**

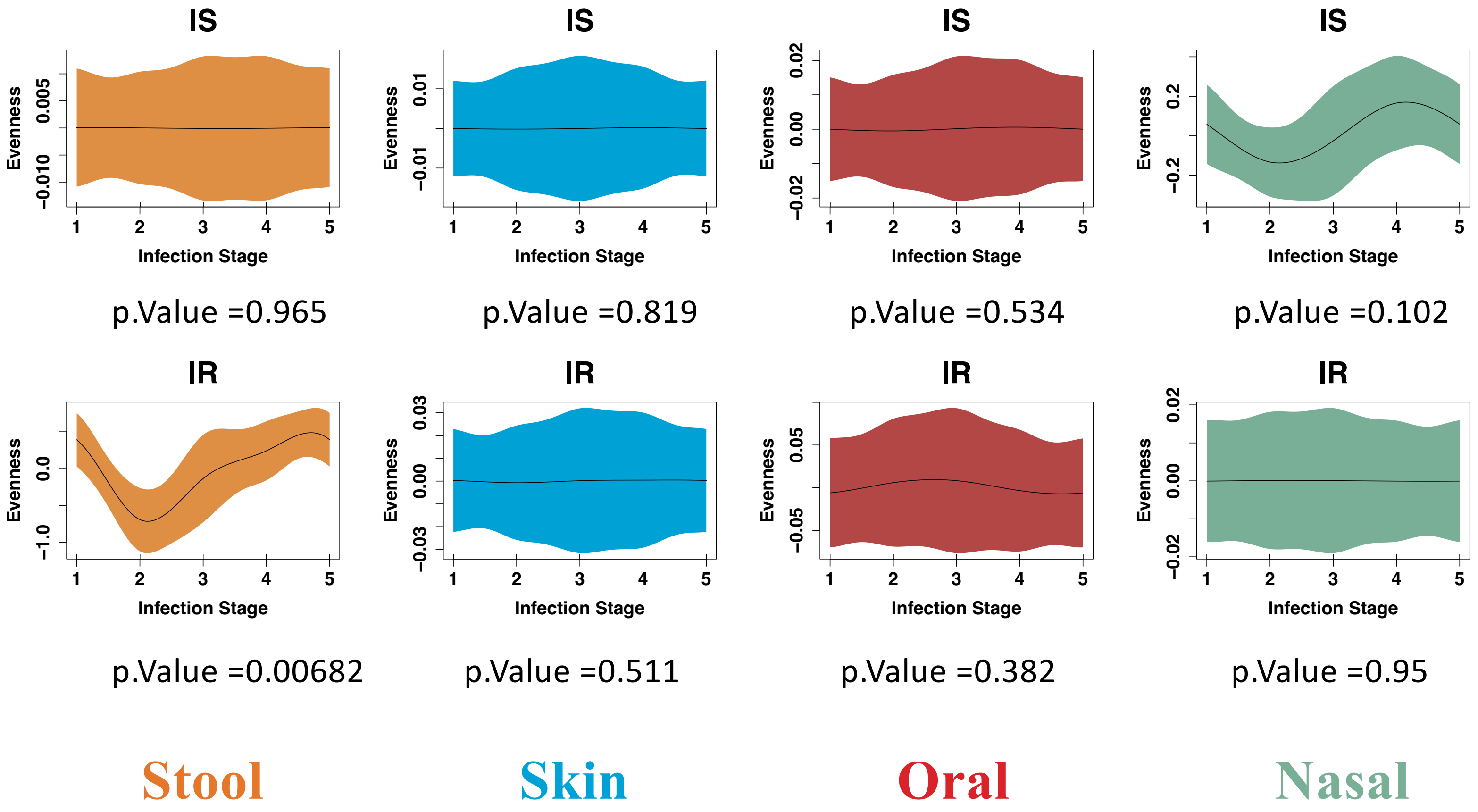

**Supplementary Figure S17: Microbial Relative Abundance Change During Respiratory Viral Infection**

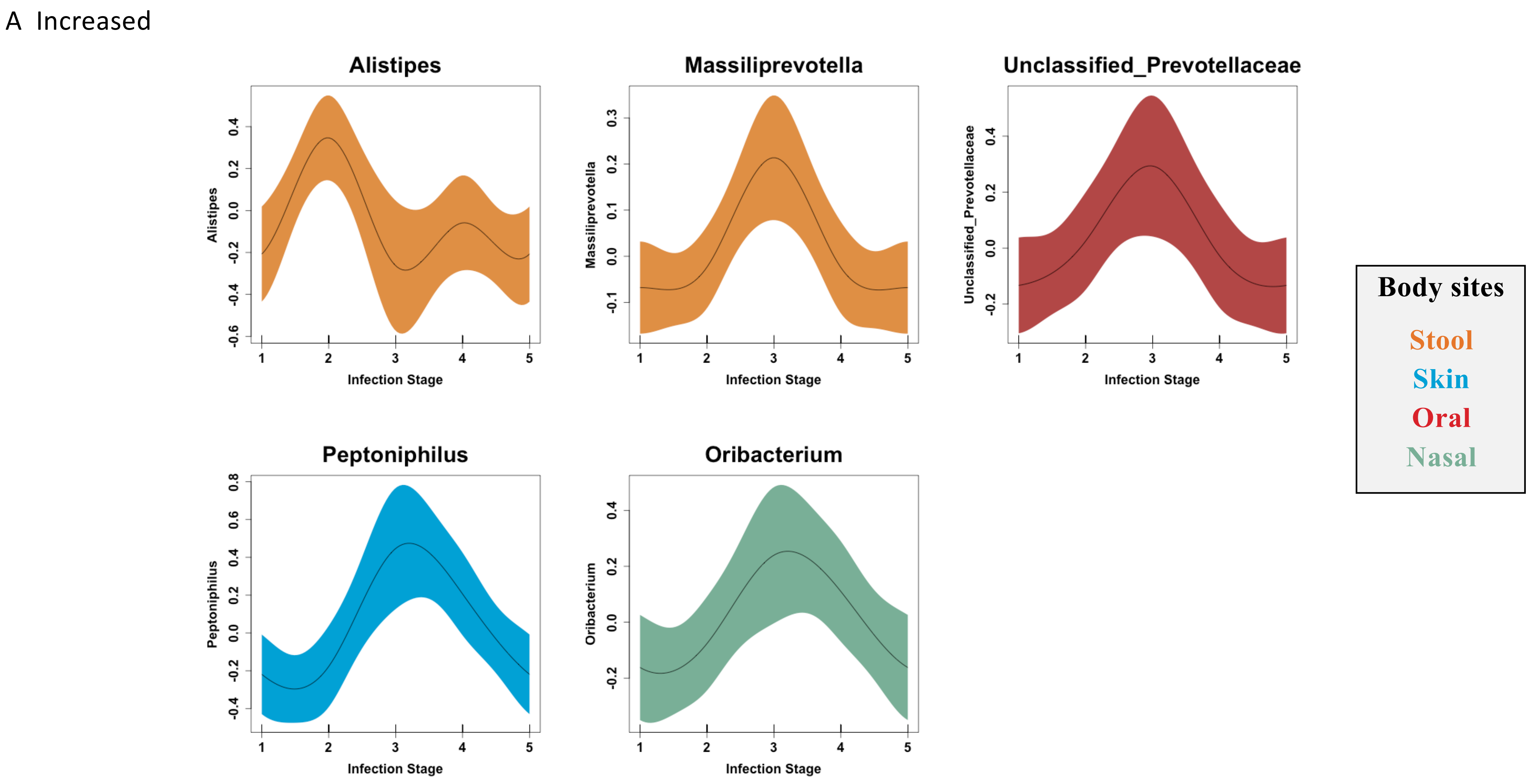

B Mixed

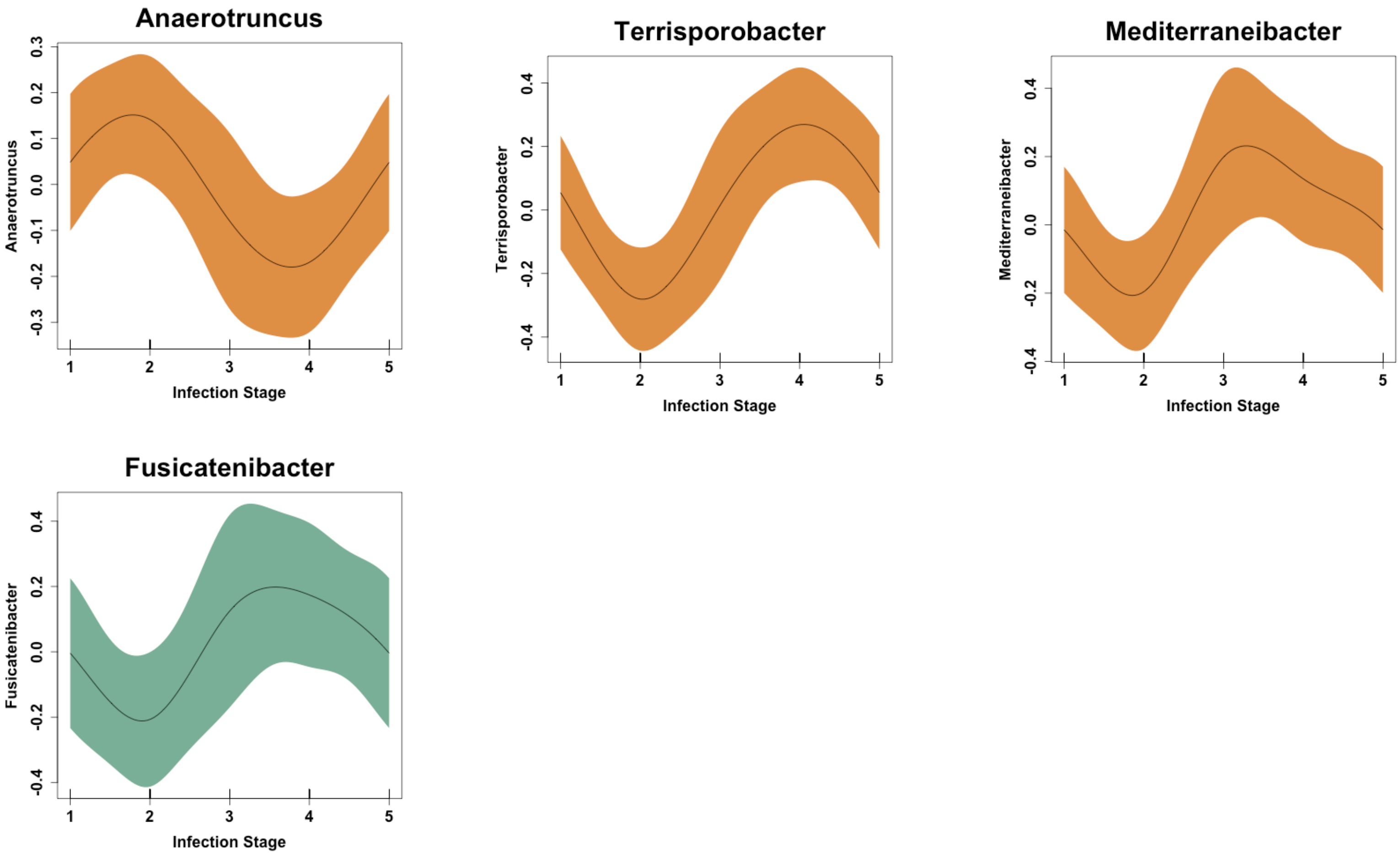
Anaerotruncus
Anaerotruncus
0.3
0.2
0.1
0.0
-0.1
-0.2
-0.3
1
2
3
4
5
Infection Stage
Terrisporobacter
Terrisporobacter
0.4
0.2
0.0
-0.2
-0.4
1
2
3
4
5
Infection Stage
Mediterraneibacter
Mediterraneibacter
0.4
0.2
0.0
-0.2
-0.4
1
2
3
4
5
Infection Stage
Fusicatenibacter
Fusicatenibacter
0.4
0.2
0.0
-0.2
-0.4
1
2
3
4
5
Infection Stage

C Decreased

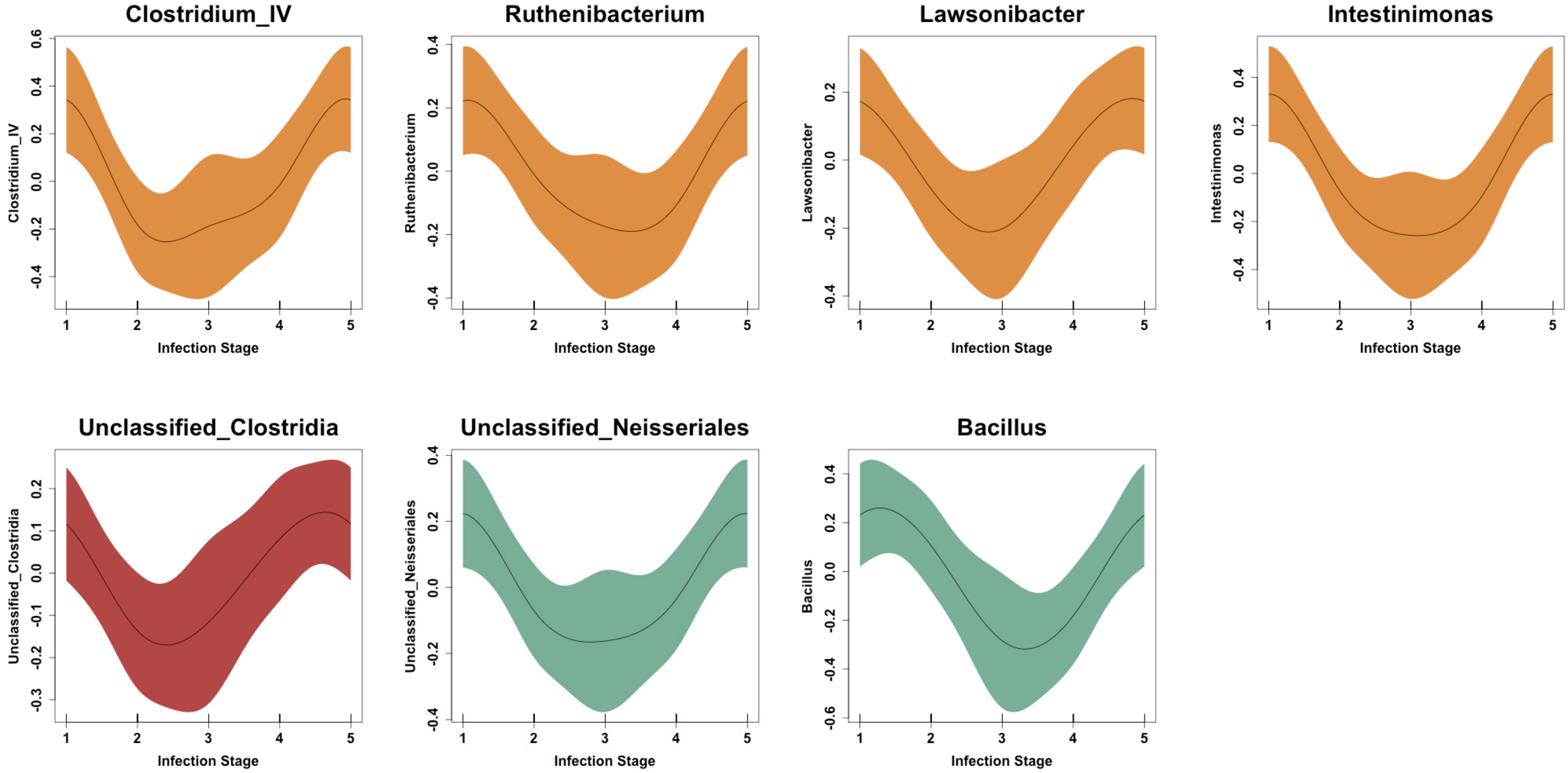
Clostridium_IV
Ruthenibacterium
Lawsonibacter
Intestinimonas
Unclassified_Clostridia
Unclassified_Neisseriales
Bacillus
Infection Stage

**Supplementary Figure Figure S18: Relationship Between the Microbiome and Cytokine Based on Their Correlation Coefficient**

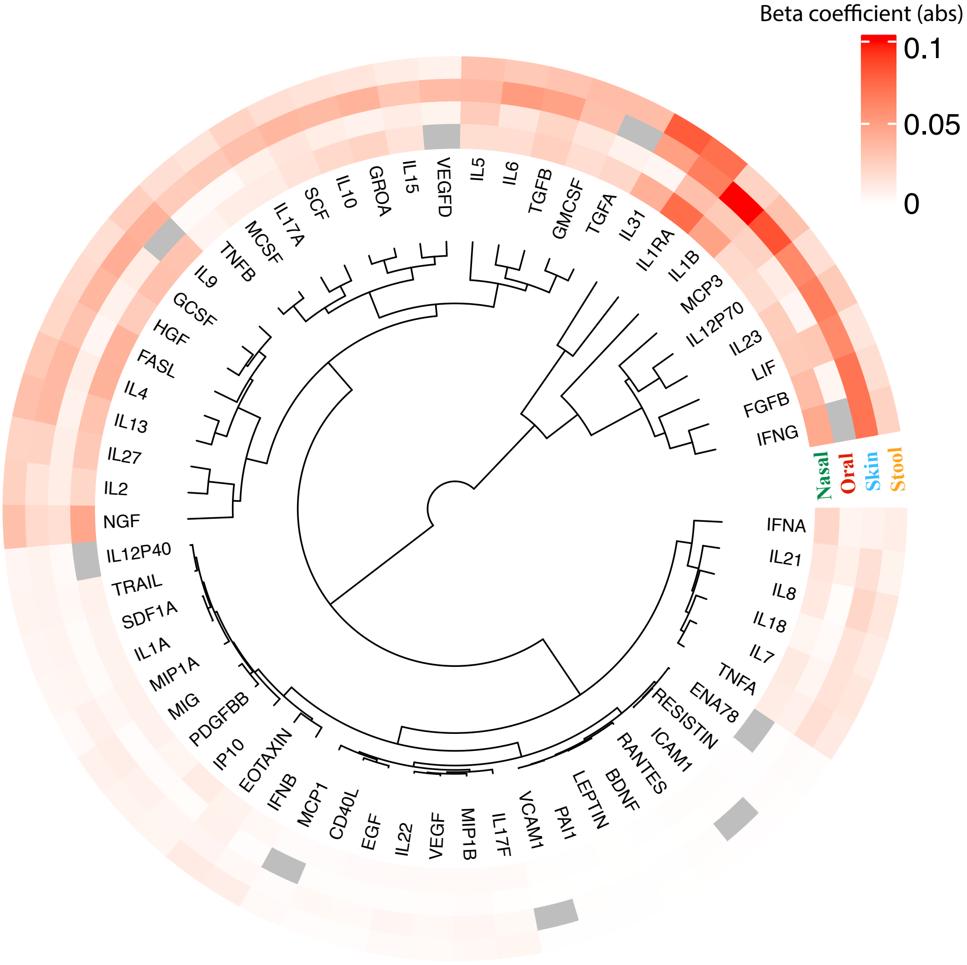

### Supplementary Figure Figure S19: Phyla Composition of Core, Middle, and Opportunistic Genera of Microbiome

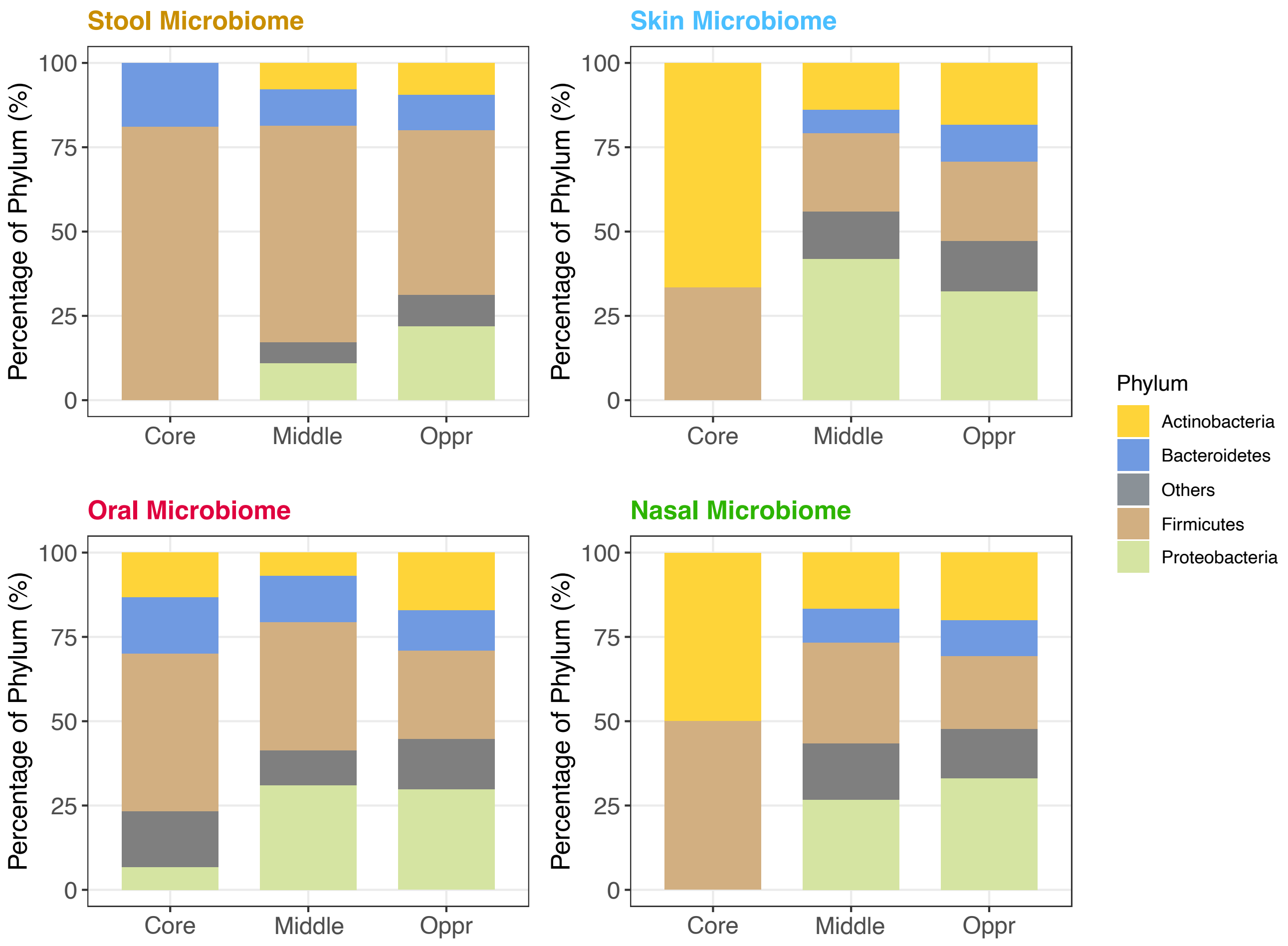

**Supplementary Figure S20: Correlation between Genera of Stool Proteobacteria and Plasma Cytokines Divided by Prevalence**

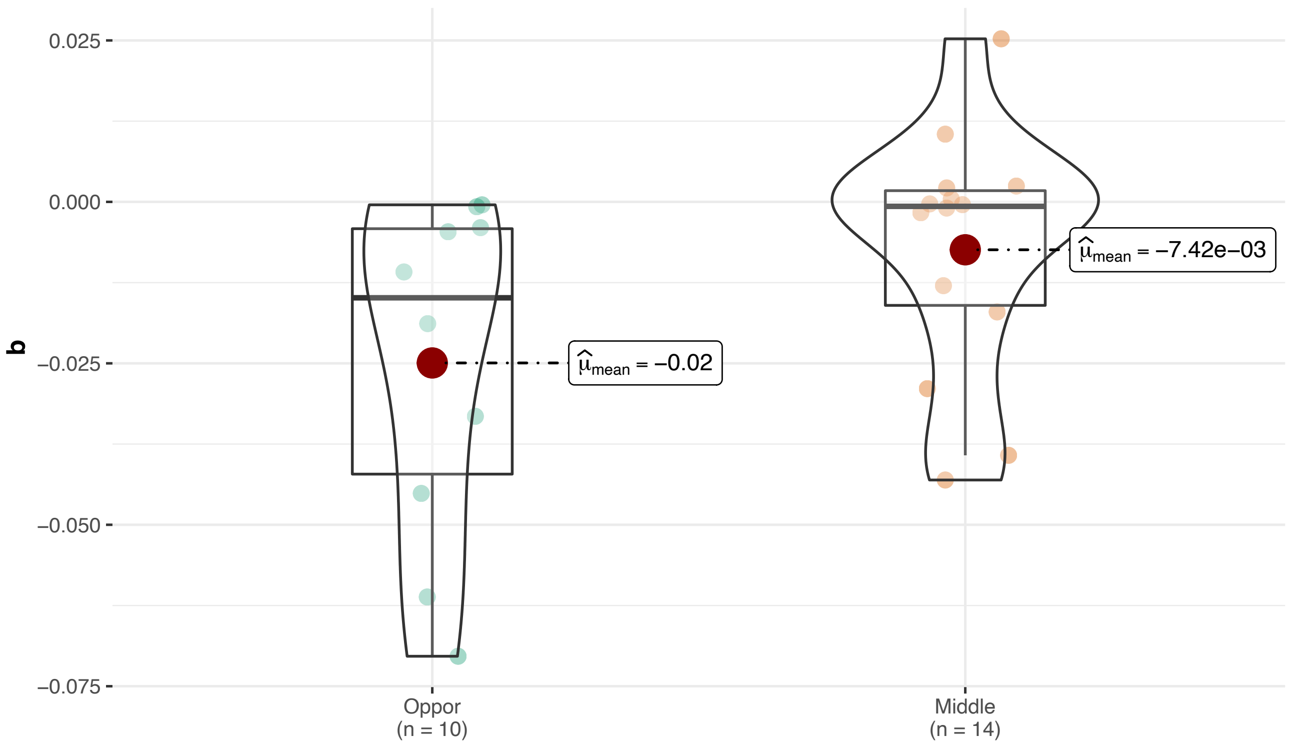

**Supplementary Figure S21: The Correlation between Body Mass Index and Plasma Leptin and Granulocyte-Macrophage Colony-Stimulating Factor**

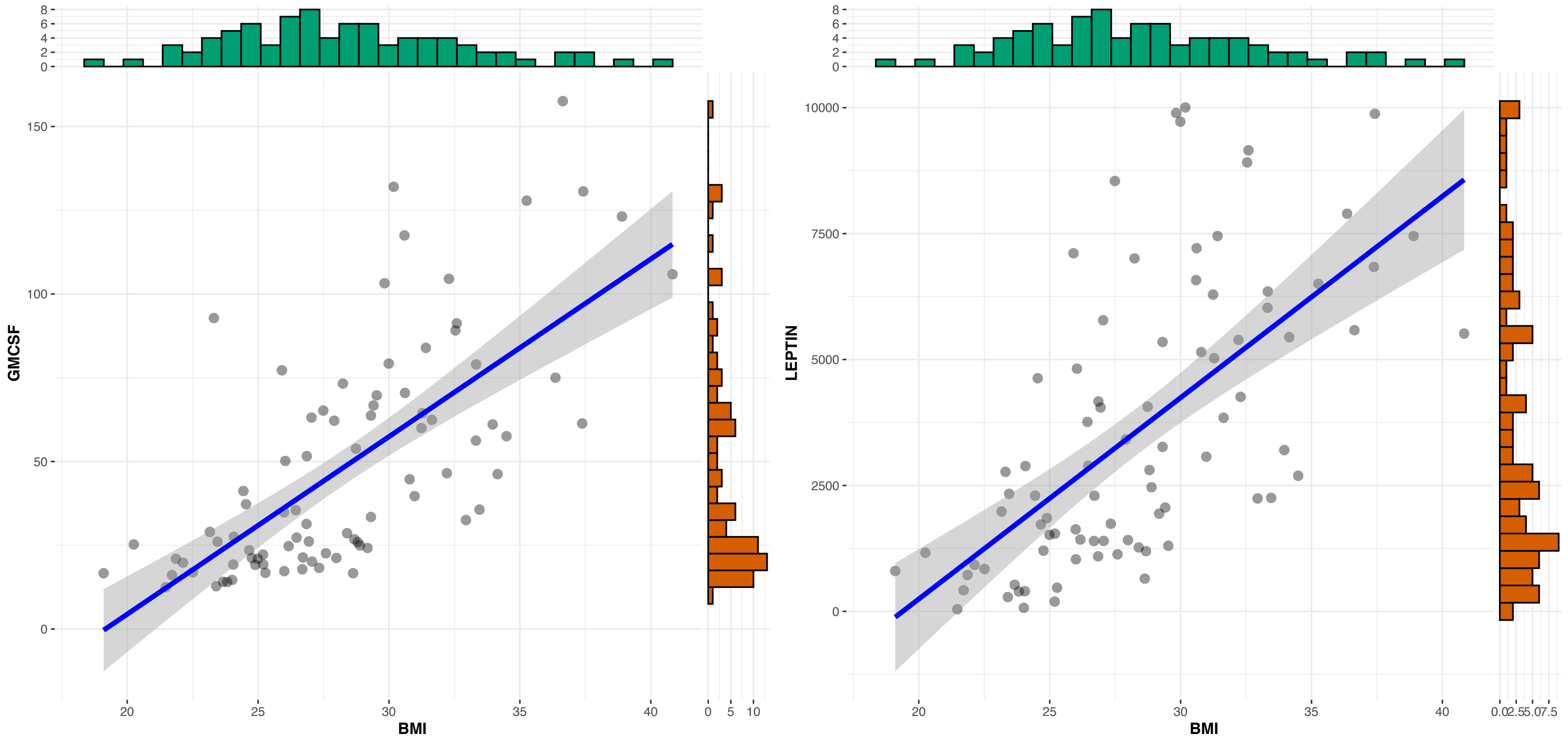

### Supplementary Figure S22: Collinearity of Metabolome, Lipidome, and Proteome

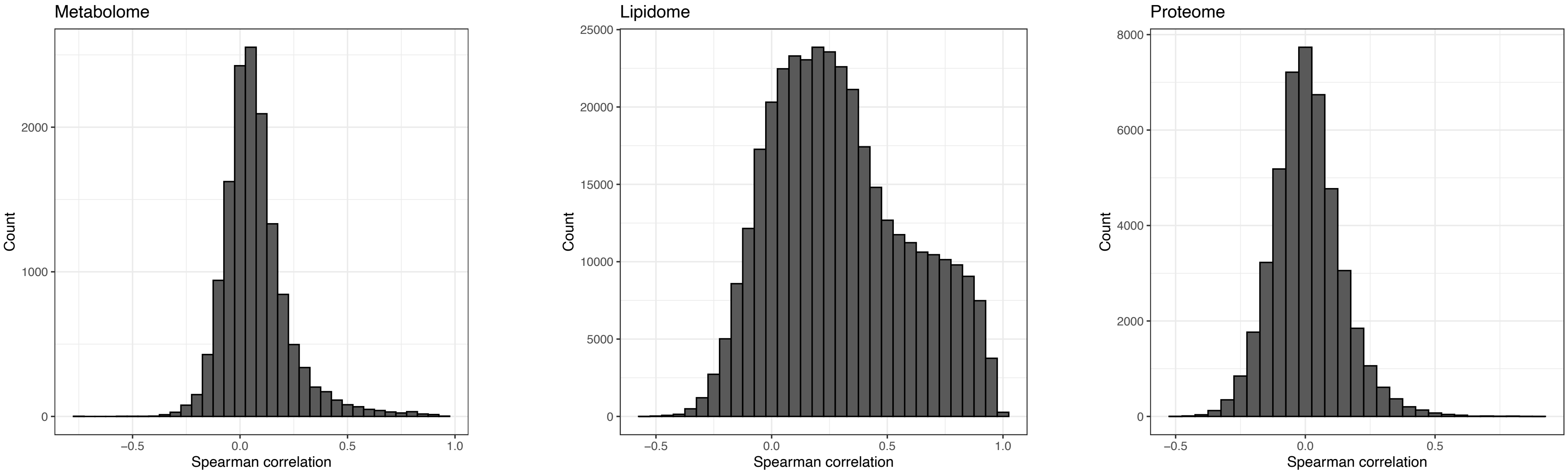

**Supplementary Figure S23: Pathway Enrichment Analysis of Proteomics and Microbiome Interactions**

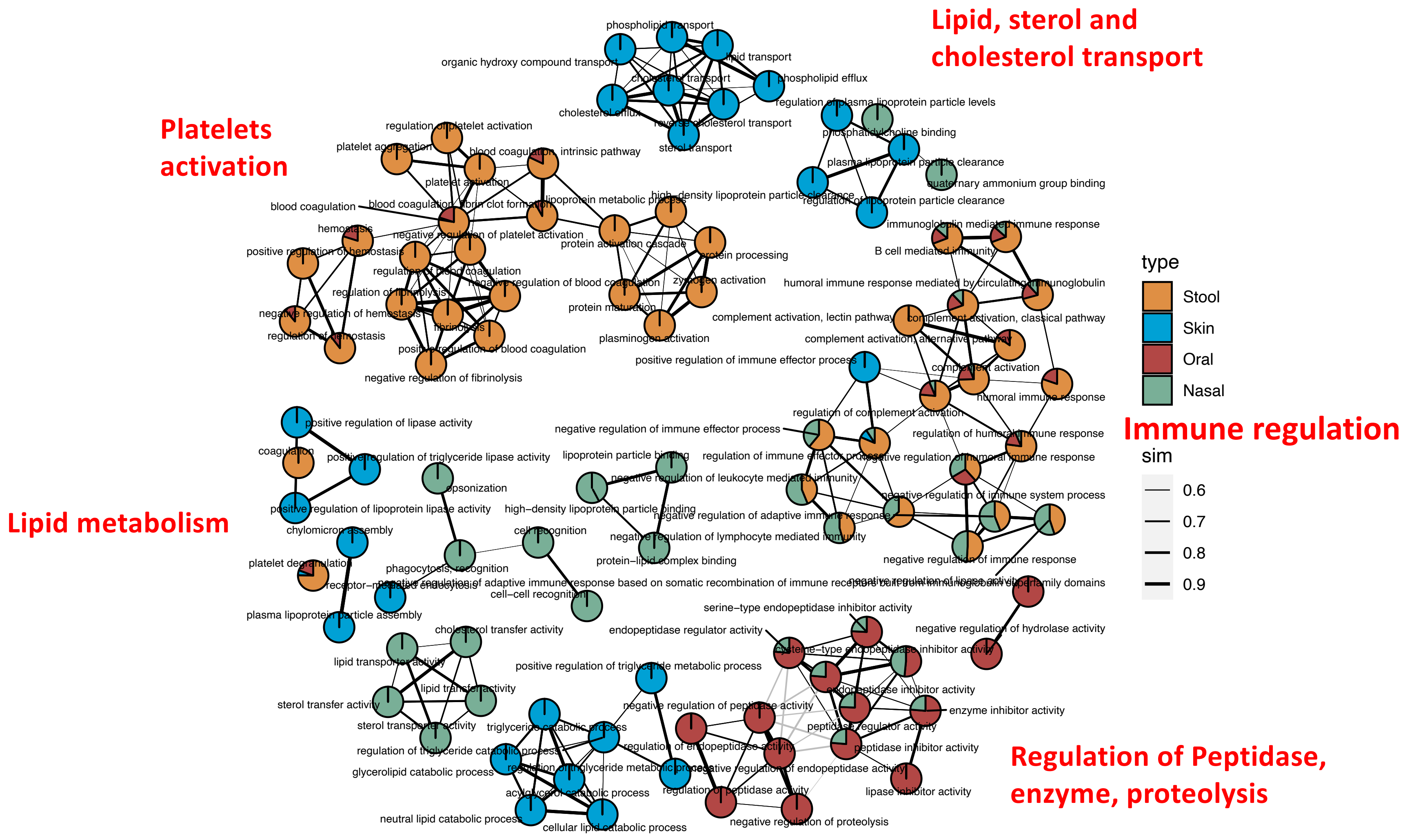

**Supplementary Figure S24: Different Interactome of the Stool Microbiome and Internal Plasma Analytics**

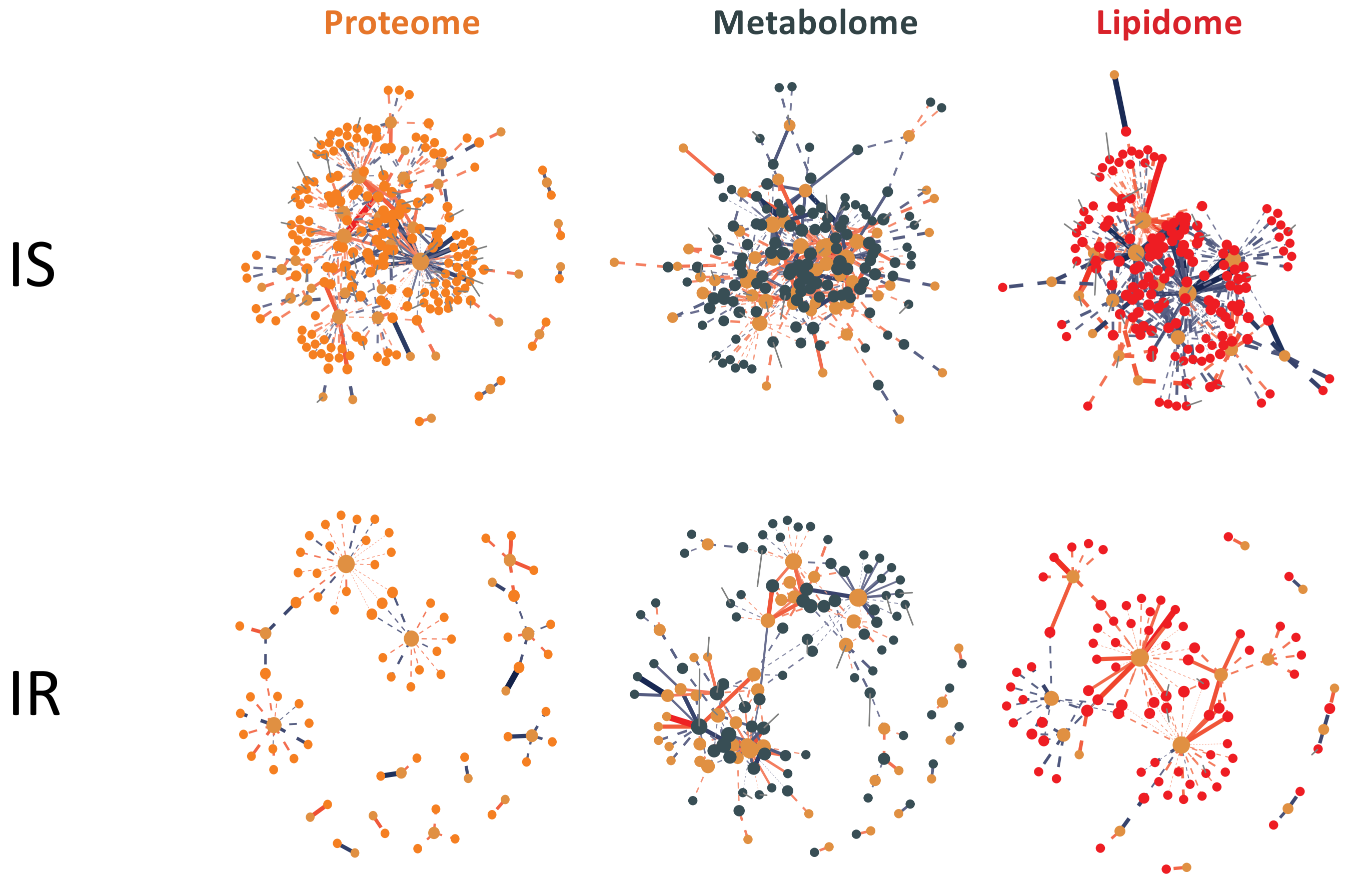


Node: t = −9.35, p = 1.18 x $10^{-10}$
Edge: t = −7.93, p = 2.27 x $10^{-08}$

Node: t = −6.12, p = 3.85 x$10^{-07}$
Edge: t = −4.02, p = 3.45 x $10^{-04}$

Node: t = −5.99, p = 6.98 x $10^{-07}$
Edge: t = −5.32, p = 5.76 x $10^{-06}$

#### Supplementary Figure S25: Correlation Network of Microbiome Richness and Multiple Internal Molecular Markers on Four Body Sites

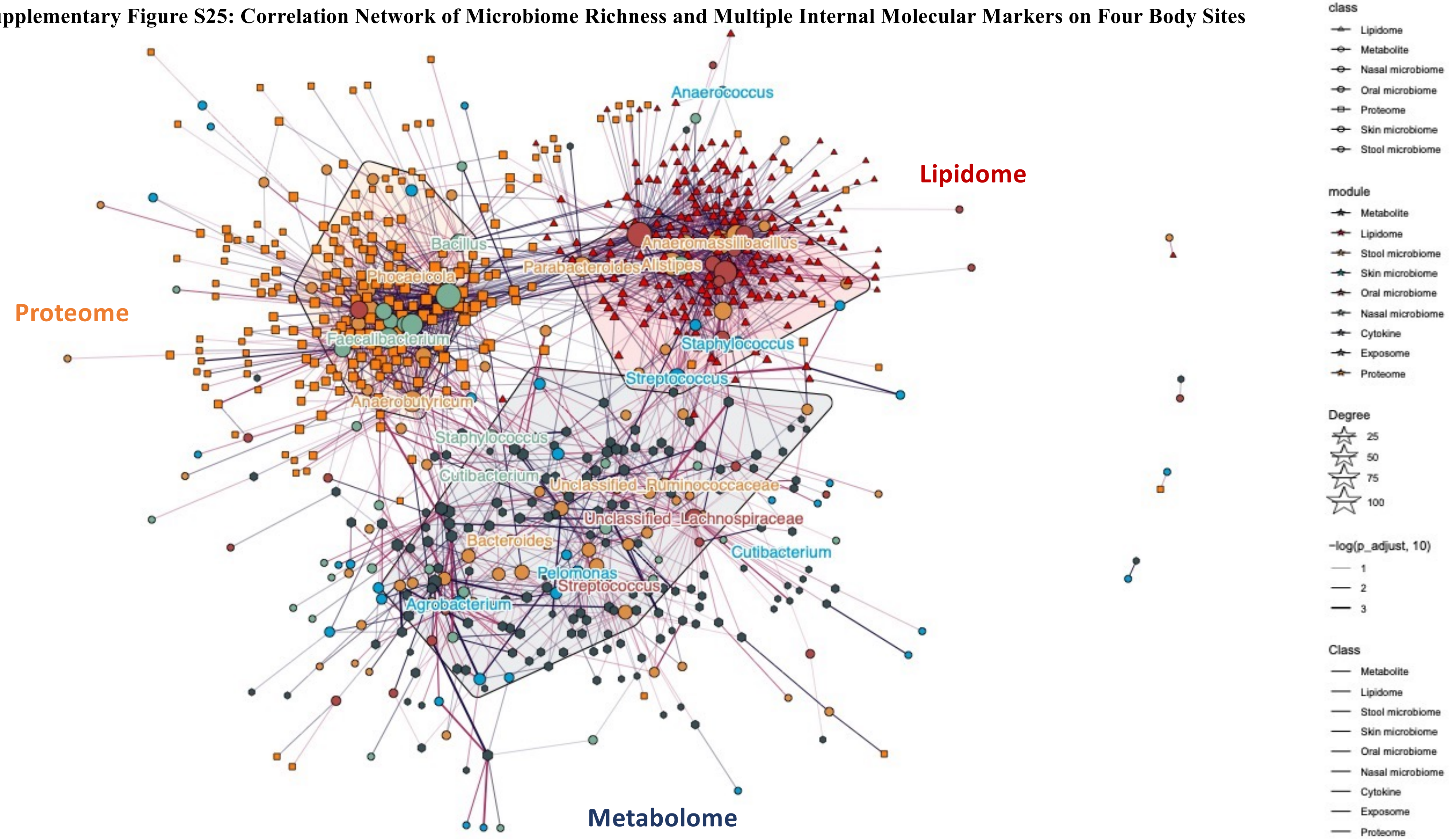

**Supplementary Figure S26: Distribution of Correlation Coefficients for Microbiome Interactions Across Four Body Sites**

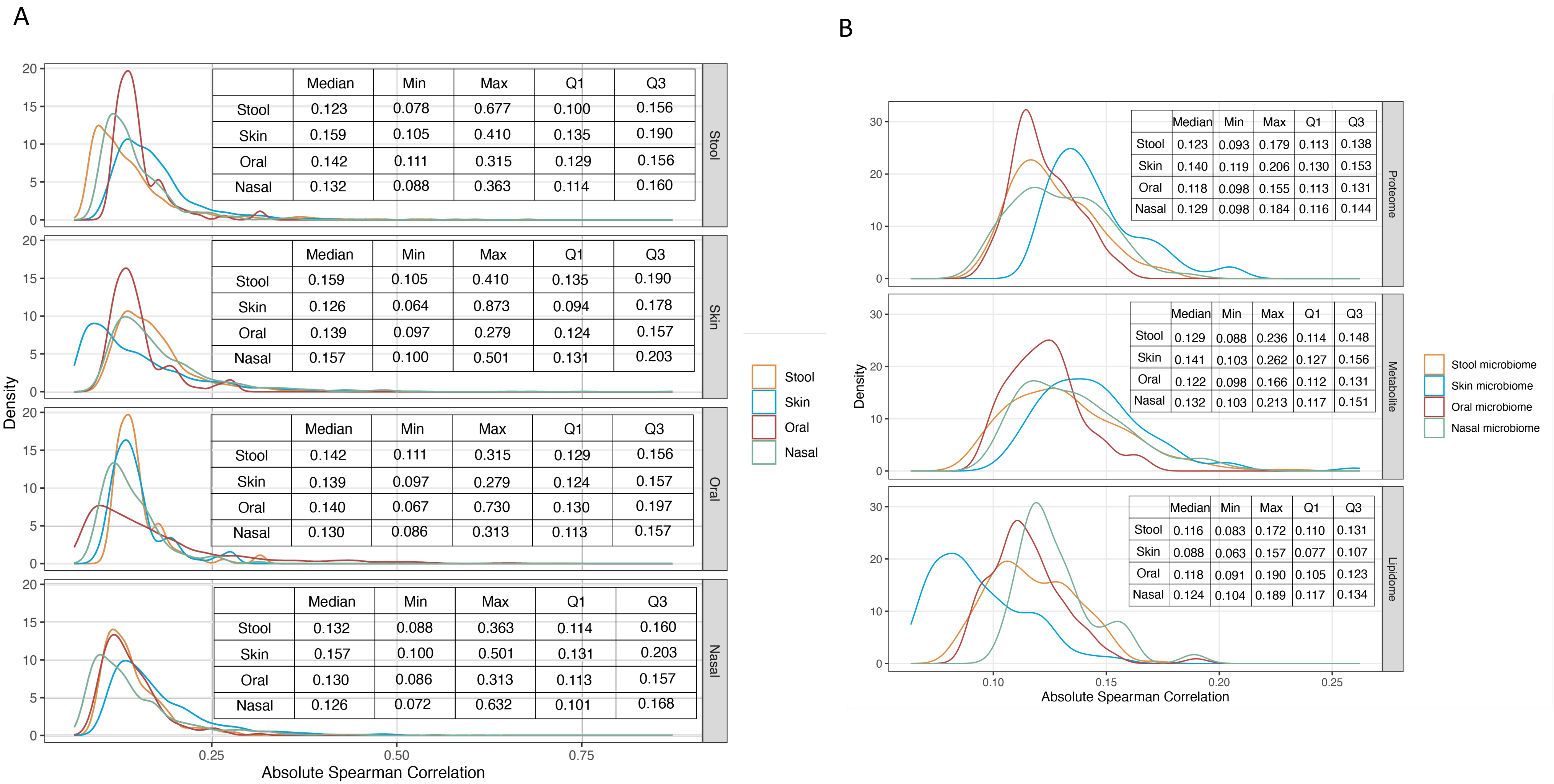


| | Median | Min | Max | Q1 | Q3 |
|---|---|---|---|---|---|
| Stool | 0.123 | 0.078 | 0.677 | 0.100 | 0.156 |
| Skin | 0.159 | 0.105 | 0.410 | 0.135 | 0.190 |
| Oral | 0.142 | 0.111 | 0.315 | 0.129 | 0.156 |
| Nasal | 0.132 | 0.088 | 0.363 | 0.114 | 0.160 |

| | Median | Min | Max | Q1 | Q3 |
|---|---|---|---|---|---|
| Stool | 0.159 | 0.105 | 0.410 | 0.135 | 0.190 |
| Skin | 0.126 | 0.064 | 0.873 | 0.094 | 0.178 |
| Oral | 0.139 | 0.097 | 0.279 | 0.124 | 0.157 |
| Nasal | 0.157 | 0.100 | 0.501 | 0.131 | 0.203 |

| | Median | Min | Max | Q1 | Q3 |
|---|---|---|---|---|---|
| Stool | 0.142 | 0.111 | 0.315 | 0.129 | 0.156 |
| Skin | 0.139 | 0.097 | 0.279 | 0.124 | 0.157 |
| Oral | 0.140 | 0.067 | 0.730 | 0.130 | 0.197 |
| Nasal | 0.130 | 0.086 | 0.313 | 0.113 | 0.157 |

| | Median | Min | Max | Q1 | Q3 |
|---|---|---|---|---|---|
| Stool | 0.132 | 0.088 | 0.363 | 0.114 | 0.160 |
| Skin | 0.157 | 0.100 | 0.501 | 0.131 | 0.203 |
| Oral | 0.130 | 0.086 | 0.313 | 0.113 | 0.157 |
| Nasal | 0.126 | 0.072 | 0.632 | 0.101 | 0.168 |

| | Median | Min | Max | Q1 | Q3 |
|---|---|---|---|---|---|
| Stool | 0.123 | 0.093 | 0.179 | 0.113 | 0.138 |
| Skin | 0.140 | 0.119 | 0.206 | 0.130 | 0.153 |
| Oral | 0.118 | 0.098 | 0.155 | 0.113 | 0.131 |
| Nasal | 0.129 | 0.098 | 0.184 | 0.116 | 0.144 |

| | Median | Min | Max | Q1 | Q3 |
|---|---|---|---|---|---|
| Stool | 0.129 | 0.088 | 0.236 | 0.114 | 0.148 |
| Skin | 0.141 | 0.103 | 0.262 | 0.127 | 0.156 |
| Oral | 0.122 | 0.098 | 0.166 | 0.112 | 0.131 |
| Nasal | 0.132 | 0.103 | 0.213 | 0.117 | 0.151 |

| | Median | Min | Max | Q1 | Q3 |
|---|---|---|---|---|---|
| Stool | 0.116 | 0.083 | 0.172 | 0.110 | 0.131 |
| Skin | 0.088 | 0.063 | 0.157 | 0.077 | 0.107 |
| Oral | 0.118 | 0.091 | 0.190 | 0.105 | 0.123 |
| Nasal | 0.124 | 0.104 | 0.189 | 0.117 | 0.134 |
